# Modeling the growth of cytosolic TDP-43 inclusion bodies and accumulated neurotoxicity of misfolded oligomers in neurons

**DOI:** 10.1101/2023.11.28.569118

**Authors:** Andrey V. Kuznetsov

## Abstract

This paper introduces a mathematical model for the growth of transactive response DNA binding protein of 43 kDa (TDP-43) inclusion bodies in neuron soma. The parameter representing the accumulated neurotoxicity caused by misfolded TDP-43 oligomers is also introduced. The model’s equations enable the numerical calculation of the concentrations of TDP-43 monomers, dimers, free oligomers, and oligomers deposited in inclusion bodies. By simulating the deposition of free oligomers into inclusion bodies, the model predicts the size of TDP-43 inclusion bodies. An approximate solution to the model equations is derived for the scenario where protein degradation machinery is dysfunctional, leading to infinite half-lives for TDP-43 dimers, monomers, and both free and deposited oligomers. This solution, valid at large times, predicts that the radius of the inclusion body increases proportionally to the cube root of time, whereas the accumulated neurotoxicity increases linearly with time. To the best of the author’s knowledge, this study is the first to model the relationship between the size of TDP-43 inclusion bodies and time, and the first to introduce the concept of accumulated neurotoxicity caused by misfolded TDP-43 oligomers. Sensitivity analysis of the approximate solution indicates that the inclusion body radius and accumulated neurotoxicity become independent of the kinetic constants at large timescales. While the numerical solution of the full mathematical model continues to work with finite half-lives, the approximate solution becomes invalid for scenarios with physiologically relevant (finite) half-lives of TDP-43 dimers, monomers, and free misfolded oligomers. In contrast to the scenario with infinite half-lives, the numerical solution reveals that, for different values of the kinetic constants, the curves representing the inclusion body radius and accumulated neurotoxicity converge to distinct curves over time.

## 1. Introduction

The presence of transactive response DNA binding protein of 43 kDa (TDP-43) inclusion bodies represents a pathological hallmark in amyotrophic lateral sclerosis (ALS) and frontotemporal lobar degeneration (FTLD) diseases (Ishii et al. 2017; Jo et al. 2020; Prasad et al. 2019; Capitini et al. 2014). ALS is distinguished by the gradual decline of motor neurons, resulting in skeletal muscle weakness and, ultimately, fatality within a span of 3 to 5 years (Oiwa et al. 2023; Sugai et al. 2018). FTLD is an early-onset neurodegenerative disease characterized by progressive deterioration in behavior and language due to the loss of neurons, primarily in the frontal or temporal lobes, with a life expectancy of 6-8 years (Grossman et al. 2023). TDP-43, a nuclear, intrinsically disordered, soluble protein (Sun et al. 2019; French et al. 2019) with a molecular weight of 43 kDa (Suk and Rousseaux 2020), is synthesized in the cytoplasm and, in healthy neurons, predominantly resides in the nucleus (Pérez-Berlanga et al. 2023; Necarsulmer et al. 2023; Meneses et al. 2021). TDP-43 is degraded through both the proteasomal and autophagy pathways (Chhangani et al. 2021). However, in TDP-43 proteinopathies, this protein undergoes mislocalization from the nucleus to the cytoplasm (Glass 2020; Suk and Rousseaux 2020), forming TDP-43 inclusion bodies characterized by ubiquitinated and hyper-phosphorylated variants of TDP-43 (Prasad et al. 2019).

Under pathological conditions, TDP-43 undergoes a conversion from its native dimeric state (Jiang et al. 2017) to a monomeric form. This transformation results in TDP-43 mislocalization from the nucleus to the cytoplasm, where it forms irreversible, amorphous aggregates (Sun et al. 2014; Oiwa et al. 2023). Subsequently, these aggregates grow into inclusion bodies. Notably, the nature of these inclusion bodies, whether they exhibit toxicity or act in a neuroprotective manner by sequestering toxic soluble oligomers, remains uncertain (Ishii et al. 2017; Glass 2020). There is a possibility that TDP-43 pathology spreads between neurons through a prion-like mechanism (Jo et al. 2020; Chong and Forman-Kay 2016).

The proposed model is based on the following hypotheses regarding the mechanism of TDP-43 aggregation. First, TDP-43 dimers dissociate into monomers, which then translocate from the nucleus to the cytoplasm (Ingólfsson et al. 2023; Oiwa et al. 2023). Subsequent aggregation events occur in the cytoplasm, where monomers convert into pathogenic oligomers with autocatalytic properties (Wells et al. 2021; French et al. 2019); de Boer et al. 2021). These pathogenic oligomers deposit into sticky TDP-43 inclusion bodies, driving their growth (Lye and Chen 2022).

It is important to distinguish between physiological and pathological oligomers. Physiological oligomers play functional roles, such as supporting RNA splicing activity in the nucleus (Lye and Chen 2022). However, as this study focuses on inclusion body growth, the term “oligomers” hereafter refers exclusively to pathological forms.

The developed model simulates the growth of TDP-43 inclusion bodies and predicts the relationship between their radius and time. This model builds upon the author’s previous research, which explored the growth of amyloid-β plaques (Kuznetsov 2024b) and Lewy bodies (Kuznetsov and Kuznetsov 2022; Kuznetsov 2024a). The growth of TDP-43 inclusion bodies is simulated using the methodology developed in Kuznetsov (2024b), Kuznetsov (2024a). The model includes equations that describe the conservation of TDP-43 monomers, dimers, free misfolded oligomers, and misfolded oligomers deposited into inclusion bodies. TDP-43 misfolded oligomers are referred to as pathological and toxic oligomers hereafter. The conversion of TDP-43 monomers into free misfolded oligomers is simulated using the Finke-Watzky (F-W) model.

This paper also proposes a criterion to characterize the neurotoxicity of misfolded TDP-43 oligomers, which are hypothesized to be the main toxic species in ALS and FTLD. An approximate solution for accumulated neurotoxicity, derived under the assumption of dysfunctional degradation machinery for TDP-43 protomers, predicts a linear increase in neurotoxicity from the onset of misfolded oligomer formation.

## 2. Materials and models

### 2.1. Governing equations

According to Oiwa et al. (2023), the initial stage of TDP-43 aggregation involves the monomerization of TDP-43 dimers:

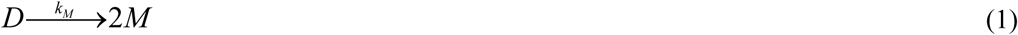

Monomeric TDP-43 lacks functionality and is more likely to escape the nucleus, possibly through passive diffusion, while also being prone to aggregation. As a result, in ALS and FTLD, most TDP-43 inclusions are found in the cytoplasm Pérez-Berlanga et al. (2023). Consequently, the model focuses on simulating TDP-43 inclusion bodies’ formation in the cytoplasm.

French et al. (2019) suggested that TDP-43 misfolded oligomers serve as critical intermediates in the transition to irreversible aggregates. To simulate TDP-43 aggregation, a phenomenological two-step F-W model, as described in Morris et al. (2008), Iashchishyn et al. (2017), was employed. The F-W model, which has been successfully applied to fit experimental data on neurological protein aggregation (Morris et al. 2008), simulates polymer aggregation through two pseudoelementary, parallel, and competing processes: continuous nucleation and autocatalytic growth catalyzed by oligomer surfaces (Finke et al. 2020). The F-W model is characterized by the following transitions:

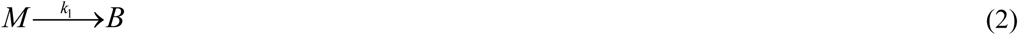

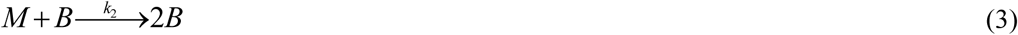

The mathematical model is derived by establishing four conservation statements for dimers, monomers, free misfolded oligomers, and misfolded oligomers deposited into inclusion bodies. The model employs the lumped capacitance approximation, with time, *t*, as the sole independent variable. The dependent variables are listed in Table 1, and the model parameters are summarized in Table 2.

**Table 1.** Dependent variables in the model.

| Symbol | Definition | Unit |
| --- | --- | --- |
| $[B]$ | Molar concentration of free toxic (pathological, misfolded) TDP-43 oligomers | $\mu\text{M}$ |
| $[D]$ | Molar concentration of TDP-43 dimers, representing the native form of TDP-43 | $\mu\text{M}$ |
| $[I]$ | Concentration of toxic TDP-43 oligomers incorporated into inclusion bodies | $\mu\text{M}$ |
| $[M]$ | Molar concentration of TDP-43 monomers | $\mu\text{M}$ |
| $r_{IB}$ | Radius of the growing inclusion body, assuming it takes on a spherical form | $\mu\text{m}$ |

**Table 2.**
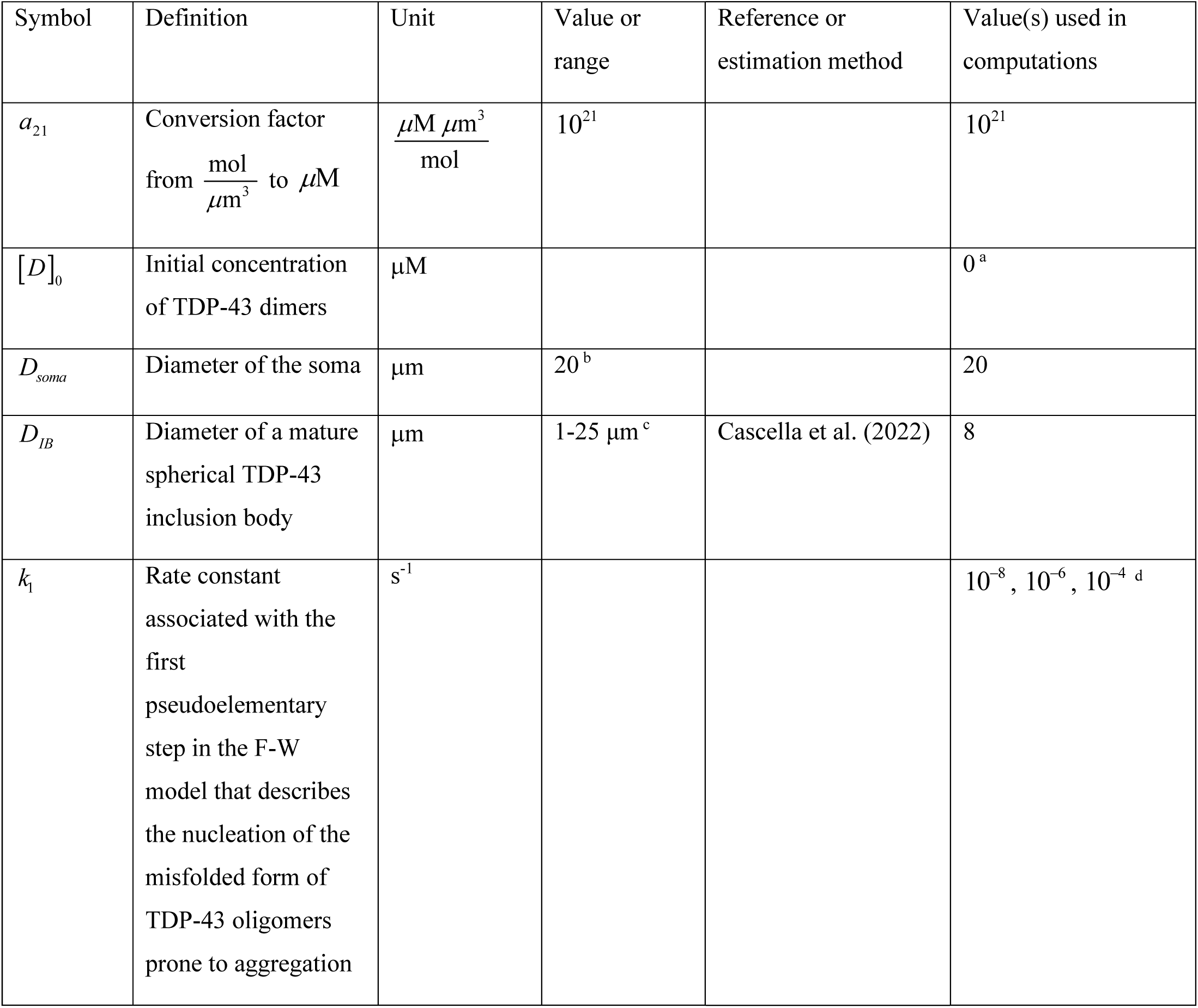
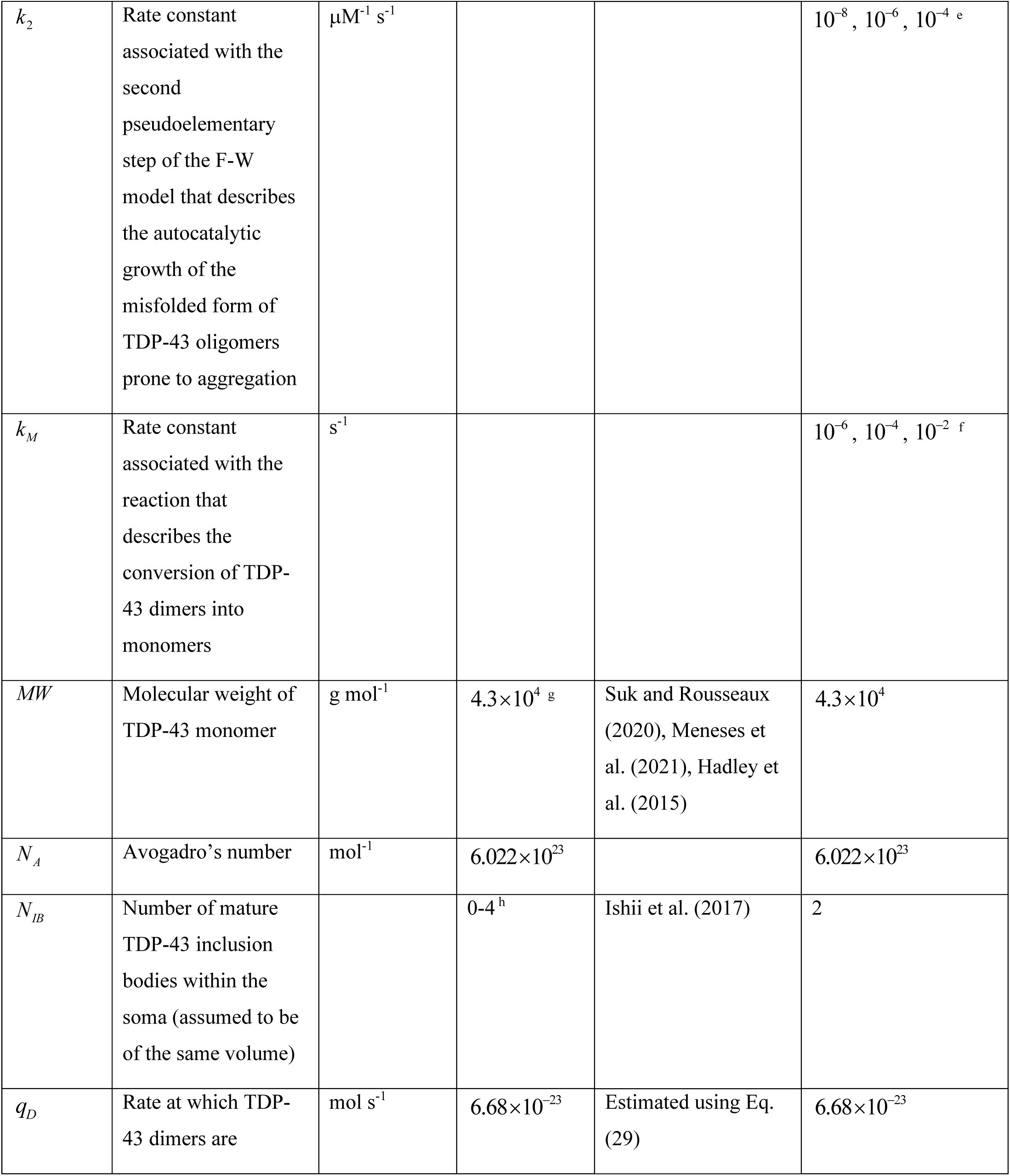
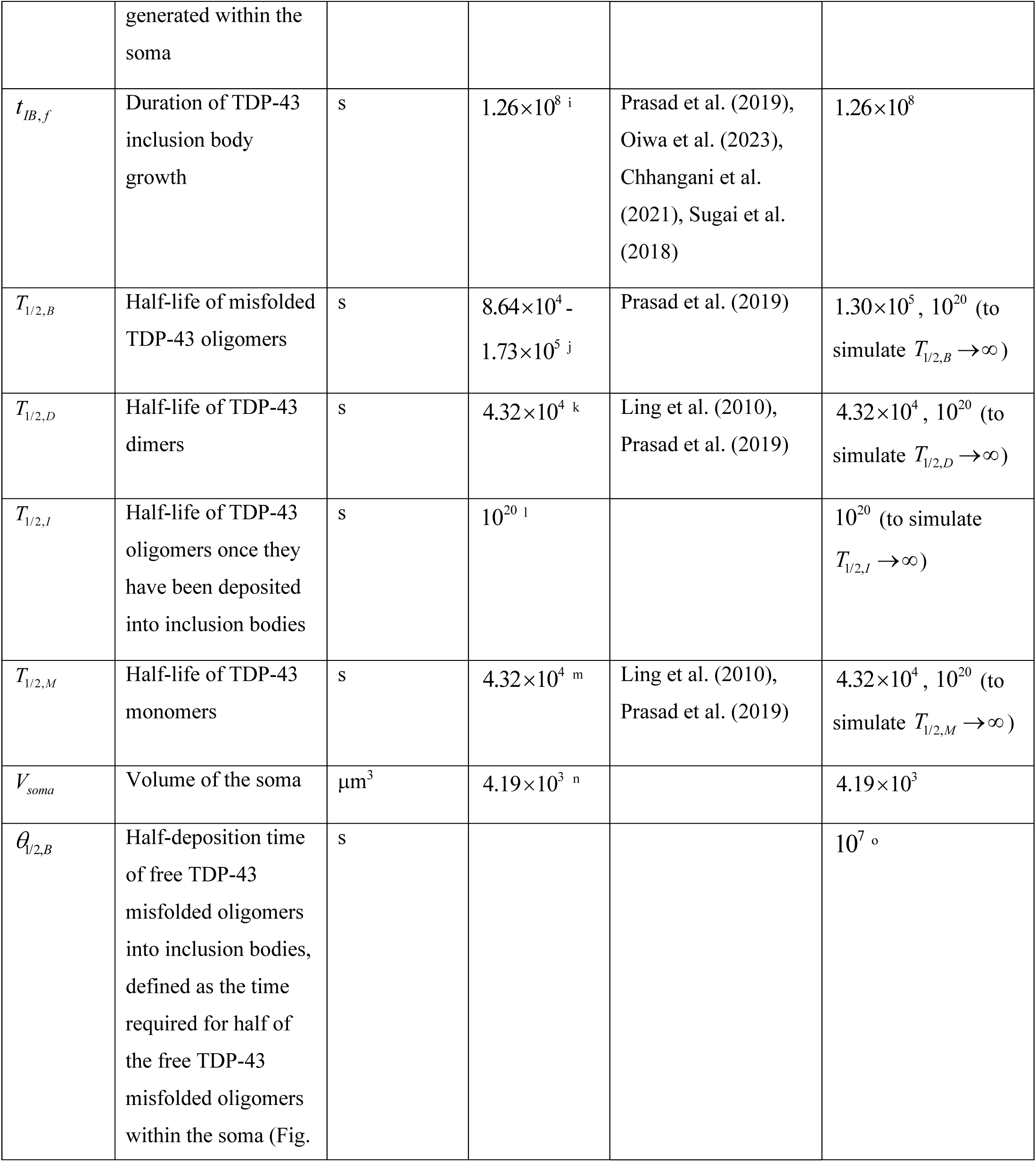
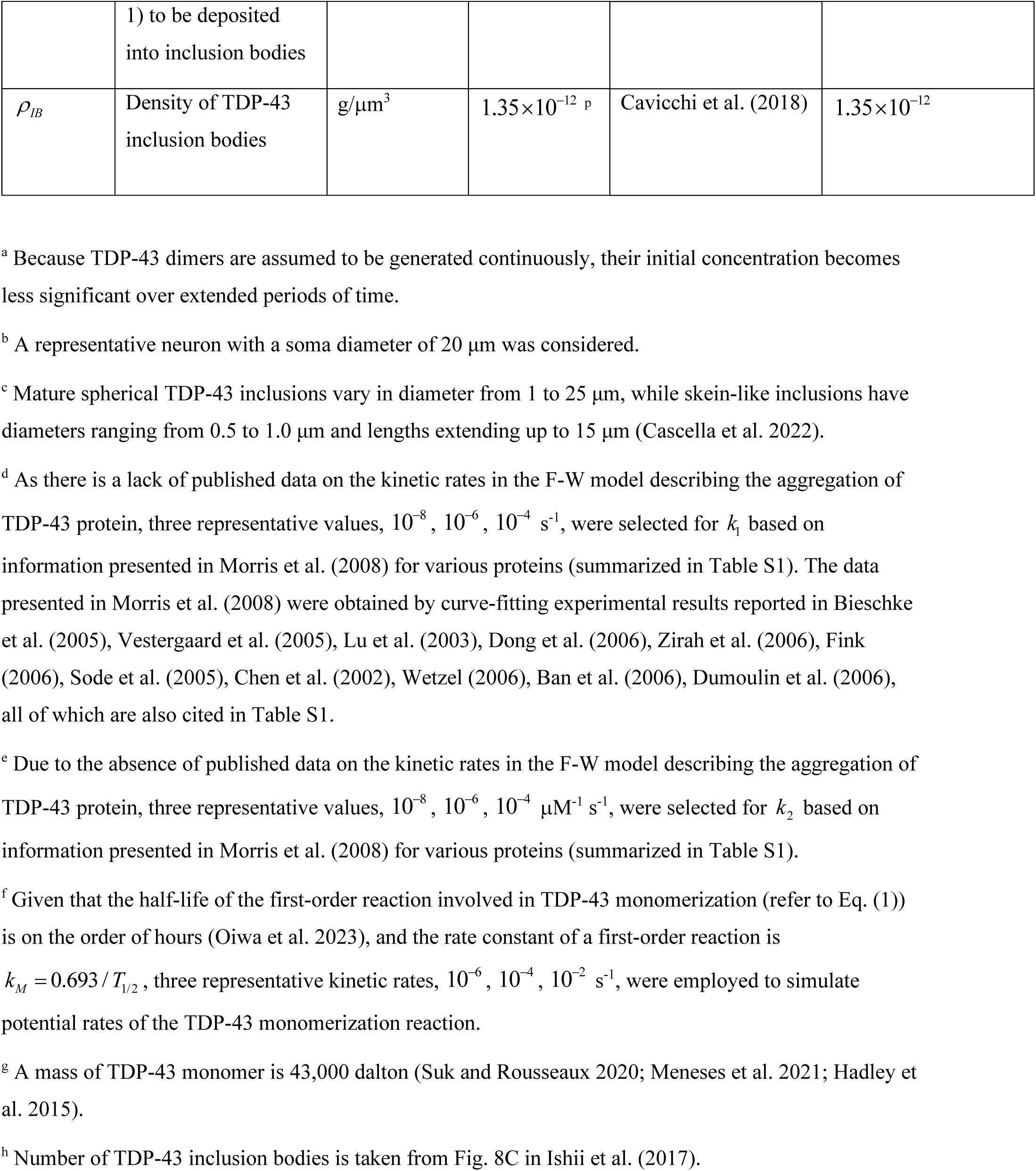
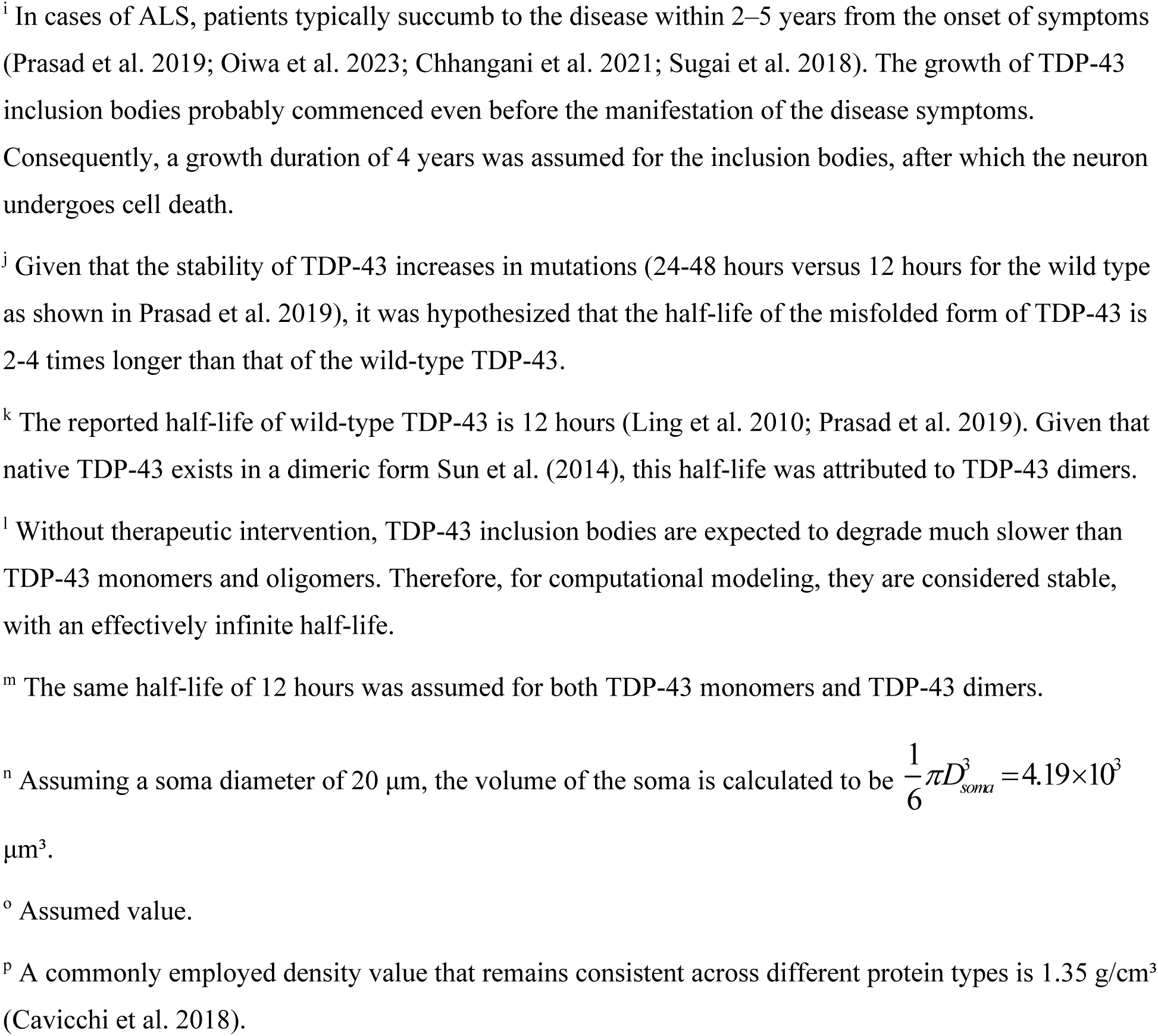
Model parameters along with their estimated values.

The conservation of TDP-43 dimers in the soma is expressed by the following equation:

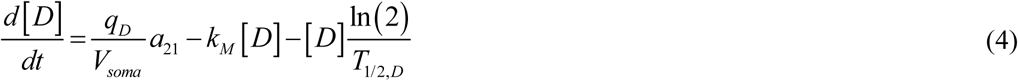

In Eq. (4), 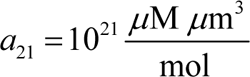 represents the conversion factor from to 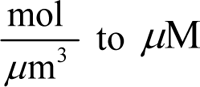. The first term on the right-hand side of Eq. (4) simulates the rate at which TDP-43 dimers, a native form of TDP-43, are produced in the soma. TDP-43 is an RNA-binding protein (Scherer et al. 2024). TDP-43 regulates its own expression by binding to specific sequences in its mRNA, thereby reducing its production rate (Ayala et al. 2011; Koehler et al. 2022). Therefore, it is likely that *q_D_* decreases as the concentration of TDP-43 dimers increases. Since the exact dependence of *q_D_* on [*D*] is unknown, *q_D_* is assumed to be constant in this study. The second term is formulated by applying the law of mass action to describe the rate of reaction in Eq. (1). The third term simulates the degradation of TDP-43 dimers due to their finite half-life. The conservation of TDP-43 monomers in the soma is described by the following equation:

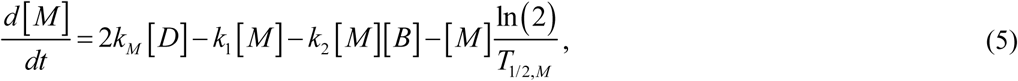

where the first term on the right-hand side describes the generation of TDP-43 monomers through their conversion from dimers. The second term simulates the decrease of monomer concentration resulting from their transformation into the misfolded active form capable of autocatalysis (pathological oligomers). In the F-W model, this process is conceptualized as a nucleation process (refer to Eq. (2)). The third term accounts for the decrease of monomer concentration due to autocatalytic conversion into misfolded oligomers (refer to Eq. (3)). The fourth term represents the degradation of TDP-43 monomers due to their finite half-life.

By expressing the conservation of TDP-43 misfolded oligomers, the following is derived:

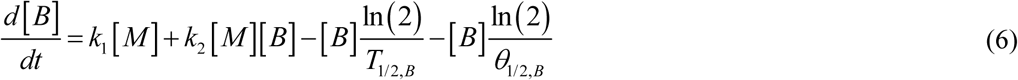

The first and second terms on the right-hand side of Eq. (6) simulate the generation of TDP-43 misfolded oligomers resulting from their production from monomers through nucleation and autocatalytic processes, respectively. The third term on the right-hand side represents the decline of TDP-43 misfolded oligomers due to their limited half-life.

The formation of TDP-43 inclusion bodies from sticky misfolded oligomers is modeled similarly to colloidal suspension coagulation in Boltachev and Ivanov (2020). In the model, free misfolded TDP-43 oligomers (*B*) are assumed to have a half-life of *θ*_1/ 2, *B*_. These misfolded oligomers deposit into TDP-43 inclusion bodies, and the concentration of the deposited oligomers is denoted as [*I*].

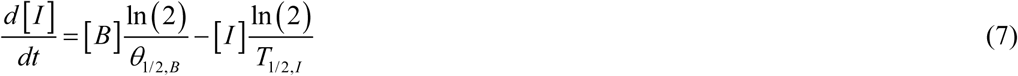

In Eq. (7), the first term on the right-hand side corresponds to the fourth term in Eq. (6) with the opposite sign, while the second term represents the half-life of deposited TDP-43 oligomers incorporated into inclusion bodies.

Eqs. (4)-(7) are solved with the following initial conditions:

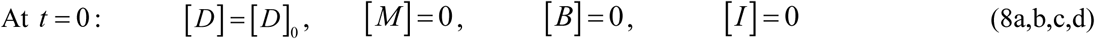

### 2.2. Criterion for assessing the accumulated neurotoxicity of pathological TDP-43 oligomers

The role of TDP-43 inclusion bodies in the neural tissue of individuals diagnosed with ALS or FTLD remains debated, as they may function as primary neurotoxic agents, harmless byproducts, or protective structures sequestering misfolded proteins (Baloh 2011). Liu et al. (2013) proposed that although TDP-43 aggregation may be correlated with neuronal cell death, it is not necessarily its direct cause. Accordingly, this study hypothesizes that TDP-43 neurotoxicity arises from misfolded TDP-43 oligomers and progressively accumulates over time as neurons are exposed to these toxic species, whereas inclusion bodies are primarily neuroprotective.

It is important to distinguish between functional and pathological (misfolded) oligomers of TDP-43. Functional oligomerization is crucial for TDP-43’s role in nuclear splicing regulation and its incorporation into cytoplasmic stress granules during cellular stress (Pérez-Berlanga et al. 2023). French et al. (2019) noted similarities between TDP-43 pathological aggregation and the aggregation processes of tau, α-synuclein, and amyloid β, where oligomers are thought to be the primary toxic species. However, the neurotoxicity of TDP-43 oligomers remains a somewhat controversial topic. Fang et al. (2014) reported that TDP-43 oligomers could cause neurotoxicity and neurite degeneration. The neurotoxicity of TDP-43 oligomers was also reported in Lee et al. (2019), Lye and Chen (2022). Yang et al. (2024) produced recombinant TDP-43 oligomers and observed minimal or no neurotoxicity. However, these oligomers differed from those naturally occurring in cells. The structural and toxic characteristics of native TDP-43 oligomers may also be influenced by post-translational modifications, genetic mutations, and interactions with various cofactors (Yang et al. 2024). Kitamura et al. (2024) proposed that TDP-43 hetero-oligomers, formed by incorporating other proteins, exhibit greater neurotoxicity than their homo-oligomer counterparts. Lye and Chen (2022) provided a comprehensive review of misfolded TDP-43 oligomers, demonstrating their neurotoxic effects in both in vitro and in vivo studies.

The precise mechanism underlying the neurotoxicity of misfolded TDP-43 oligomers remains unknown. Some TDP-43 oligomers may be inherently more neurotoxic than others, or neurotoxicity could result from a heterogeneous mixture of oligomers with varying structures, stability, and concentrations. The mixture of oligomers may induce nonspecific neurotoxicity by interacting with membrane proteins, disrupting membrane lipids, triggering oxidative stress, and altering membrane dielectric properties and ion permeability (Benilova et al. 2012). If neurotoxicity results from the combined effects of various oligomers, a parameter quantifying the accumulated time-dependent damage caused by free misfolded TDP-43 oligomers can be defined, following Kuznetsov (2025a), Kuznetsov (2025b), as:

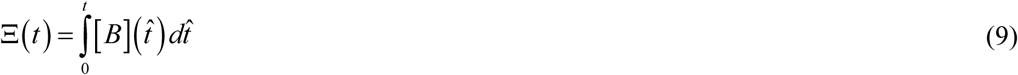

If multiple distinct oligomeric TDP-43 species contribute to neurotoxicity with different weighting factors *ω*_1_ and *ω*_2_, then the neurotoxicity criterion can be reformulated as:

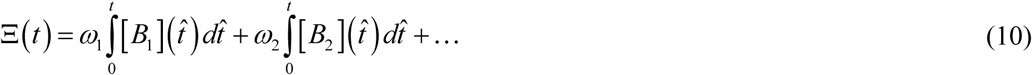

Eqs. (9) and (10) provide microscale, or cellular-level, criteria for assessing TDP-43 neurotoxicity. These equations enable the evaluation of neurotoxicity for individual neurons.

### 2.3. Approximate solution for the case with *T*_1/2,*D*_ →∞, *T*_1/2,*M*_ →∞, *T*_1/2,*B*_ →∞, and *T*_1/2,*I*_ →∞

In the case of dysfunctional proteolytic machinery — namely, *T*_1/ 2,*D*_ →∞, *T*_1/ 2,*M*_ →∞, and *T*_1/2,*B*_ →∞ — the numerical simulations in Figs. S1-S3 indicate that the concentrations of TDP-43 dimers, monomers, and free misfolded oligomers—denoted as [*D*], [*M*], and [*B*], respectively—gradually converge over time, reaching asymptotic steady-state values. This section focuses on determining these equilibrium concentrations. The analytical solution derived here, labeled as “approximate” in the figures, is obtained by applying steady-state conditions to Eqs. (4)-(6), yielding the following equations:

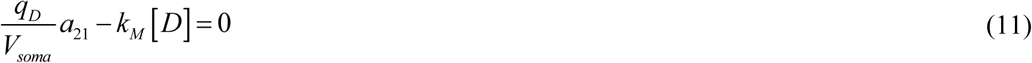

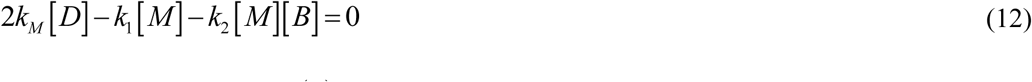

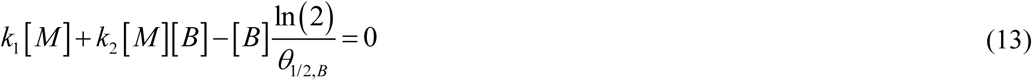

Rearranging Eq. (11) to express [*D*] yields:

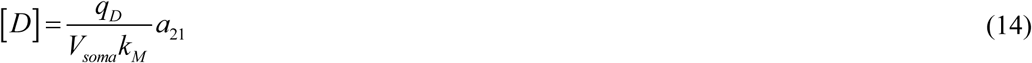

By adding Eq. (11) twice to Eqs. (12) and (13), the following expression is obtained:

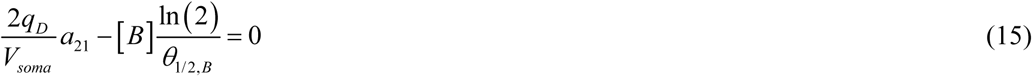

Rearranging Eq. (15) to express [*B*] gives:

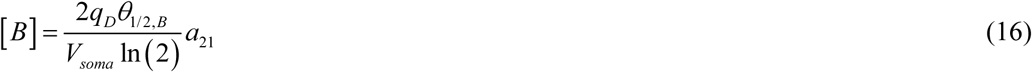

Incorporating Eqs. (14) and (16) into Eq. (12) and solving for [*M*] gives:

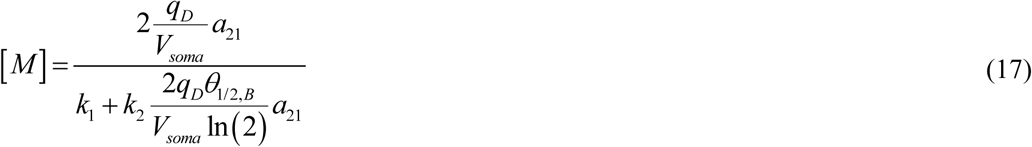

The numerically predicted concentration of dimers, [*D*], quickly approaches a constant value, which is determined by the balance between dimer production and their conversion into monomers (Fig. S1). Fig. S1 demonstrates a strong agreement between the numerical and approximate (given by Eq. (14)) solutions for [*D*] once time exceeds 0.002 years. As monomers convert into oligomers, the numerically predicted concentration of monomers, [*M*], decreases over time, initially rapidly and then gradually (Fig. S2). The numerical and approximate (given by Eq. (17)) solutions for [*M*] converge at large times, but the time it takes for the solutions to align depends on the value of *k*_1_. The convergence is slower for larger values of *k*_1_, taking approximately 1.5 years for the numerical solution to match the approximate solution when *k* = 10^−4^ s^-1^. Note that [*M*] converges to distinct asymptotic values, which depend on *k* (Eq. (17)). The numerical solution for the concentration of free misfolded TDP-43 oligomers, [*B*], exhibits a gradual increase to an asymptotic value over time, with the numerical and analytical (given by Eq. (16)) solutions (the latter gives a constant asymptotic value) converging after approximately 2.5 years (Fig. S3).

As *T*_1/ 2,*I*_ →∞, Eq. (7) can be reformulated as:

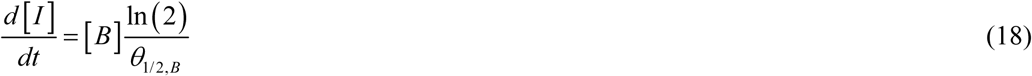

By incorporating Eq. (16) into Eq. (18), the following expression is obtained:

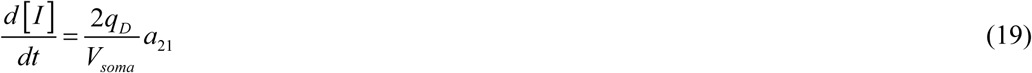

Integrating Eq. (19) and applying the initial condition (8d) yields the following expression:

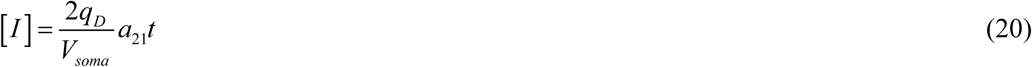

Fig. S4 illustrates that after approximately one year, the numerical solution for [*I*] exhibits a linear increase. The numerical and approximate solutions for [*I*] are similar but not identical. The discrepancy arises from differences between the numerical and approximate solutions for the concentration of free misfolded TDP-43 oligomers, [*B*] (Fig. S3), leading to initial variations in the predicted concentrations of deposited oligomers, [*I*], which persist over time.

Substituting the approximate solution for [*B*] from Eq. (16) into Eq. (9) yields the following expression for accumulated neurotoxicity:

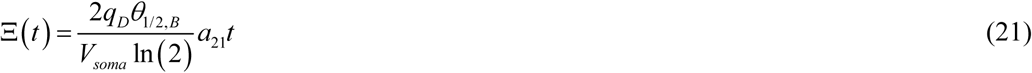

Eq. (21) provides an approximate solution for Ξ(*t*), indicating that accumulated neurotoxicity increases in direct proportion to disease duration.

### 2.4. Modeling the growth dynamics of TDP-43 inclusion bodies

The growth of TDP-43 inclusion bodies over time (see Fig. 1) is simulated using the following approach. The total number of TDP-43 monomers incorporated into a single inclusion body at time *t* is given by the following equation, adapted from Watzky et al. (2008):

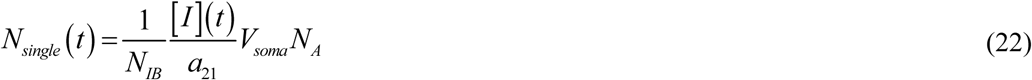

**Fig. 1.**
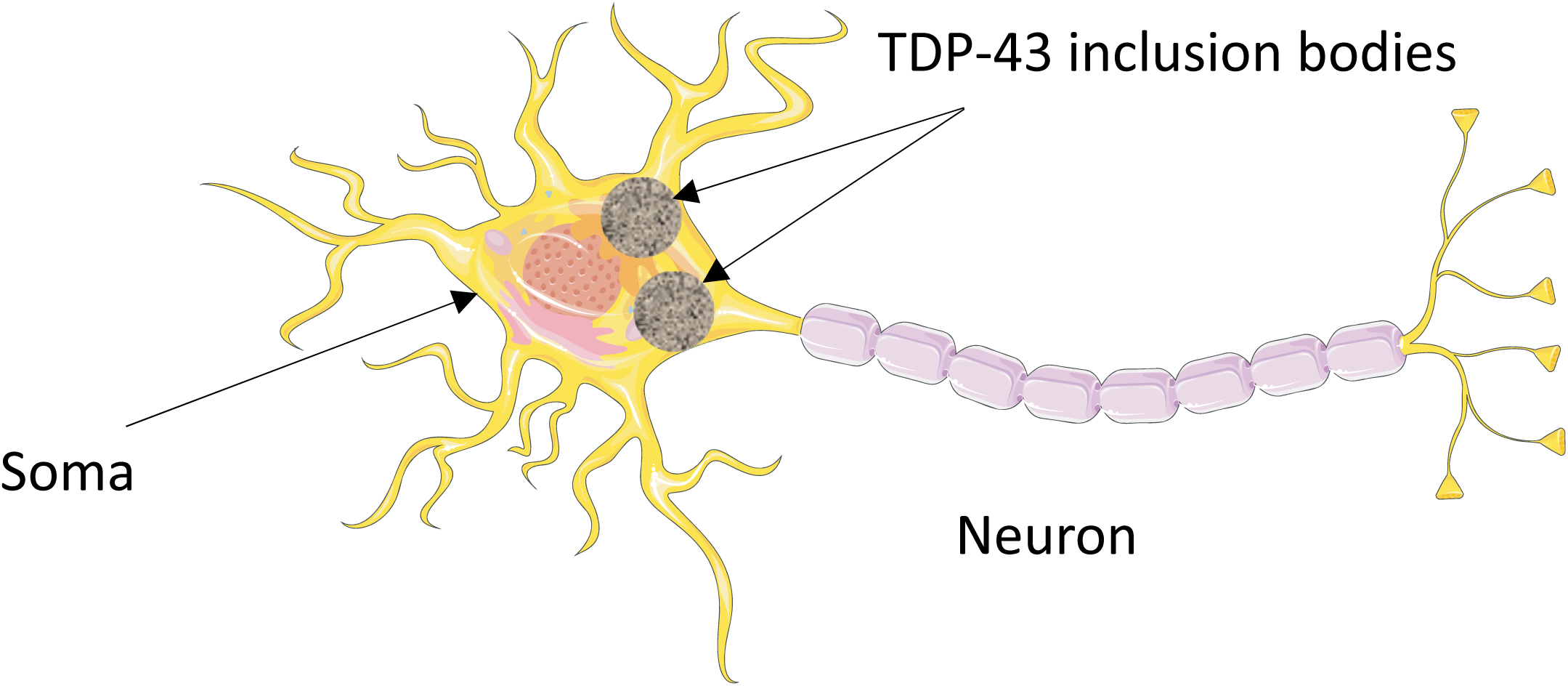
A diagram depicting a neuron with two TDP-43 inclusion bodies of equal size located in the soma. The inclusion bodies shown in Fig. 1 are not drawn to scale. *Figure generated with the aid of servier medical art, licensed under a creative commons attribution 3.0 generic license.* http://Smart.servier.com.

The F-W model does not distinguish between the molecular weights of monomers (native form) and misfolded oligomers (active form capable of autocatalysis); see Eqs. (2) and (3). In Eq. (13), *N_IB_* represents the number of inclusion bodies in the soma, with the assumption that all inclusion bodies have equal volumes.

Alternatively, *N_single_* (*t*) can be calculated as follows (Watzky et al. 2008):

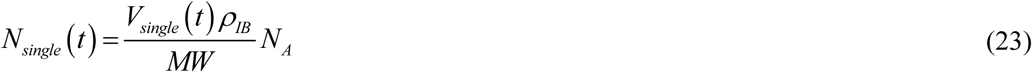

where *MW* denotes the molecular weight of TDP-43 monomers, defined as the sum of the atomic weights of all the atoms within a monomer.

By equating the right-hand sides of Eqs. (22) and (23) and solving for the volume of a single inclusion body, the following equation is obtained:

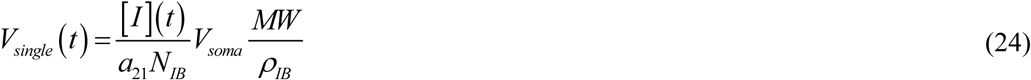

Substituting Eq. (20) into Eq. (24) yields:

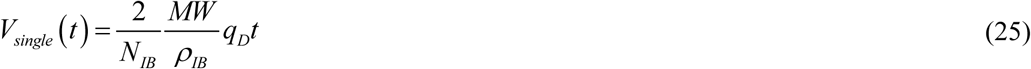

This implies that the volume of a single inclusion body in the soma increases linearly with both the rate of TDP-43 dimer generation and time.

The model includes separate equations for free misfolded TDP-43 oligomers (phase *B*) and oligomers deposited into TDP-43 inclusions (phase *I*). While free oligomers are initially highly dynamic structures, the model does not require resolving their shape—only an assumption about the shape of the inclusion bodies (phase *I*) is necessary.

TDP-43 inclusions within the soma can vary in shape, ranging from skein-like to rounded forms (Cascella et al. 2022). To simplify calculations, all inclusion bodies are assumed to be spherical and of uniform size. The volume of a spherical inclusion body is

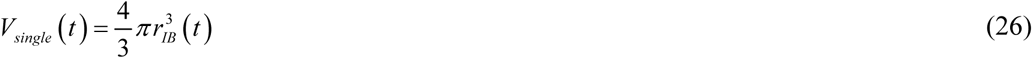

It is important to note that the model predicts the volume of the inclusion body, as in Eq. (25). In Eq. (26), the parameter *r_IB_* can be interpreted as the effective radius, representing the radius of a sphere with the same volume as the inclusion body. This assumption does not impact the model’s predictions regarding the growth of inclusion volume over time, as the model is independent of convexity. Therefore, variations in radius across the inclusion body structure do not affect the results.

Equating the right-hand sides of Eqs. (24) and (26) and solving for the radius of the inclusion body yields the following result:

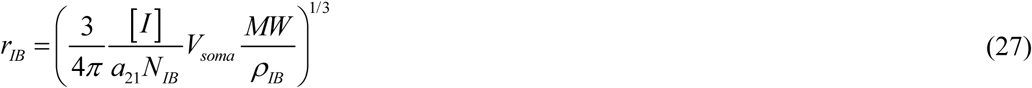

Using the approximate solution for [*I*] from Eq. (20) in Eq. (27), the following approximate solution for *r_IB_* for the case of *T*_1/ 2,*D*_ →∞, *T*_1/ 2,*M*_ →∞, *T*_1/2,*B*_ →∞, and *T*_1/ 2,*I*_ →∞ is derived:

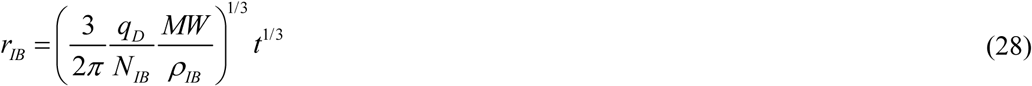

As indicated by Eq. (28), the radius of the inclusion body grows proportionally to the cube root of time. This growth is influenced by the production rate of TDP-43 dimers, *q_D_*.

Eq. (28) can be utilized to estimate the rate at which TDP-43 dimers are generated in the soma. If, after the growth period *t_IB_*, *_f_* the inclusion body’s radius attains the size *r_IB_*, *_f_*, then, according to Eq. (28), *q_D_* can be determined as:

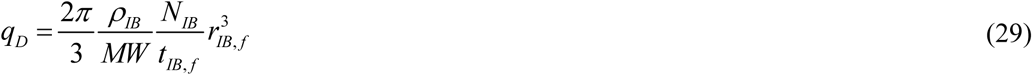

### 2.5. Sensitivity analysis of the inclusion body radius and accumulated neurotoxicity of free misfolded TDP-43 oligomers with respect to model parameters

The study investigated the sensitivity of the inclusion body’s radius, *r_IB_*, to various parameters in Eq. (28). Local sensitivity coefficients (Beck and Arnold 1977; Zadeh and Montas 2010; Zi 2011; Kuznetsov and Kuznetsov 2019) were computed, representing the first-order partial derivatives of the inclusion body’s radius with respect to, for example, the rate of TDP-43 dimer production, *q_D_*, denoted by

The dimensionless relative sensitivity coefficients (Zadeh and Montas 2010; Kacser et al. 1995) were determined as the ratio of the relative change in the inclusion body’s radius, Δ*r_IB_* / *r_IB_*, to the relative change in the respective parameter, in this case, Δ*q_D_* / *q_D_*:

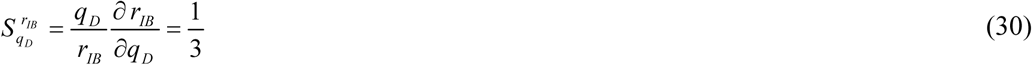

The approximate solution provided in Eq. (28) was used to calculate the derivative in Eq. (30). Importantly, the result in Eq. (30) is time-independent, indicating that the sensitivity remains constant at any given time. The positive sign of *S ^rIB^* suggests that *r* increases with an increase in *q_B_*. Similarly,

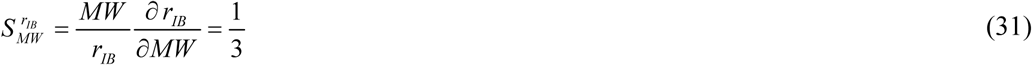

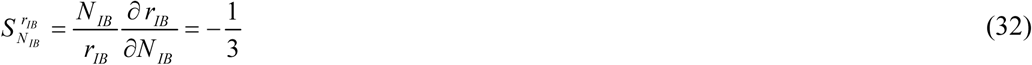

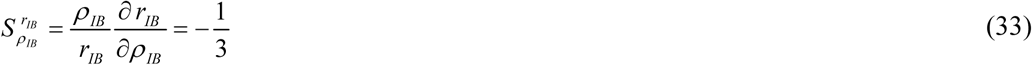

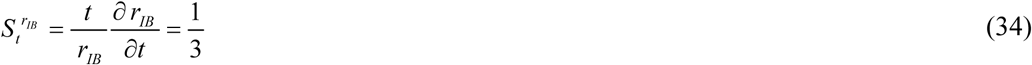

Similarly, the dimensionless relative sensitivity coefficients characterizing the dependence of accumulated neurotoxicity on model parameters can be obtained using Eq. (21):

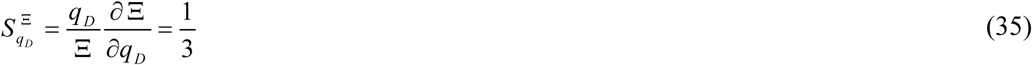

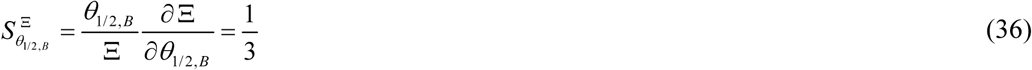

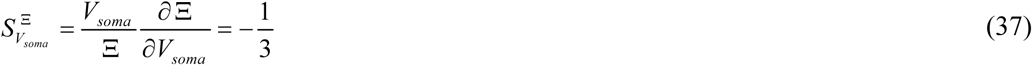

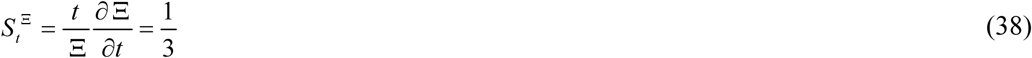

At large times, the sensitivity of *r_IB_* to the kinetic constants *k*_1_, *k*_2_, and *k_M_* is zero, as derived from the approximate solution (28). However, since this solution is valid only for long-term behavior, the numerical solution was used to evaluate the sensitivity of *r_IB_* to these kinetic constants at earlier times. For instance:

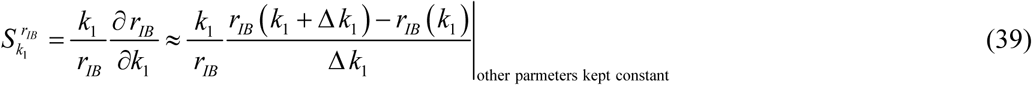

where Δ *k* = 10^−3^ *k* represents the step size. Sensitivity coefficients were computed using different step sizes to ensure that they remained unaffected by variations in step size.

Similarly, for large times, the sensitivity of Ξ to the kinetic constants *k*_1_, *k*_2_, and *k_M_* is also zero, as indicated by the approximate analytical solution (21). Given that this solution applies only to large timescales, numerical simulations were employed to determine the sensitivity of Ξ to these kinetic constants at shorter time scales. For example:

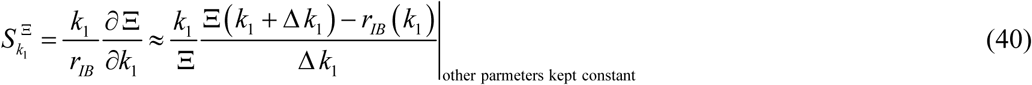

### 2.6. Numerical solution

Eqs. (4)–(7), with initial conditions (8), were solved using MATLAB’s well-validated ODE45 solver (MATLAB R2020b, MathWorks, Natick, MA, USA). To guarantee solution accuracy, the error tolerance parameters, RelTol and AbsTol, were set to 1e-10. The convergence was established by calculating the residuals.

## 3. Results

Figs. 2, 5, 7, 9, 11, and 13 illustrate the radius of a TDP-43 inclusion body in a scenario where the soma contains two equally sized inclusion bodies. The model predicts the concentration of free misfolded TDP-43 oligomers, which are gradually deposited into the inclusion bodies, driving their growth. To simplify the interpretation, both inclusion bodies are assumed to be spherical, allowing the total volume of TDP-43 inclusion bodies to be expressed in terms of the radii of two identical spheres. Figs. 3, 6, 8, 10, 12, and 14 illustrate the accumulated neurotoxicity resulting from free misfolded TDP-43 oligomers.

**Fig. 2.**
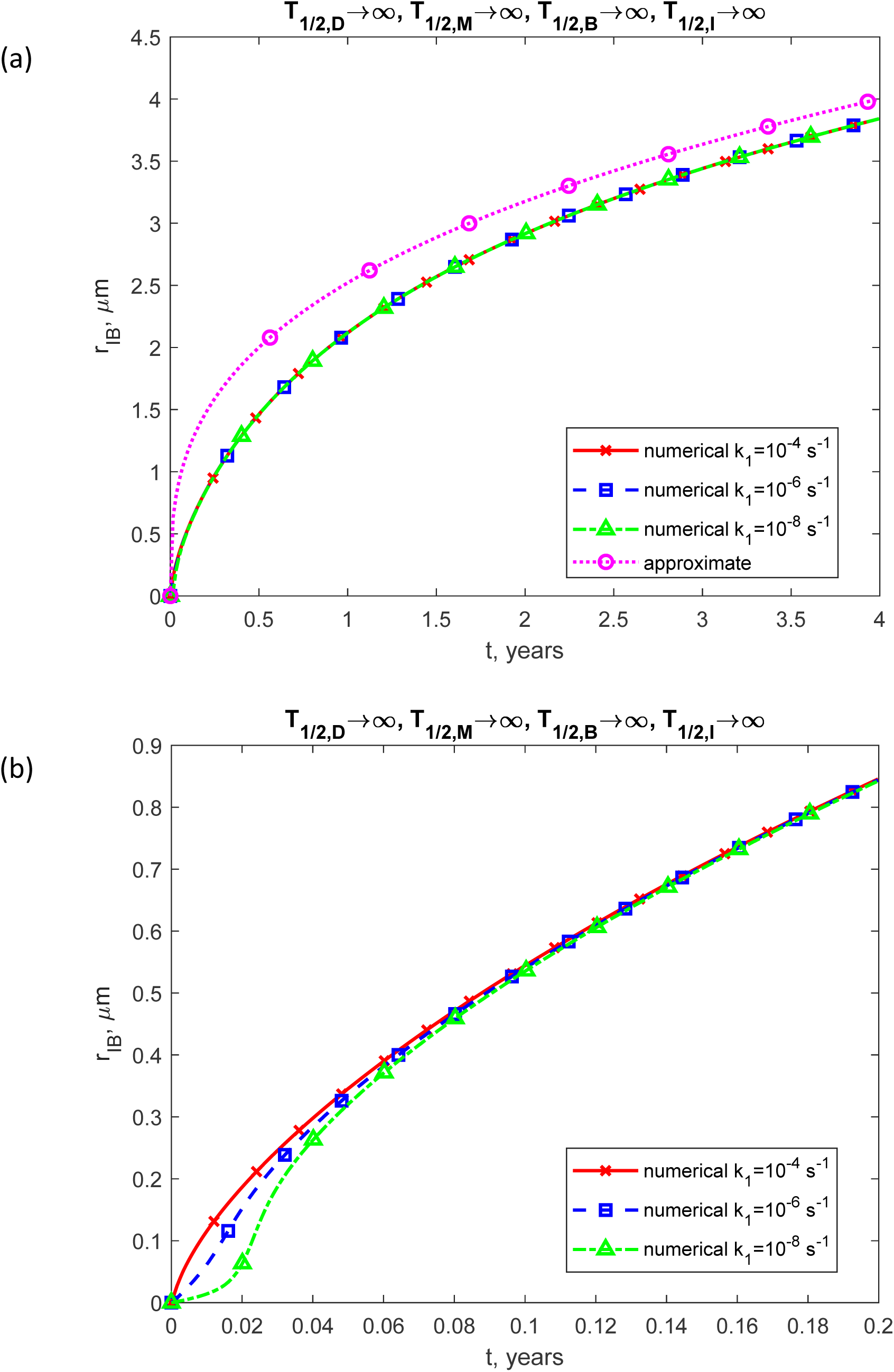
(a) The radius of the growing inclusion body over time (in years) is shown for three different values of *k*_1_. The model predicts the total volume of all inclusion bodies in the soma as *N_IB_V_single_*. Assuming all inclusion bodies have the same volume, the volume of a single inclusion body, denoted by *V_single_*, is calculated in Eqs. (24) and (25). To facilitate interpretation, the predicted volume is converted into the radius of a sphere with an equivalent volume. (b) Similar to Fig. 2a, but with a focus on the specific time range of [0, 0.2 years]. The concentration of the misfolded form of TDP-43, [*B*], is obtained by numerically solving Eqs. (4)–(7) with initial conditions (8), while the radius of the inclusion body is calculated using Eq. (27). The approximate solution for the radius is determined using Eq. (28). The presented scenario assumes dysfunctional protein degradation machinery, with *T*_1/ 2,*D*_ →∞, *T*_1/ 2,*M*_ →∞, *T*_1/2,*B*_ →∞, and *T*_1/2,*I*_ →∞. The case with *k*_2_ = 10^−6^ μM^-1^ s^-1^, *k_M_* = 10^−4^ s^-1^, and *θ*_1/2,*B*_ =10^7^ s.

**Fig. 3.**
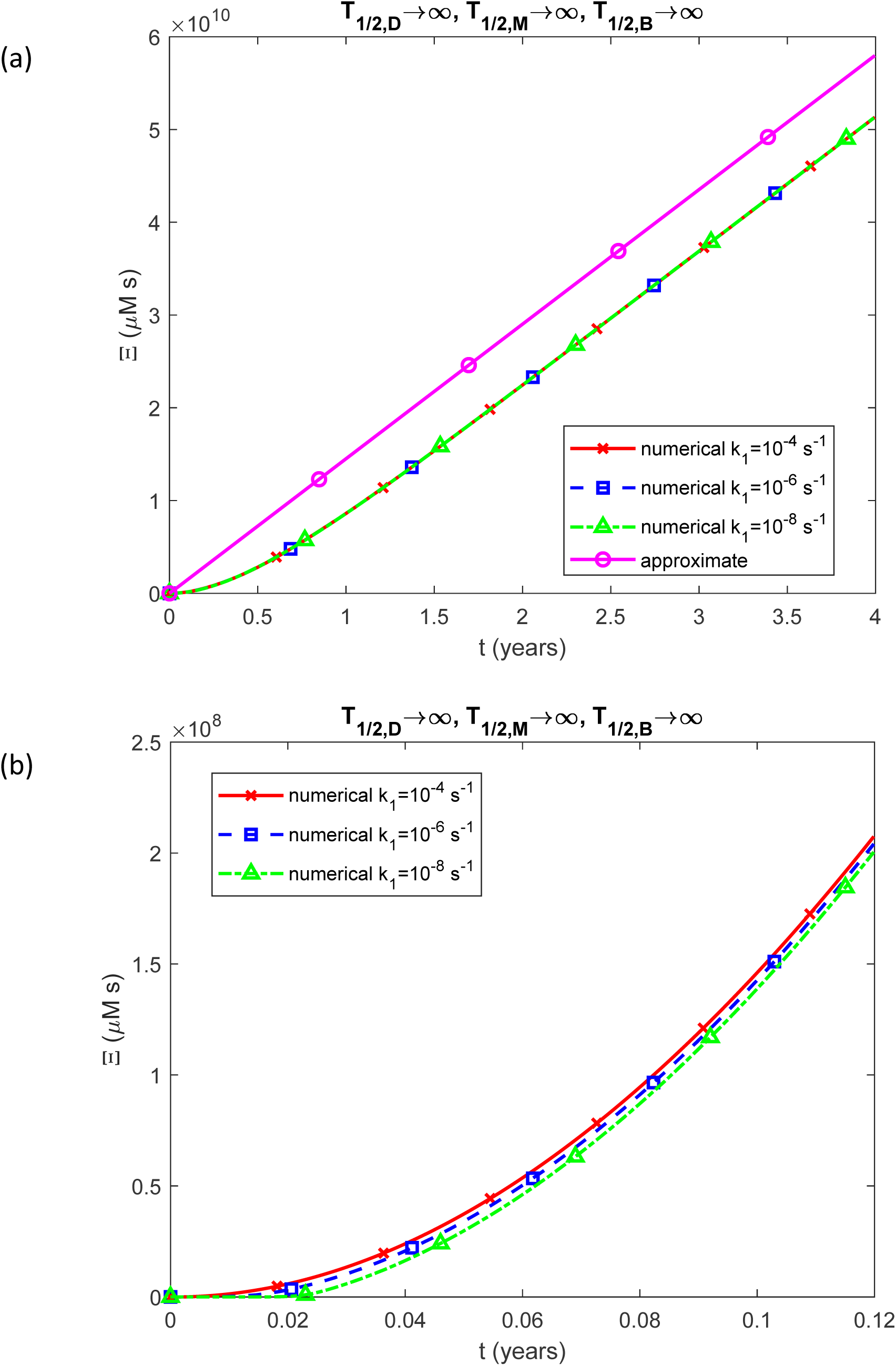
(a) Accumulated neurotoxicity of pathological TDP-43 oligomers, Ξ, as a function of time (in years). Results are shown for three different values of *k*_1_. (b) Similar to Fig. 3a, but with a focus on the specific time range of [0, 0.12 years]. The numerical solution is obtained by solving Eqs. (4)-(7) with initial conditions (8). The approximate solution is determined using Eq. (21). The presented scenario assumes dysfunctional protein degradation machinery with *T*_1/ 2,*D*_ →∞, *T*_1/ 2,*M*_ →∞, and *T*_1/ 2,*B*_ →∞. The case with *k*_2_ = 10^−6^ μM s^-1^, *k*= 10 s^-1^, and *θ* =10^-4^ s^-1^. Note that the approximate solution does not depend on *k*_1_.

**Fig. 4.**
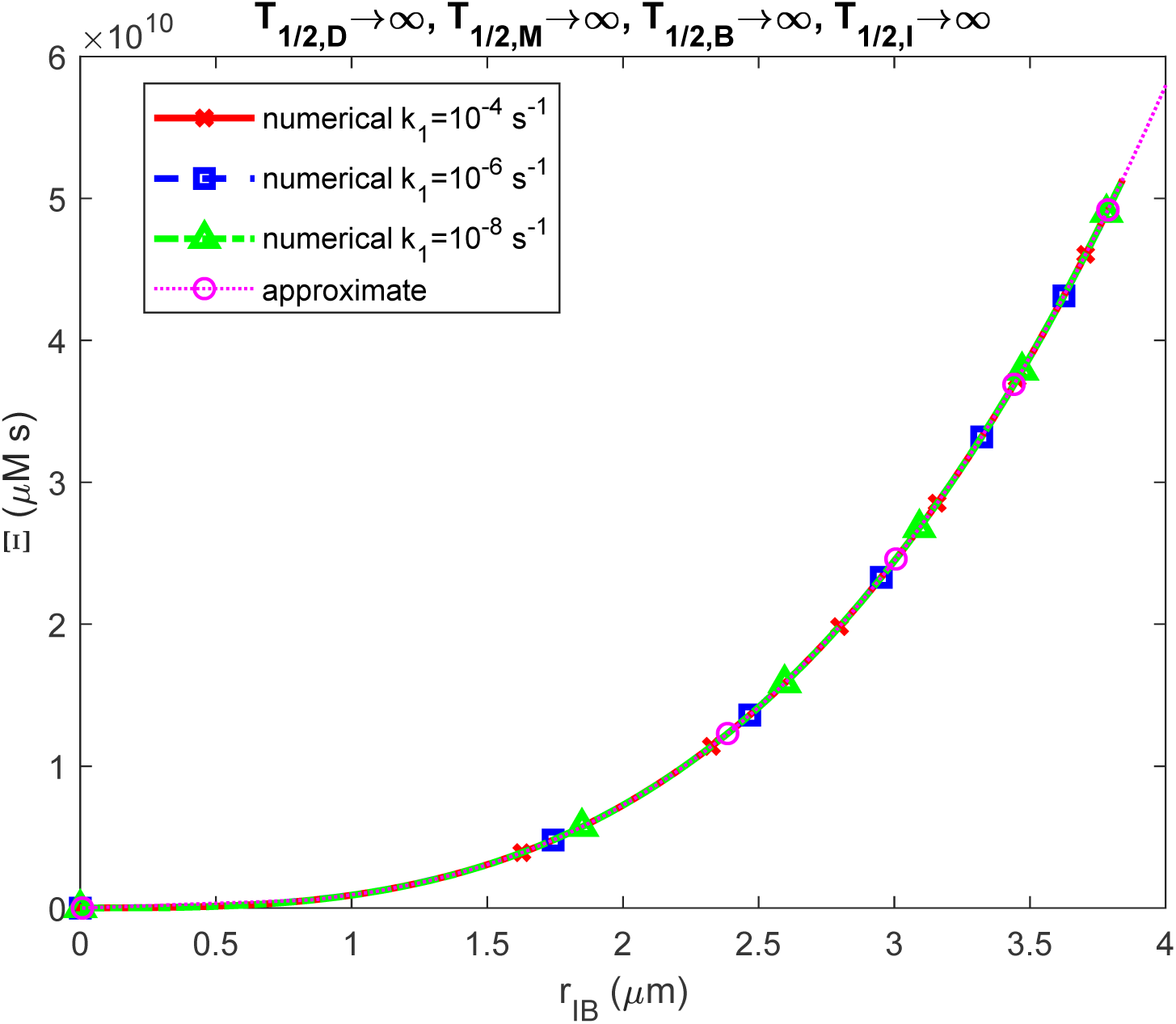
Accumulated neurotoxicity of pathological TDP-43 oligomers, Ξ, as a function of the growing inclusion body radius, *r_IB_*. Results are shown for three different values of *k*_1_. The depicted scenario assumes a finite half-life for TDP-43 dimers, monomers, and pathological TDP-43 oligomers ( *T*_1/ 2,*D*_ →∞, *T*_1/ 2,*M*_ →∞, and *T*_1/ 2,*B*_ →∞), as well as an intermediate half-deposition time for free misfolded TDP-43 oligomers into inclusion bodies (*θ* =10^7^ s).

**Fig. 5.**
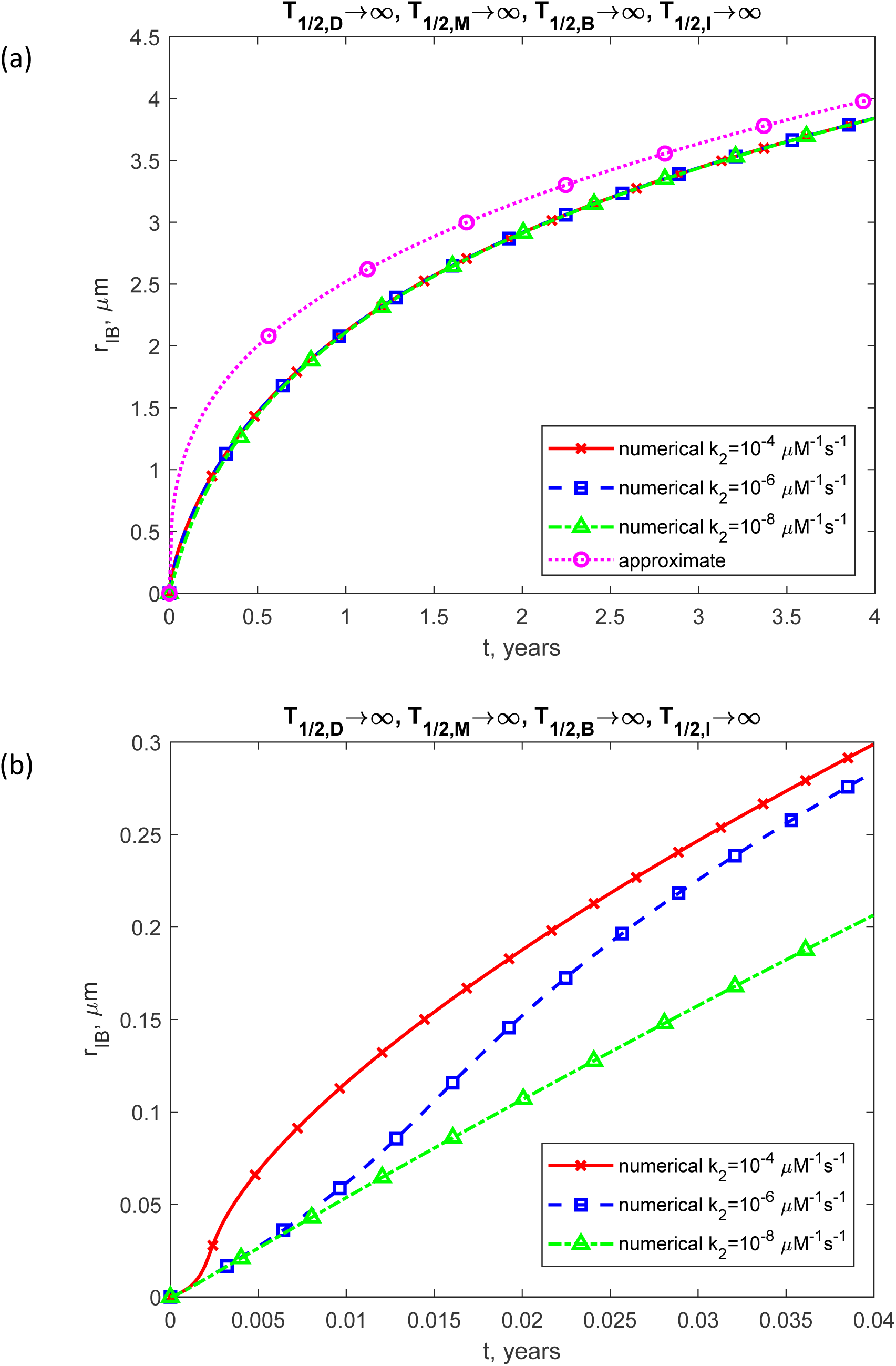
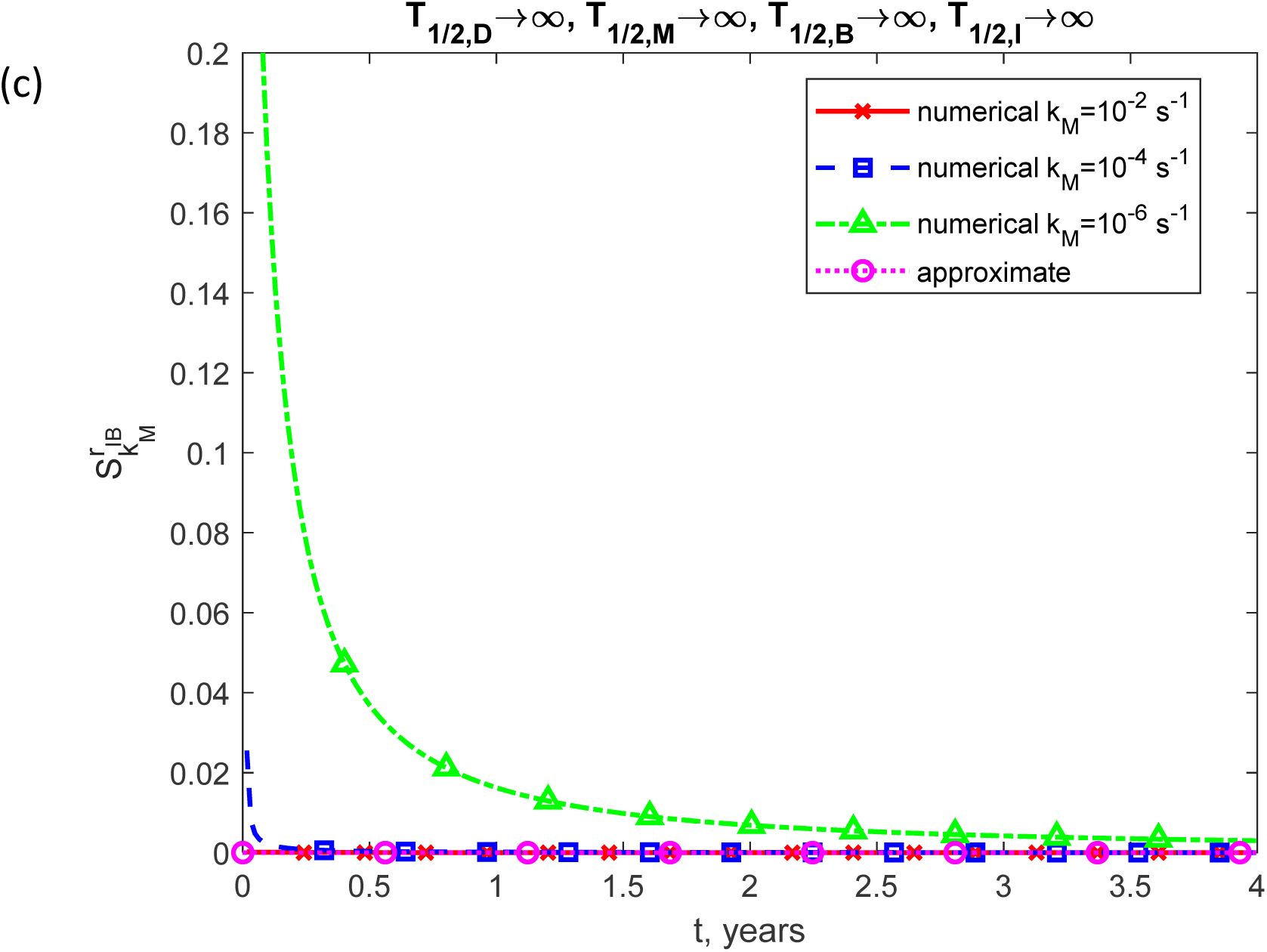
(a) The radius of the growing inclusion body as a function of time (in years). Results are shown for three different values of *k*_2_. (b) Similar to Fig. 5a, but with a focus on the specific time range of [0, 0.04 years]. The approximate solution is determined using Eq. (28). The presented scenario assumes dysfunctional protein degradation machinery with *T*_1/ 2,*D*_ →∞, *T*_1/ 2,*M*_ →∞, and *T*_1/ 2,*B*_ →∞. The case with *k* = 10^−6^ s^-1^, *k* = 10^−4^ s^-1^, and *θ* =10^7^ s.

**Fig. 6.**
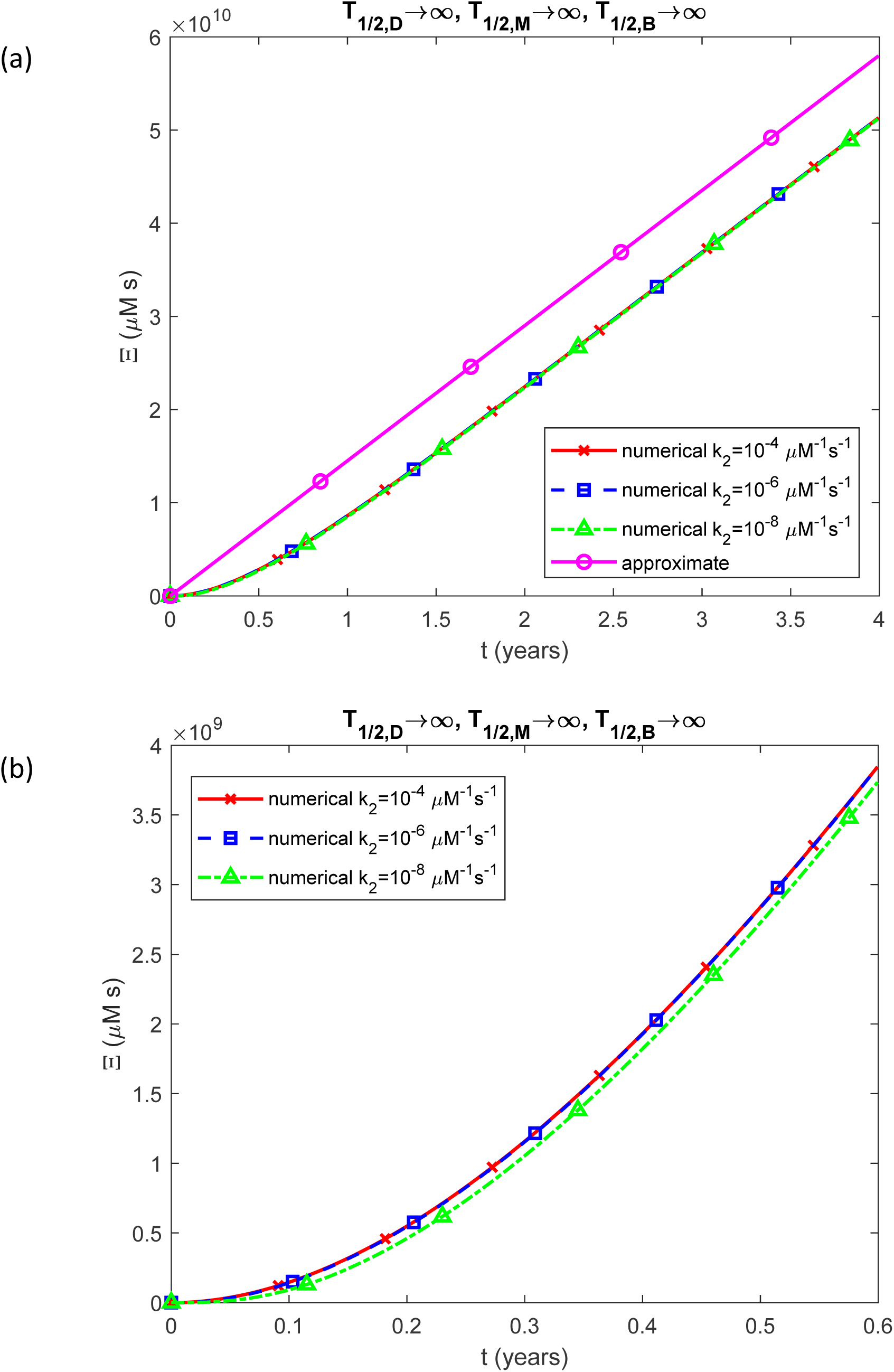
(a) Accumulated neurotoxicity of pathological TDP-43 oligomers, Ξ, as a function of time (in years). Results are shown for three different values of *k*_2_. (b) Similar to Fig. 6a, but with a focus on the specific time range of [0, 0.6 years]. The numerical solution is obtained by solving Eqs. (4)-(7) numerically with initial conditions (8). The approximate solution is determined using Eq. (21). The presented scenario assumes dysfunctional protein degradation machinery with *T*_1/ 2,*D*_ →∞, *T*_1/ 2,*M*_ →∞, and *T*_1/2,*B*_ →∞. The case with *k*_1_ = 10^−6^ s^-1^, *k_M_* = 10^−4^ s^-1^, and *θ*_1/2,*B*_ =10^7^ s. Note that the approximate solution does not depend on *k*_2_.

**Fig. 7.**
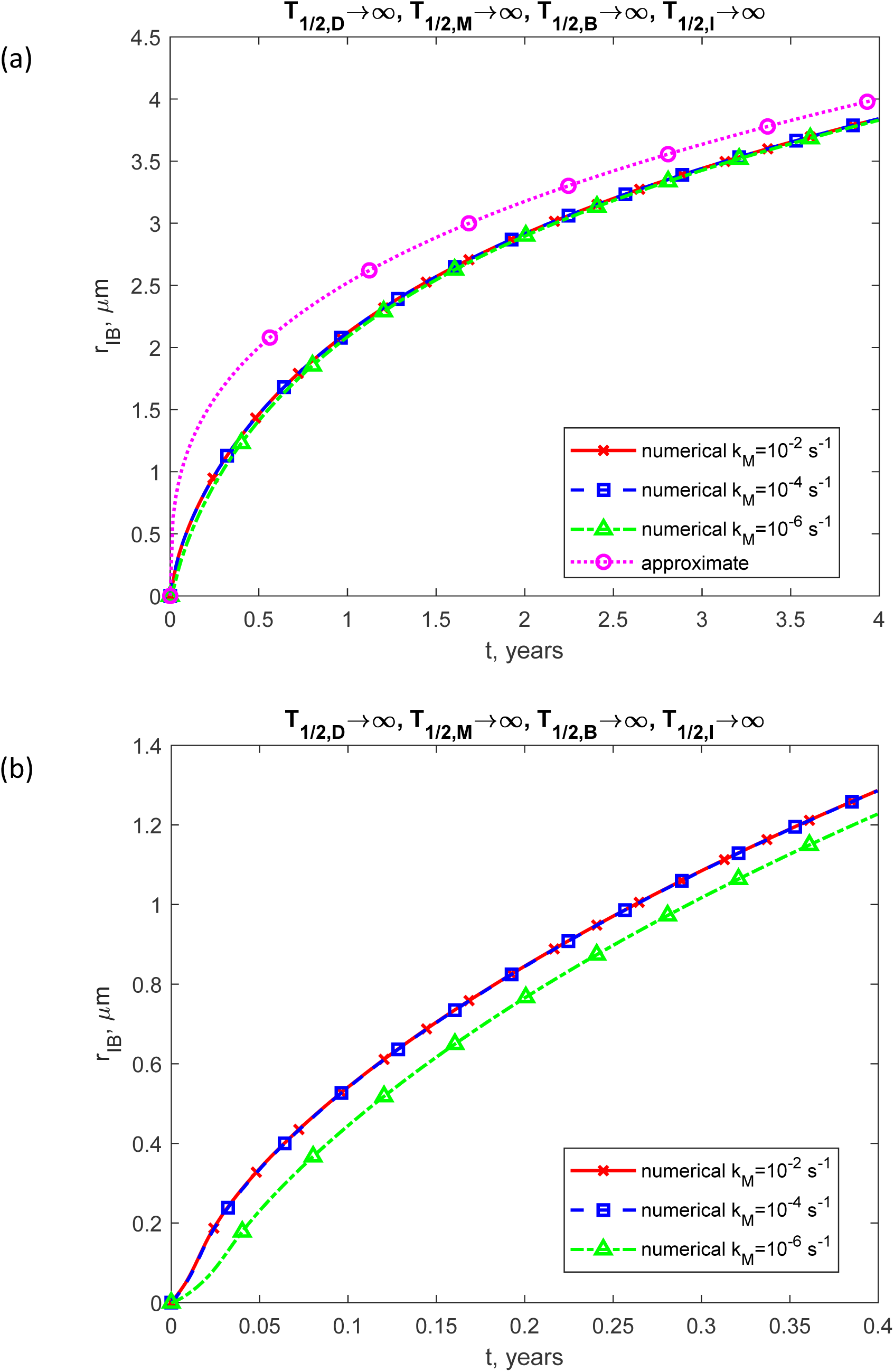
(a) The radius of the growing inclusion body as a function of time (in years). The results are displayed for three values of *k_M_*. (b) Similar to Fig. 7a, but with a focus on the specific time range of [0, 0.4 years]. The approximate solution is determined using Eq. (28). The presented scenario assumes dysfunctional protein degradation machinery with *T*_1/ 2,*D*_ →∞, *T*_1/ 2,*M*_ →∞, and *T*_1/ 2,*B*_ →∞. The case with *k* = 10^−6^ s^-1^, *k* = 10^−4^ s^-1^, and *θ* =10^7^ s.

**Fig. 8.**
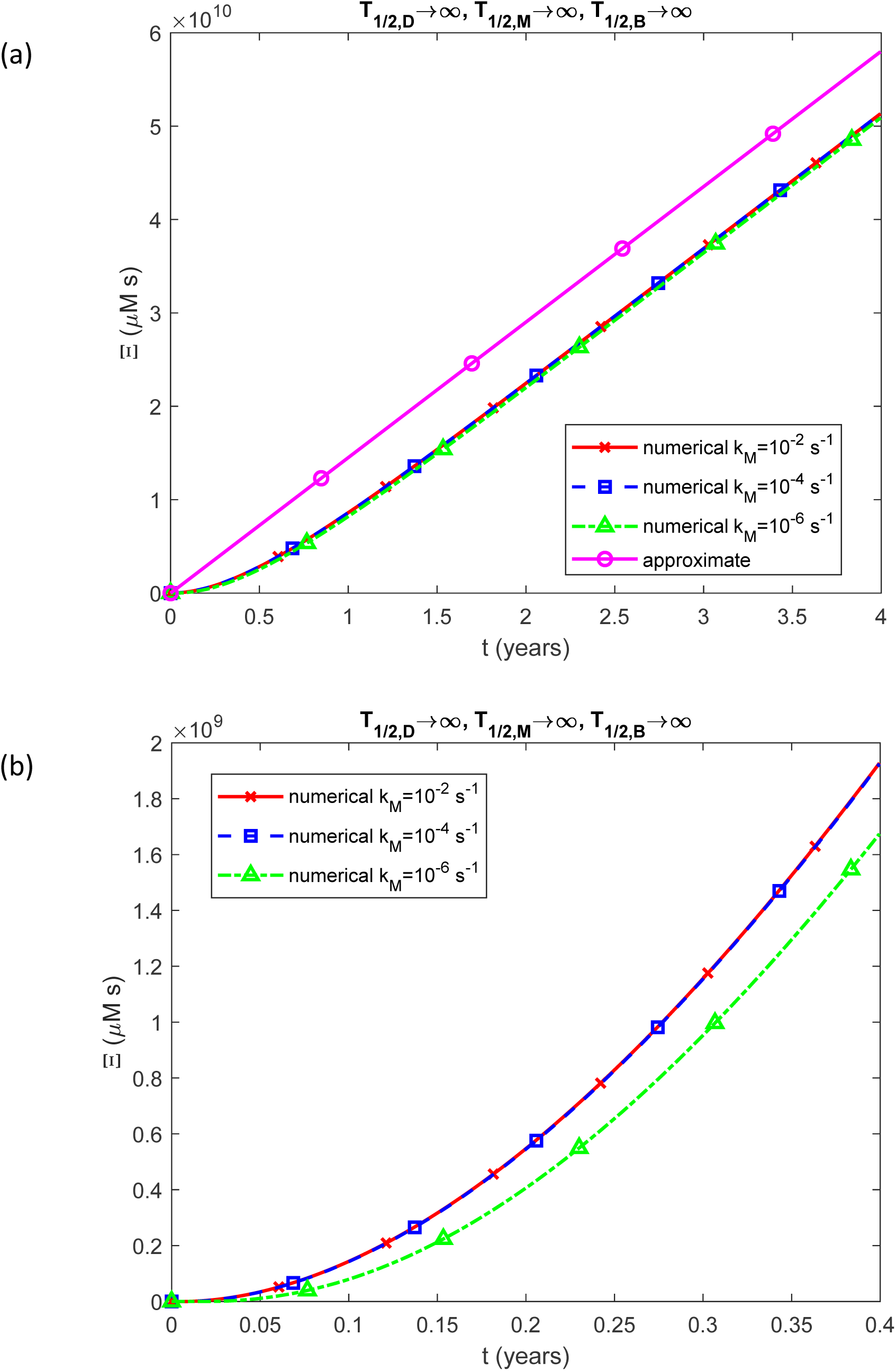
(a) Accumulated neurotoxicity of pathological TDP-43 oligomers, Ξ, as a function of time (in years). Results are shown for three different values of *k_M_*. (b) Similar to Fig. 8a, but with a focus on the specific time range of [0, 0.4 years]. The numerical solution is obtained by solving Eqs. (4)-(7) numerically with initial conditions (8). The approximate solution is determined using Eq. (21). The presented scenario assumes dysfunctional protein degradation machinery with *T*_1/ 2,*D*_ →∞, *T*_1/ 2,*M*_ →∞, and *T*_1/2,*B*_ →∞. The case with *k*_1_ = 10^−6^ s^-1^, *k_M_* = 10^−4^ s^-1^, and *θ*_1/2,*B*_ =10^7^ s. Note that the approximate solution does not depend on *k_M_*.

**Fig. 9.**
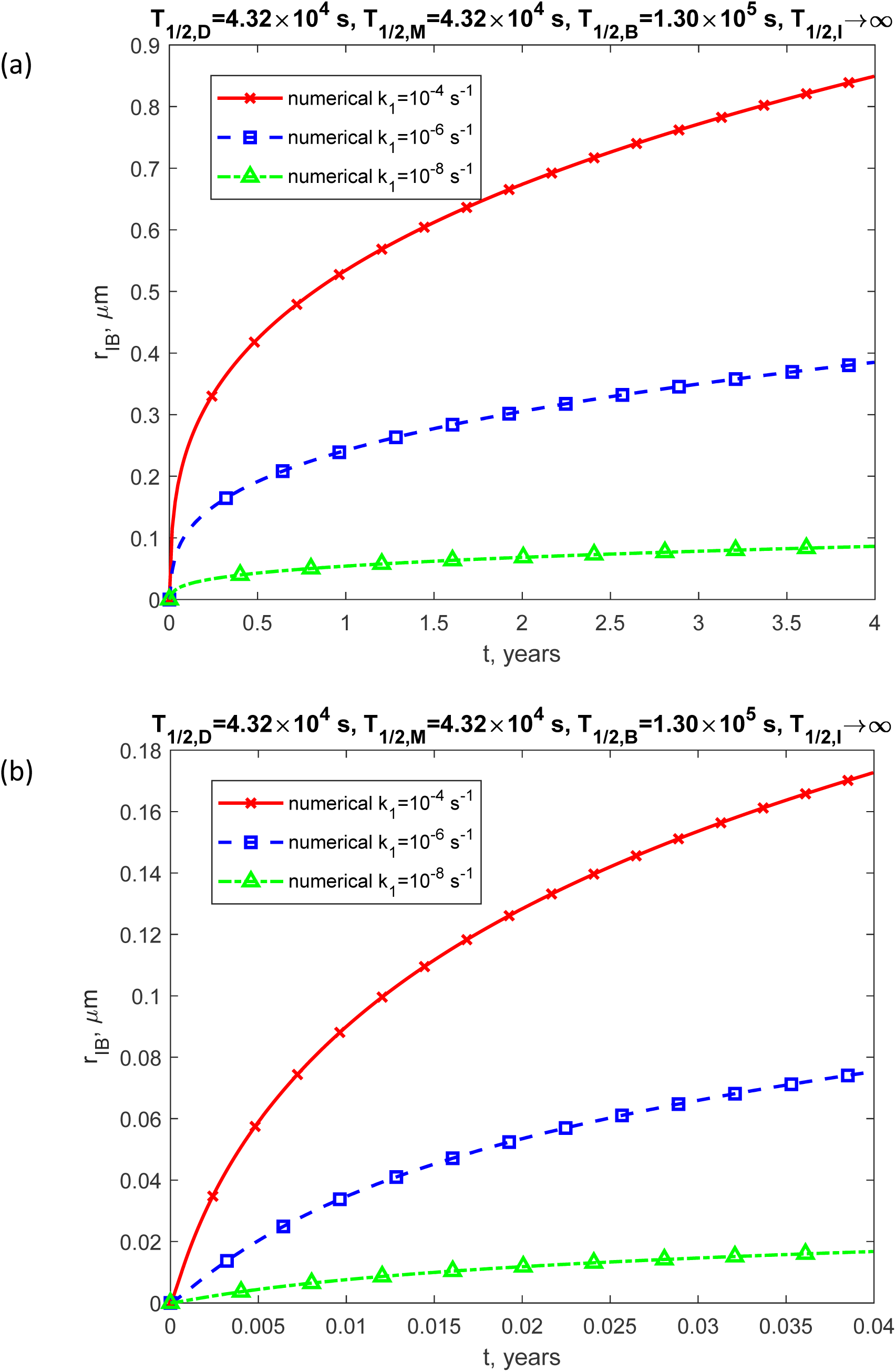
(a) The radius of the growing inclusion body versus time (in years). The results are shown for three values of *k*_1_. (b) Similar to Fig. 9a, but with a focus on the specific time range of [0, 0.04 years]. The presented scenario assumes physiologically relevant values for the half-lives of TDP-43 dimers, monomers, and free misfolded oligomers: *T*_1/2,*D*_ = 4.32×10^4^ s^-1^, *T*_1/2,*M*_ = 4.32×10^4^ s, *T*_1/2,*B*_ =1.30×10^5^ s, and *T*_1/ 2,*I*_ →∞. The case with *k*_2_ = 10^−6^ μM s^-1^, *k* = 10^-4^ s^-1^, and *θ* =10^-4^ s^-1^.

**Fig. 10.**
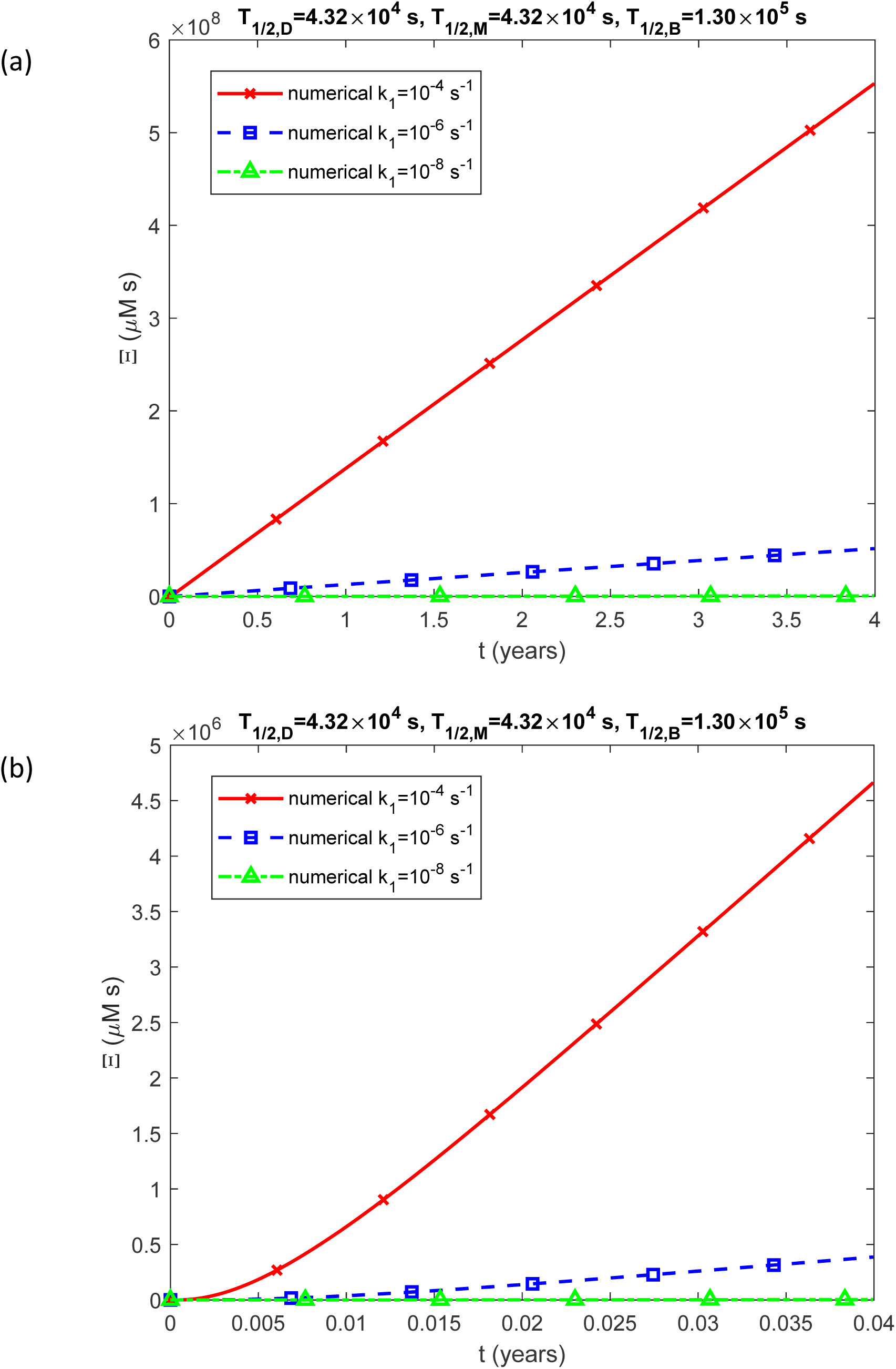
(a) Accumulated neurotoxicity of pathological TDP-43 oligomers, Ξ, as a function of time (in years). Results are shown for three different values of *k*_1_. (b) Similar to Fig. 10a, but with a focus on the specific time range of [0, 0.04 years]. The numerical solution is obtained by solving Eqs. (4)-(7) numerically with initial conditions (8). The presented scenario assumes physiologically relevant values for the half-lives of TDP-43 dimers, monomers, and free misfolded oligomers: *T*_1/2,*M*_ = 4.32×10^4^ s, *T*_1/2,*B*_ = 4.32×10^4^ s, and *T*_2_ =1.30×10^5^ s. The case with *k* = 10^−6^ μM^-1^ s^-1^, *k* = 10^−4^ s^-1^, and *θ_M_* =10^7^ s.

Among all the model parameters listed in Table 2, the author had the least certainty regarding the values of the kinetic constants *k*_1_, *k*_2_, and *k_M_*. An investigation was conducted to assess the consistency of the model’s predictions, focusing on two key relationships: the direct proportionality of the TDP-43 inclusion body radius to the cube root of time, as described in Eq. (28), and the linear increase in accumulated neurotoxicity over time, as described in Eq. (21). This analysis was performed across a broad range of kinetic constant values.

Since inclusion bodies composed of misfolded TDP-43 are highly resistant to degradation (Ling et al. 2010), the half-life of TDP-43 oligomers deposited within them is assumed to be infinite (i.e. *T*_1/ 2,*I*_ →∞) in all figures. Figs. 2-8 are calculated for the scenario of dysfunctional protein degradation machinery, *T*_1/ 2,*D*_ →∞, *T*_1/ 2,*M*_ →∞, and *T*_1/ 2,*B*_ →∞. These figures show the effects of kinetic constants *k*_1_, *k*_2_, and *k_M_*, respectively.

Fig. 2 shows that for *t* > 0.2 years the inclusion body’s radius, *r_IB_*, remains unaffected by the kinetic constant, *k*_1_, that governs the conversion of TDP-43 monomers into a misfolded, aggregating form by nucleation. This is consistent with the approximate solution given by Eq. (28), indicating that the radius is insensitive to the kinetic constants. However, the validity of the approximate solution is limited to extended timeframes (for the parameter values used in computing Fig. 2, it is *t* > 0.2 years). Fig. 2 also shows that numerical solutions are close to the approximate solution given by Eq. (28), indicating that the inclusion body’s radius grows approximately in proportion to *t*^1/3^. The observed discrepancy between numerical and approximate solutions for *r_IB_* (Fig. 2) stems from variations between the numerical and approximate solutions for the concentration of free misfolded TDP-43 oligomers, [*B*] (Fig. S3). These initial differences influence the predicted deposition rates of free misfolded oligomers into inclusion bodies, leading to a persistent deviation in their accumulated concentrations over time, [*I*] (Fig. S4). Consequently, this results in a slight but systematic difference between the numerical and approximate solutions for the inclusion body radius, *r_IB_* (Fig. 2). The approximate solution shown in Fig. 2a predicts larger values of *r_IB_* than the numerical solution because it yields higher concentrations of [*B*] and [*I*] than the numerical solution, as illustrated in Figs. S3a and S4a.

Fig. 3 demonstrates that for *t* > 0.12 years, the accumulated neurotoxicity, Ξ, remains unaffected by the kinetic constant *k*_1_, which governs the nucleation-driven conversion of TDP-43 monomers into misfolded, aggregation-prone oligomers. This aligns with the approximate solution derived in Eq. (21), which suggests that accumulated neurotoxicity is independent of kinetic parameters over extended timeframes. However, the validity of this approximation is constrained to later stages of disease progression (for the parameter values used in Fig. 3, this corresponds to *t* > 0.12 years). Additionally, Fig. 3 shows that the numerical results closely match the approximate solution given by Eq. (21), indicating that accumulated neurotoxicity increases approximately proportionally to time. The slight systematic deviation between the numerical and approximate solutions for Ξ (Fig. 3) can be attributed to differences between the numerical and approximate solutions for the concentrations of free misfolded TDP-43 oligomers, [*B*], at the early stages of the process (Fig. S3).

The data depicting Ξ(*t*) in Fig. 3a and *r_IB_* (*t*) in Fig. 2a have been combined to generate Fig. 4, which illustrates the relationship between accumulated neurotoxicity and the inclusion body radius, Ξ= *f* (*r_IB_*). In ALS, larger TDP-43 inclusion bodies are often correlated with more severe disease progression and poorer prognosis (Jo et al. 2020). If the largest plaque sizes correspond to the most advanced disease stages and neuronal death occurs when plaques reach a critical size, the neurotoxicity threshold at which neurons die, Ξ, can be estimated from Fig. 5 to be approximately 5×10^10^ µMꞏs, which corresponds to a plaque radius of around 3.5 µm. The approximate solution for Ξ= *f* (*r_IB_*) is derived using Eqs. (21) and (28):

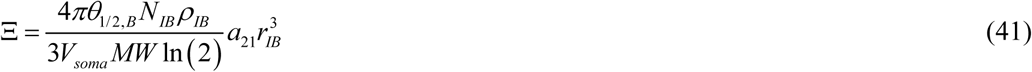

As shown in Fig. 4, the agreement between the numerical and approximate solutions is excellent.

For different values of *k*_2_, which represent the rate of autocatalytic aggregation of TDP-43 monomers, the numerical solution for the inclusion body radius, *r_IB_*, gradually approaches the approximate solution given by Eq. (28) as time progresses (Fig. 5). The discrepancy between the numerical and approximate solutions for *r_IB_* arises from differences in concentrations of free misfolded oligomers, [*B*], predicted by numerical and approximate solutions during the first 2.5 years of the TDP-43 aggregation process (Fig. S7a). This variation leads to differing amounts of oligomers being deposited into inclusion bodies, explaining the shift between numerical and approximate solutions in Fig. 5a. Fig. 6 illustrates that for *t* > 0.6 years the accumulated neurotoxicity, Ξ, is not influenced by the kinetic constant *k*_2_. This observation is consistent with the approximate solution given by Eq. (21), indicating that over long periods, accumulated neurotoxicity remains independent of kinetic constants.

The concentrations of TDP-43 dimers, monomers, and free misfolded oligomers converge to those given by the approximate solution (Figs. S5-S8). Note that as *t* →∞, [*M*] converges to different asymptotic values, each depending on the value of *k*_2_, as described in Eq. (17) (Fig. S6). The concentration of misfolded oligomers deposited into inclusion bodies reaches a linearly increasing curve with time (Fig. S8a), attributed to the constant supply of TDP-43 dimers at a rate of *q_D_*.

Testing the numerical solution for various values of *k_M_*, characterizing the rate of conversion of TDP-43 dimers into monomers, revealed that the numerical solution for *r_IB_* converges to the cube root of time dependence, as predicted by the approximate solution in Eq. (28). The rate of convergence depends on the value of *k_M_* (see Fig. 7), with the slowest convergence observed for the smallest tested value of 10s^-1^.

At longer timescales, the accumulated neurotoxicity, Ξ, increases linearly with time, consistent with the approximate solution presented in Eq. (21) (Fig. 8a). In contrast, over shorter time scales, Ξ appears to exhibit a quadratic dependence on time (Fig. 8b). This aligns with the approximate solution for the neurotoxicity of amyloid β oligomers derived by Kuznetsov (2025a), which is applicable to shorter timeframes than the solution presented in this paper. That solution predicts that accumulated neurotoxicity increases quadratically with time.

The concentration of TDP-43 dimers, [*D*] (Fig. S9), is influenced by *k_M_*, as it is inversely proportional to *k_M_* for a constant value of *q_D_* (see Eq. (14)). Numerical solutions converge to the approximate analytical solutions predicted by Eq. (14), resulting in three distinct curves corresponding to different values of *k_M_* (Fig. S9). In contrast, the concentrations of monomers, [*M*], and free misfolded oligomers, [*B*], converge to the analytical solutions given by Eqs. (17) and (16), respectively, which are independent of the value of *k_M_* (Figs. S10 and S11). The concentration of misfolded TDP-43 oligomers deposited into inclusion bodies, [*I*], exhibits a linear dependence on time at longer timescales (Fig. S12a). A slight discrepancy between the numerical solution and the approximate solution given by Eq. (20) (Fig. S12a) arises from differences in the concentrations of free oligomers, [*B*], predicted by the numerical and analytical solutions during the initial 2.5 years of the process (Fig. S11a).

Figs. 9-14 are computed for physiologically relevant half-lives of TDP-43 dimers, monomers, and free misfolded oligomers ( *T*_1/2,*D*_ = 4.32×10^4^ s, *T*_1/2,*M*_ = 4.32×10^4^ s, and *T*_1/2,*B*_ =1.30×10^5^ s), indicating functional proteolytic machinery. These figures show the effects of kinetic constants *k*_1_, *k*_2_, and *k_M_*, respectively. The approximate solution is not plotted, as it is exclusively applicable to the scenario involving infinite half-lives of TDP-43 dimers, monomers, and free misfolded oligomers.

In contrast to the case with infinite half-lives, the curves representing the radii of inclusion bodies do not converge to a single trajectory over time. Instead, for different values of the kinetic constants ( *k*_1_, *k*_2_, and *k_M_*), the curves differ, depending on the balance between the production of TDP-43 protomers and their degradation due to their finite half-lives (compare Fig. 9 with Fig. 2, Fig. 11 with Fig. 5, and Fig. 13 with Fig. 7). For finite half-lives, the final size of inclusion bodies after four years of growth is significantly smaller than that for infinite half-lives. For instance, in Fig. 9a, with *k* =10^−4^ s^-1^, the inclusion body reaches a size of 0.85 µm after four years, whereas in Fig. 2a, it grows to 4 µm.

**Fig. 11.**
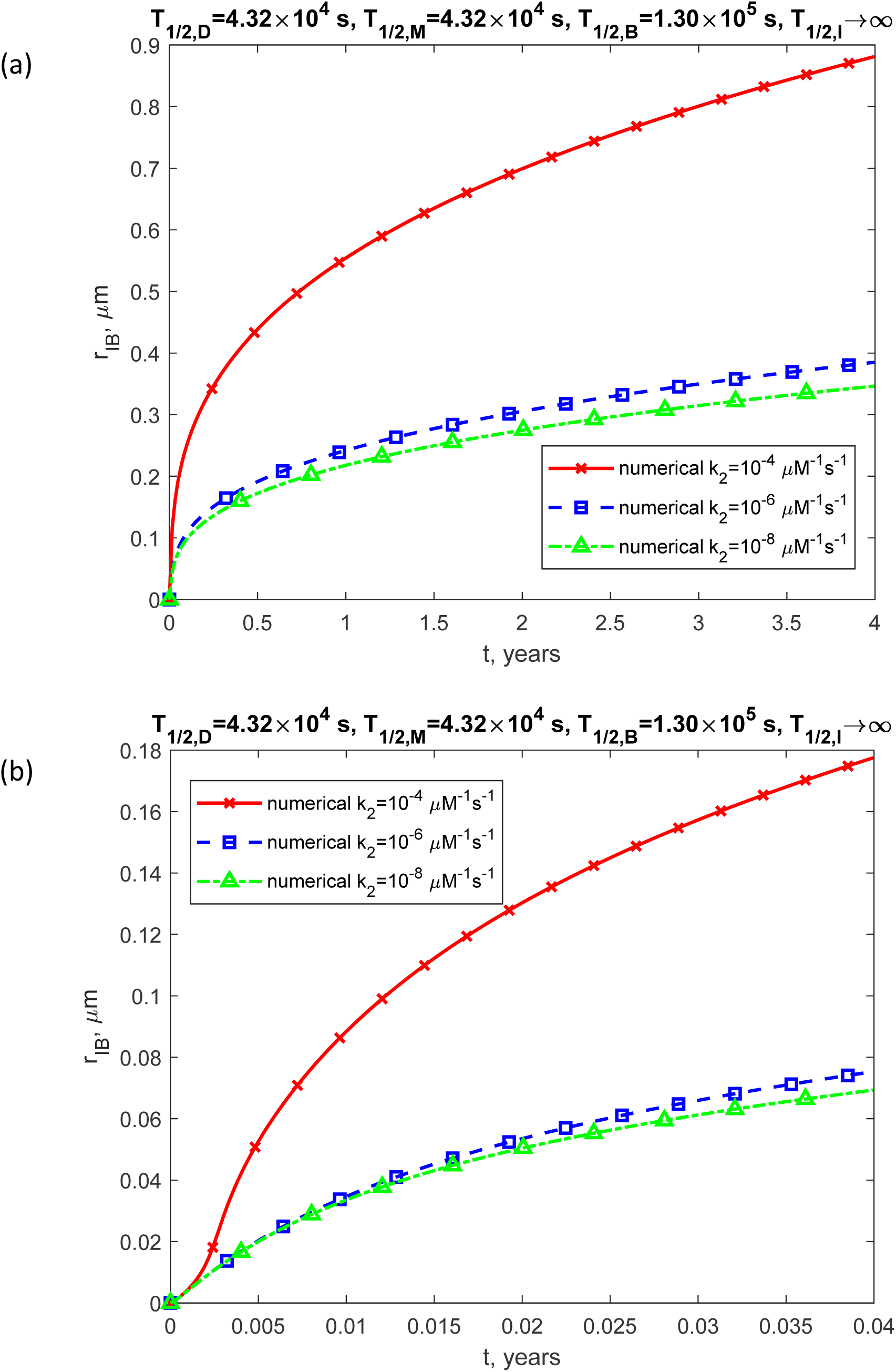
(a) The radius of the growing inclusion body as a function of time (in years). The results are displayed for three values of *k*_2_. (b) Similar to Fig. 11a, but with a focus on the specific time range of [0, 0.04 years]. The presented scenario assumes physiologically relevant values for the half-lives of TDP-43 dimers, monomers, and free misfolded oligomers: *T* = 4.32×10^4^ s, *T*_1/2,*M*_ = 4.32×10^4^ s, *T*_1/2,*B*_ =1.30×10^5^ s, and *T*_1/2,*I*_ →∞. The case with *k*_1_ = 10^−6^ s^-1^, *k_M_* = 10^−4^ s^-1^, and *θ*_1/2,*B*_ =10^7^ s.

**Fig. 12.**
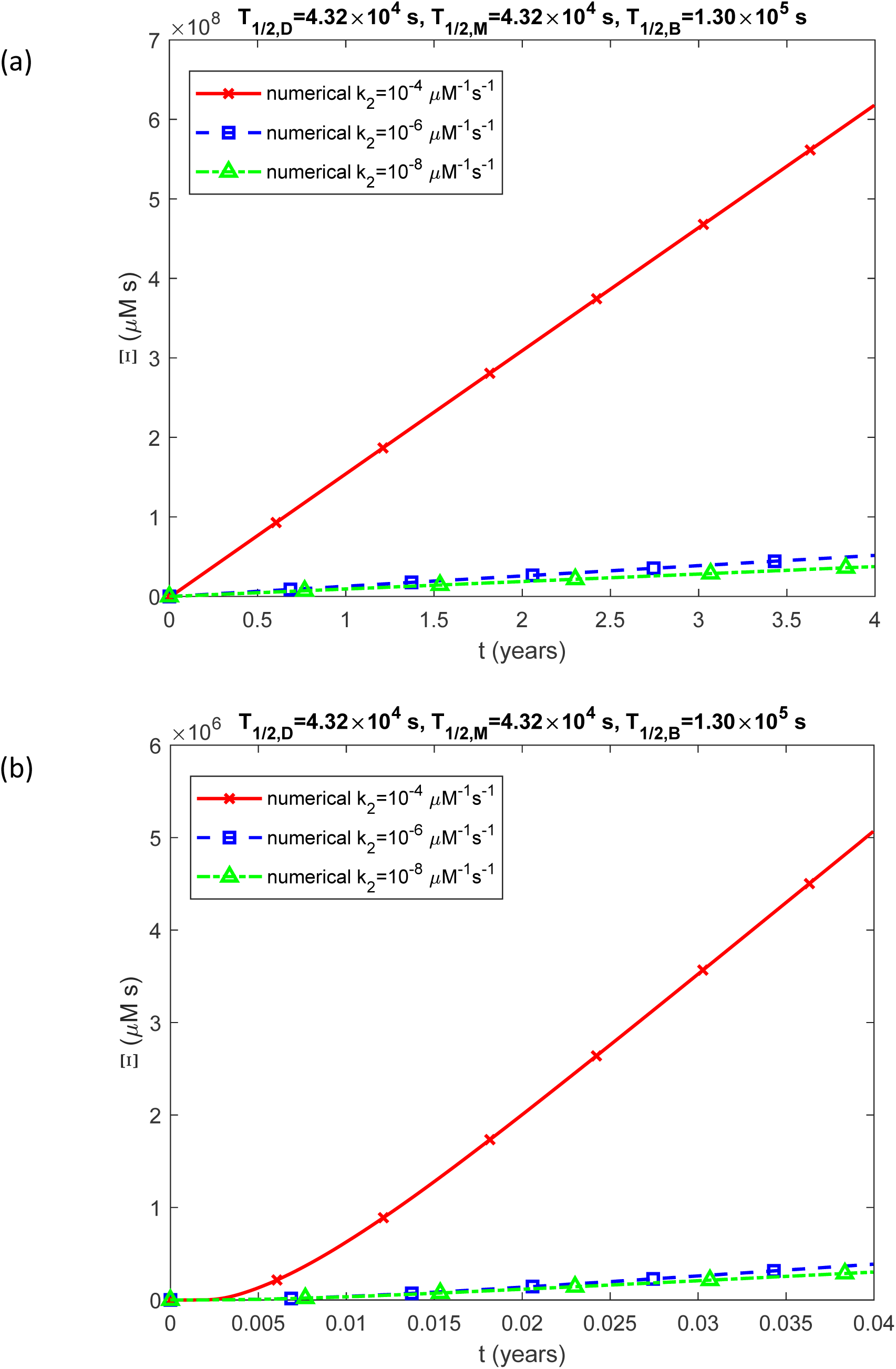
(a) Accumulated neurotoxicity of pathological TDP-43 oligomers, Ξ, as a function of time (in years). Results are shown for three different values of *k*_2_. (b) Similar to Fig. 12a, but with a focus on the specific time range of [0, 0.04 years]. The numerical solution is obtained by solving Eqs. (4)-(7) numerically with initial conditions (8). The presented scenario assumes physiologically relevant values for the half-lives of TDP-43 dimers, monomers, and free misfolded oligomers: *T*_1/2,*M*_ = 4.32×10^4^ s, *T*_1/2,*B*_ = 4.32×10^4^ s, and *T* =1.30×10^5^ s. The case with *k*_1_ = 10^−6^ s^-1^, *k_M_* = 10^−4^ s^-1^, and *θ*_1/2,*B*_ =10^7^ s.

**Fig. 13.**
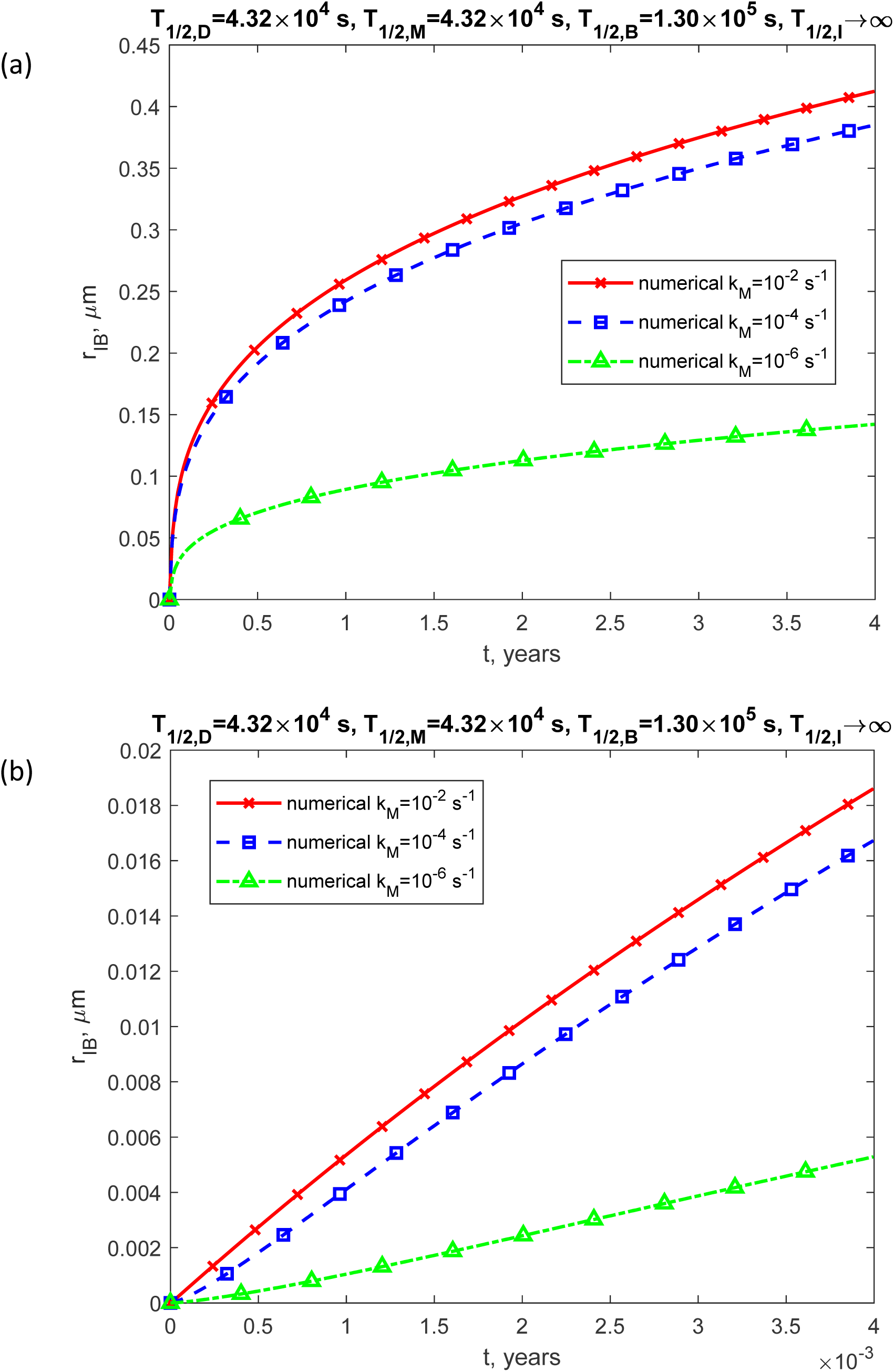
(a) The radius of the growing inclusion body as a function of time (in years). The results are shown for three different values of *k_M_*. (b) Similar to Fig. 13a, but with a focus on the specific time range of [0, 0.004 years]. The presented scenario assumes physiologically relevant values for the half-lives of TDP-43 dimers, monomers, and free misfolded oligomers: *T*_1/2,*D*_ = 4.32×10^4^ s, *T*_1/2,*M*_ = 4.32×10^4^ s, *T*_1/2,*B*_ =1.30×10^5^ s, and *T*_1/2,*I*_ →∞. The case with *k*_1_ = 10^−6^ s^-1^, *k*_2_ = 10^−6^ μM^-1^ s^-1^, and *θ*_1/2,*B*_ =10^7^ s.

Similarly, in the case of finite half-lives, the curves representing accumulated neurotoxicity do not converge to a single curve over time (compare Fig. 10 with Fig. 3, Fig. 12 with Fig. 6, and Fig. 14 with Fig. 8). However, even with finite half-lives, accumulated neurotoxicity increases linearly with time (Figs. 10a, 12a, and 14a). This relationship arises from the constant supply of TDP-43 dimers, *q_D_*, in the model, as described by Eq. (4).

**Fig. 14.**
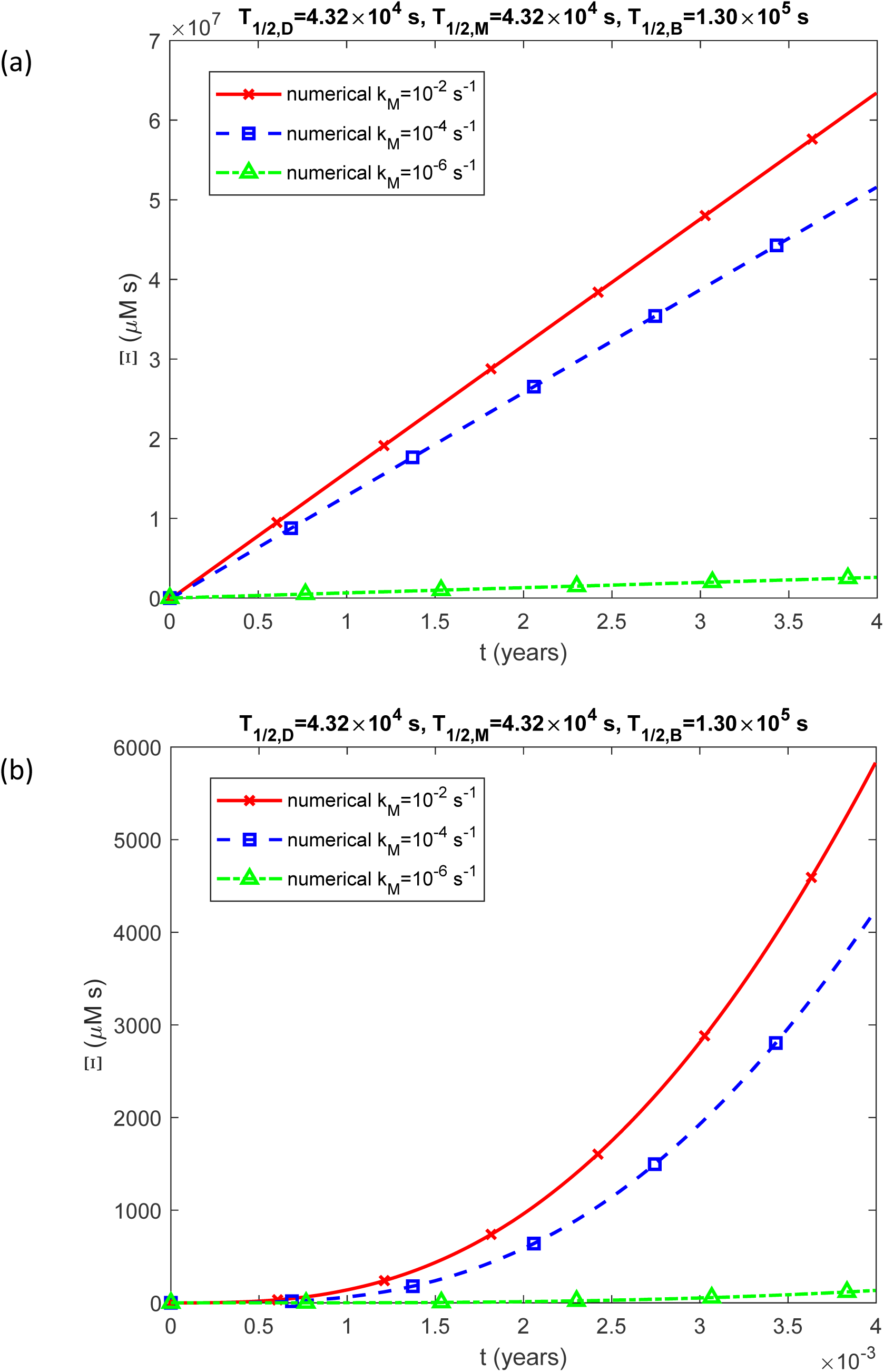
(a) Accumulated neurotoxicity of pathological TDP-43 oligomers, Ξ, as a function of time (in years). Results are shown for three different values of *k_M_*. (b) Similar to Fig. 14a, but with a focus on the specific time range of [0, 0.004 years]. The numerical solution is obtained by solving Eqs. (4)-(7) numerically with initial conditions (8). The presented scenario assumes physiologically relevant values for the half-lives of TDP-43 dimers, monomers, and free misfolded oligomers: *T*_1/2,*M*_ = 4.32×10^4^ s, *T* = 4.32×10^4^ s, and *T*_1/2,*B*_ =1.30×10^5^ s. The case with *k*_1_ = 10^−6^ s^-1^, *k*_2_ = 10^−6^ μM^-1^ s^-1^, and *θ* =10^7^ s.

The concentration of TDP-43 dimers, [*D*], remains independent of the kinetic constants *k*_1_ and *k*_2_ (Figs. S13 and S17) but varies with the value of *k_M_* (Fig. S21). In contrast, the concentration of TDP-43 monomers, [*M*], is influenced by *k*_1_, *k*_2_, and *k_M_* (Figs. S14, S18, and S22). Similarly, the concentration of free misfolded TDP-43 oligomers, [*B*], depends on *k*_1_, *k*_2_, and *k_M_* (Figs. S15, S19, and S23). The concentration of misfolded TDP-43 oligomers deposited into inclusion bodies, [*I*], is also affected by *k*_1_, *k*_2_, and *k_M_* (Figs. S16, S20, and S24). Notably, [*I*] exhibits a linear dependence on time.

**Fig. 15.**
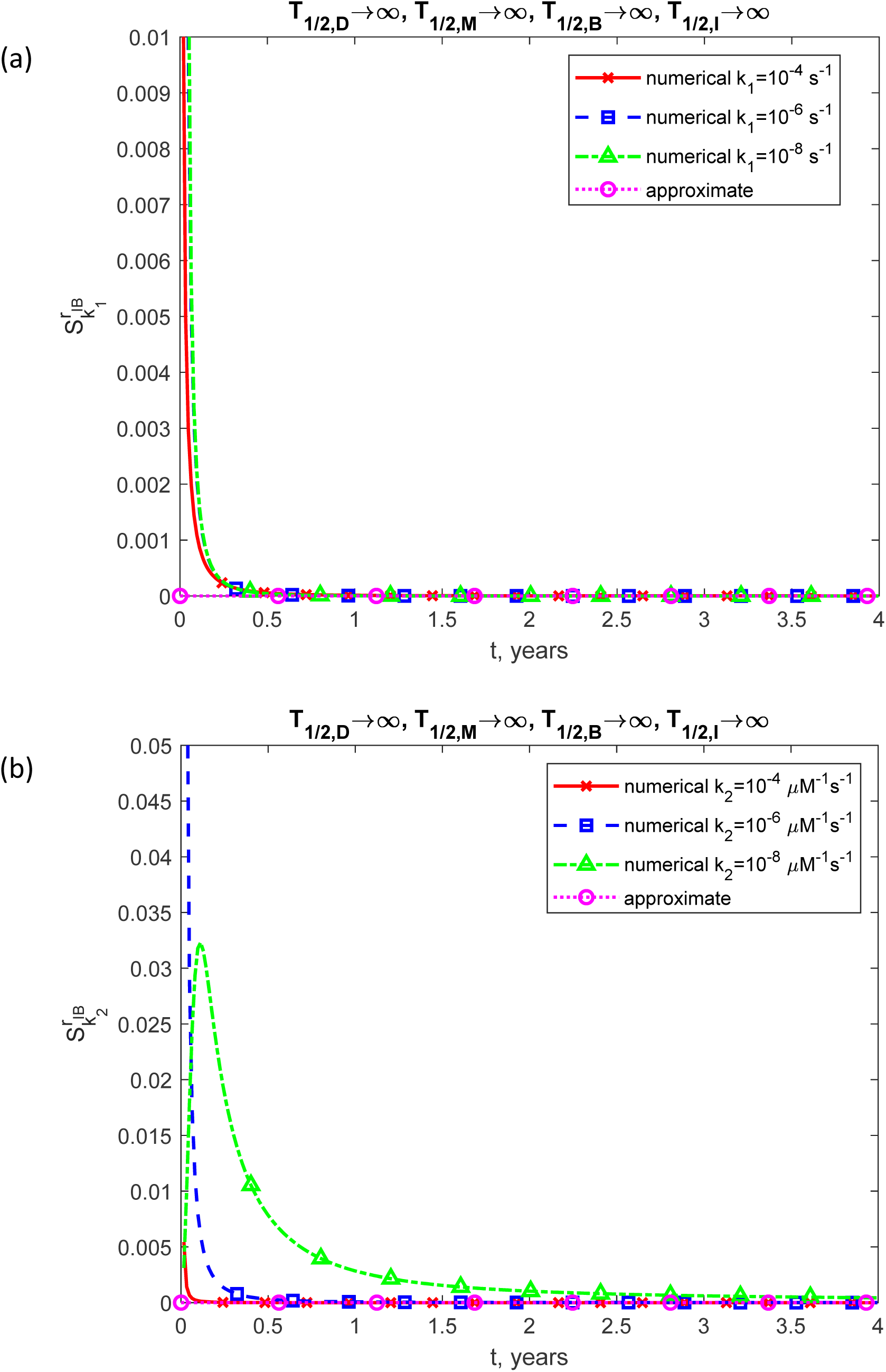
The dimensionless sensitivity of the inclusion body’s radius, *r_IB_*, to: (a) the rate constant associated with the first pseudoelementary step in the F-W model, describing the formation of free misfolded oligomers through nucleation, *k*_1_ (calculated for *k* = 10^−6^ μM^-1^ s^-1^, *k_M_* = 10^−4^ s^-1^, *θ*_1/2,*B*_ =10^7^ s for three distinct values of *k*_1_); (b) the rate constant associated with the second pseudoelementary step in the F-W model, describing the formation of free misfolded oligomers through autocatalytic growth, *k*_2_ (calculated for *k*_1_ = 10^−6^ s^-1^, *k_M_* = 10^−4^ s^-1^, *θ*_1/2,*B*_ =10^7^ s for three distinct values of *k*_2_); (c) the rate constant linked to the reaction describing the conversion of TDP-43 dimers into monomers, *k_M_* (calculated for *k*_1_ = 10^−6^ s^-1^, *k*_2_ = 10^−6^ μM^-1^ s^-1^, *θ*_1/2,*B*_ =10^7^ s for three distinct values of *k_M_*).

**Fig. 16.**
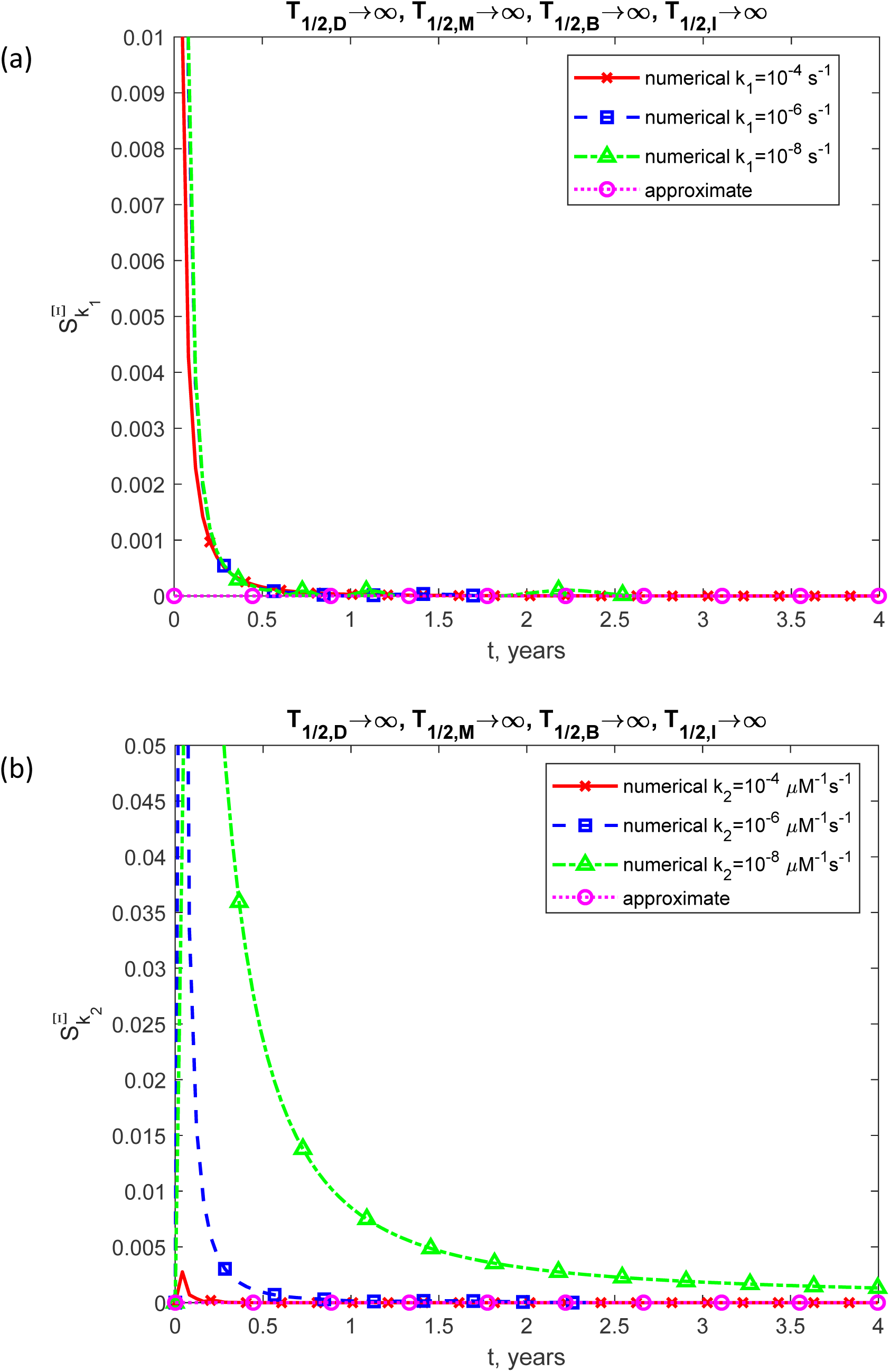
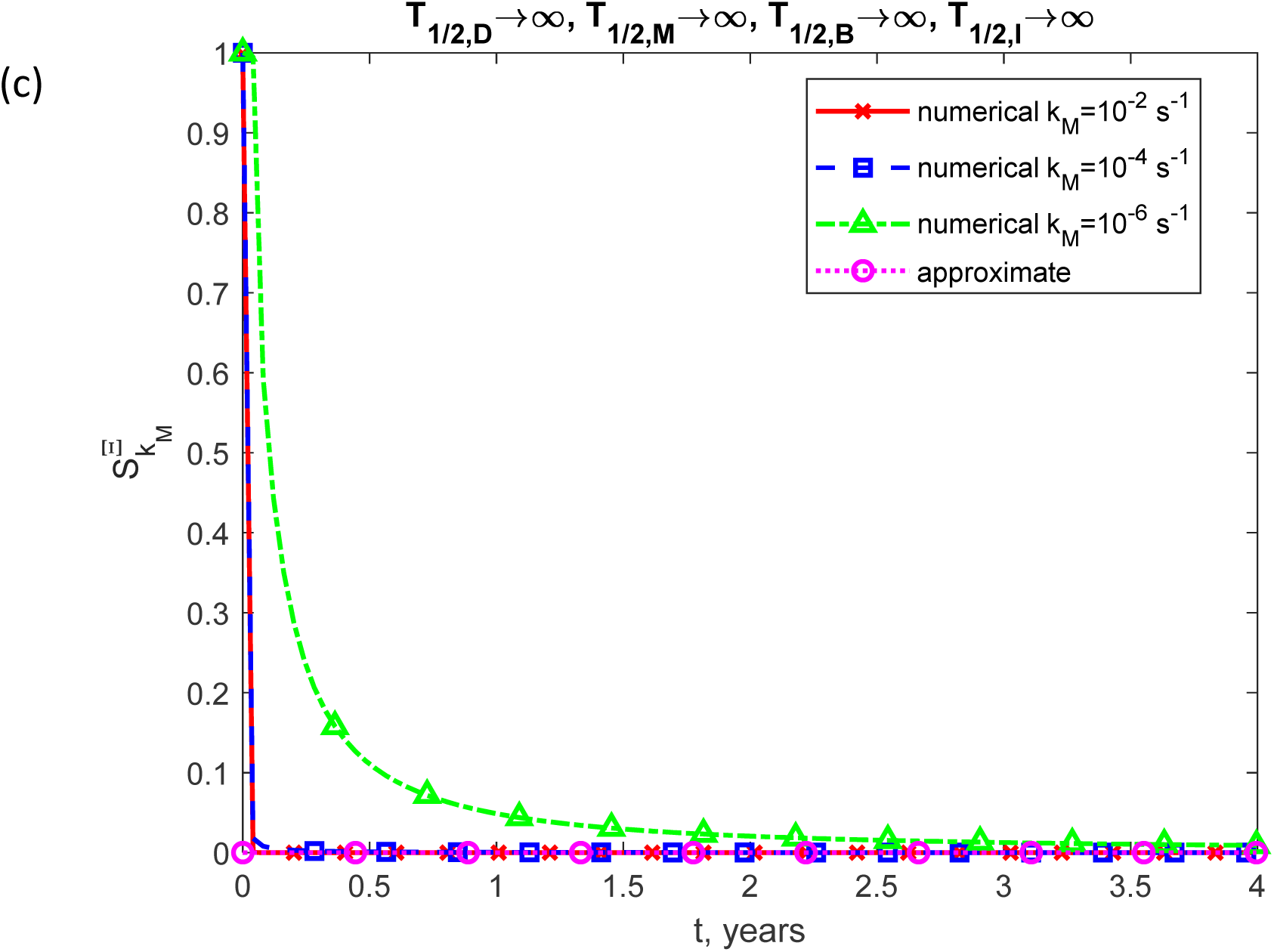
The dimensionless sensitivity of the accumulated neurotoxicity of pathological TDP-43 oligomers, Ξ, to: (a) the rate constant associated with the first pseudoelementary step in the F-W model, describing the formation of free misfolded oligomers through nucleation, *k*_1_ (calculated for *k* = 10^−6^ μM^-1^ s^-1^, *k_M_* = 10^−4^ s^-1^, *θ*_1/2,*B*_ =10^7^ s for three distinct values of *k*_1_); (b) the rate constant associated with the second pseudoelementary step in the F-W model, describing the formation of free misfolded oligomers through autocatalytic growth, *k*_2_ (calculated for *k*_1_ = 10^−6^ s^-1^, *k_M_* = 10^−4^ s^-1^, *θ*_1/2,*B*_ =10^7^ s for three distinct values of *k*_2_); (c) the rate constant linked to the reaction describing the conversion of TDP-43 dimers into monomers, *k_M_* (calculated for *k*_1_ = 10^−6^ s^-1^, *k*_2_ = 10^−6^ μM^-1^ s^-1^, *θ*_1/2,*B*_ =10^7^ s for three distinct values of *k_M_*).

Fig. 15 examines the sensitivities of the inclusion body radius, *r_IB_*, to the kinetic constants *k*_1_, *k*_2_, and *k_M_* in a scenario where TDP-43 dimers, monomers, and free misfolded oligomers have an infinite half-life. At short timescales, the sensitivities to *k*_1_, *k*_2_, and *k_M_* are positive, indicating that *r_IB_* increases with increasing values of these kinetic constants. At larger timescales, the sensitivities of *r_IB_* to all three kinetic constants gradually approach zero, consistent with the predictions obtained from the approximate analytical solution (Eq. (28)).

Fig. 16 examines the sensitivities of accumulated neurotoxicity, Ξ, to the kinetic constants *k*_1_, *k*_2_, and *k_M_* in a scenario where TDP-43 dimers, monomers, and free misfolded oligomers have infinite half-lives. At small timescales, the sensitivities to *k*_1_, *k*_2_, and *k_M_* are positive. At larger timescales, the sensitivities of Ξ to all three kinetic constants gradually approach zero, consistent with the predictions of the approximate analytical solution (Eq. (21)).

## 4. Discussion, limitations of the model, and future directions

This paper develops a model for the growth of TDP-43 inclusion bodies and introduces the concept of accumulated neurotoxicity caused by free misfolded TDP-43 oligomers. Numerical and approximate analytical solutions for neurotoxicity as a function of time were obtained. The model predicts the concentrations of TDP-43 dimers, monomers, free misfolded oligomers, and misfolded oligomers deposited into inclusion bodies.

In the case of dysfunctional protein degradation machinery—where protein dimers, monomers, and free misfolded oligomers have infinite half-lives—the model predicts that the inclusion body radius grows proportionally to the cube root of time, while accumulated neurotoxicity increases linearly over time.

Since the kinetic constants governing TDP-43 monomerization, nucleation, and autocatalytic growth are not well-documented in the literature, an extensive analysis was conducted to assess the consistency of model predictions across a broad range of parameter values. The results indicate that the inclusion body radius and accumulated neurotoxicity are influenced by these kinetic constants only at short timescales.

The approximate solution, valid for infinite half-lives of TDP-43 protomers and long timescales, reveals that the concentration of TDP-43 dimers, [*D*], is unaffected by the nucleation and autocatalytic kinetic constants, *k*_1_ and *k*_2_, but depends on the monomerization kinetic constant, *k_M_* (see Eq. (14)). In contrast, the concentration of TDP-43 monomers, [*M*], is independent of *k_M_* but varies with *k*_1_ and *k*_2_ (see Eq. (17)). The concentrations of free misfolded oligomers and those incorporated into inclusion bodies, [*B*] and [*I*], respectively, are independent of *k*_1_, *k*_2_, and *k_M_* (see Eqs. (16) and (20)). The approximate solution also indicates a linear increase in the concentration of deposited TDP-43 oligomers, [*I*], over time, attributed to the continuous synthesis of TDP-43 dimers in the cytoplasm (more precisely, TDP-43 monomers are synthesized and subsequently combined into dimers). The obtained results suggest that stabilization of TDP-43 dimers in the nucleus, which are not prone to aggregation, can be a viable therapeutic approach.

The approximate solution is no longer valid when considering physiologically relevant half-lives for TDP-43 dimers, monomers, and free misfolded oligomers. Unlike the case of infinite half-lives, the curves representing the inclusion body radii and accumulated neurotoxicity do not converge to a single trajectory over time for different values of the kinetic constants. The concentrations of dimers, monomers, and free misfolded oligomers reach distinct constant values, which depend on the values of the kinetic constants. Similarly, the concentrations of deposited misfolded oligomers converge to distinct linear trends over time.

When the half-lives of TDP-43 dimers, monomers, and free misfolded oligomers are assumed to be infinite, the sensitivities of the inclusion body radius to the kinetic constants governing nucleation, autocatalysis, and monomerization ( *k*_1_, *k*_2_, and *k_M_*) are positive at early times. As time progresses, all sensitivities gradually approach zero, indicating that at large timescales, the inclusion body radius becomes independent of these kinetic constants.

When the half-lives of TDP-43 dimers, monomers, and free misfolded oligomers are infinite, the sensitivities of accumulated neurotoxicity to the kinetic constants are initially positive but gradually decline to zero over time. This implies that, at large timescales, accumulated neurotoxicity is no longer affected by the kinetic constants.

The analysis presented here has limitations, primarily due to its reliance on a simplified mechanism of TDP-43 cytoplasmic mislocalization and subsequent aggregation, compounded by the absence of precise values for kinetic constants.

Investigating the biological mechanisms underlying TDP-43 misfolding would be highly valuable. Future studies should concentrate on developing more precise models for TDP-43 oligomerization, covering the entire spectrum of oligomeric species, including trimers, tetramers, and multimers. Furthermore, the processes of TDP-43 phosphorylation and ubiquitination must be integrated into these models. Future models should incorporate simulations of TDP-43 shuttling between the nucleus and cytoplasm, the kinetics of TDP-43 binding to RNAs, and the impact of these processes on TDP-43 aggregation.

Additionally, they should distinguish between physiological and pathological oligomerization. Investigating the potential prion-like spread of pathological TDP-43 species is also a crucial avenue for future exploration. Comparing the model’s predictions with future experimental data will be essential for providing quantitative evidence to either support or refute the proposed theory.

Beyond neuronal cytoplasmic inclusions, neuropathological studies have also documented neuronal intranuclear inclusions and dystrophic neurites in both neurons and certain glial cells (Lee, E. B. et al. 2017). These diverse pathological features are frequently linked to the structural brain atrophy observed in neurodegenerative diseases (Youssef et al. 2024). Given the well-established role of TDP-43 as a nucleocytoplasmic shuttling protein, the current model’s primary focus on cytosolic aggregation represents a limitation. Future extensions of the model should therefore consider incorporating aggregation dynamics across multiple subcellular compartments to better capture the full spectrum of TDP-43 pathology.

While the present model focuses on the aggregation dynamics of canonical TDP-43, it does not currently account for the heterogeneity of TDP-43 species often observed in disease states. Recent investigations have identified specific pathogenic variants, such as the 35-kDa N-terminally truncated species (Met85-TDP-35) arising from an alternative translation initiation site (Xiao et al. 2015). This variant is upregulated under pathological conditions and has been demonstrated to form cytoplasmic aggregates and induce motor neuron death. The exclusion of these alternative isoforms represents a limitation of the current framework. Future extensions of this model should therefore aim to incorporate the coexistence and interaction of multiple toxic TDP-43 species, examining how such variants might modulate the formation kinetics of misfolded oligomers and the subsequent growth of inclusion bodies.

The current model analyzes TDP-43 aggregation in isolation; however, in the context of FTLD, TDP-43 pathology often co-occurs with other proteinopathies. Recent evidence suggests that these co-pathologies may exacerbate neurodegeneration (Youssef et al. 2025). Consequently, limiting the simulation to a single aggregating species simplifies the complex proteostatic environment of the disease. A promising direction for future research is the extension of this mathematical framework to simulate the interaction kinetics between TDP-43 and other aggregating proteins, thereby broadening the model’s applicability to a wider spectrum of neurodegenerative disorders.

Incorporating modern mathematical techniques developed for biological problems—particularly those that offer tools for accounting for the elasticity of domain boundaries (in this case, the neuron)—would greatly enhance future models (Alghamdi et al. 2024; Sher Akbar et al. 2023; Maraj et al. 2023; Ghailan et al. 2023; Akbar et al. 2023; Akram and Akbar 2023; Akbar and Muhammad 2023; Akbar et al. 2024; Noreen and Akhtar 2023).

## Acknowledgment

The author acknowledges the support provided by the National Science Foundation (grant DMS-2451660) and the Alexander von Humboldt Foundation through the Humboldt Research Award.

## Abbreviations

ALS: amyotrophic lateral sclerosis
F-W: Finke-Watzky
FTLD: frontotemporal lobar degeneration
TDP-43: transactive response DNA binding protein of 43 kDa

## Supplemental Materials

### S1. Supplemental tables

**Table S1.**
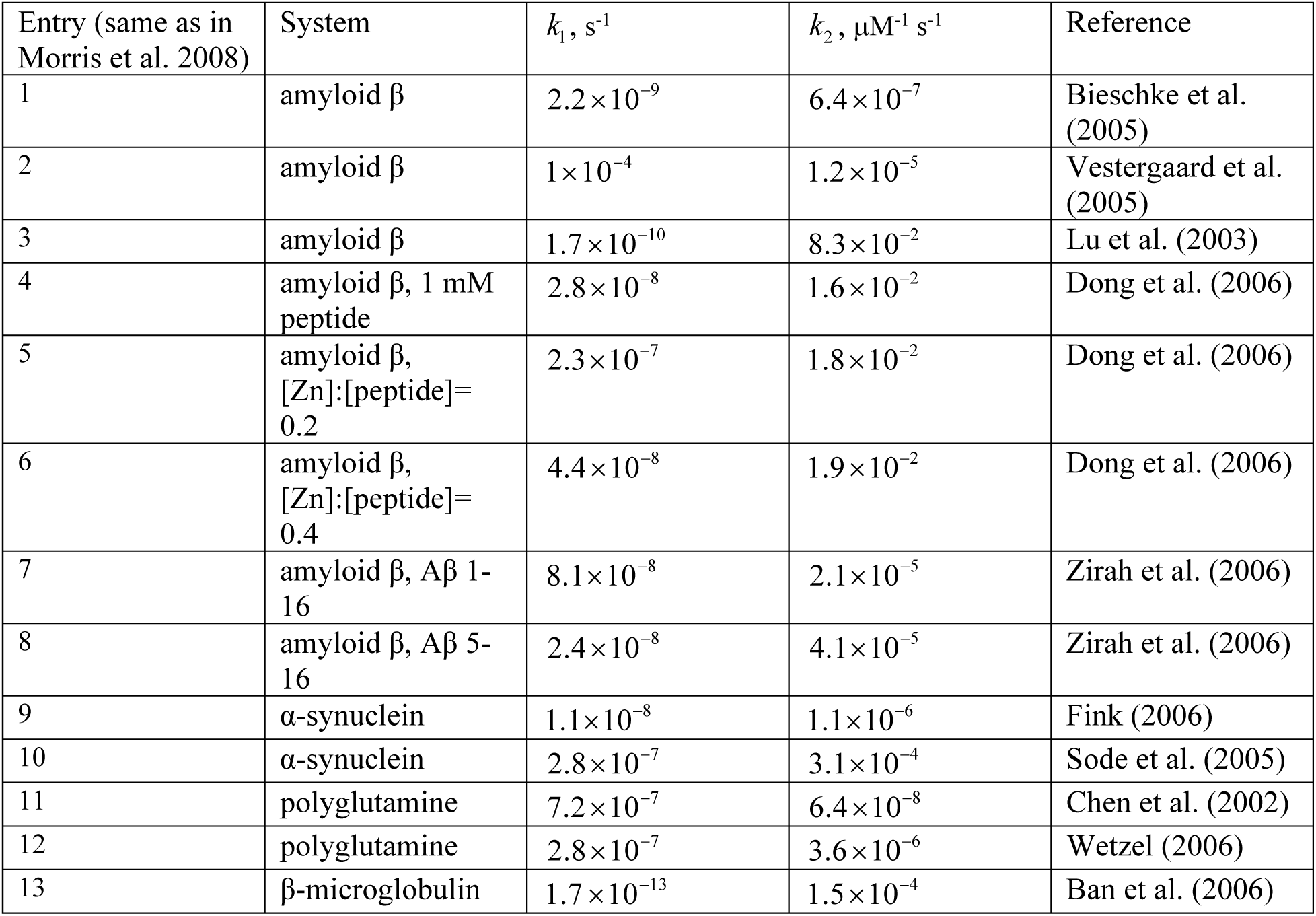
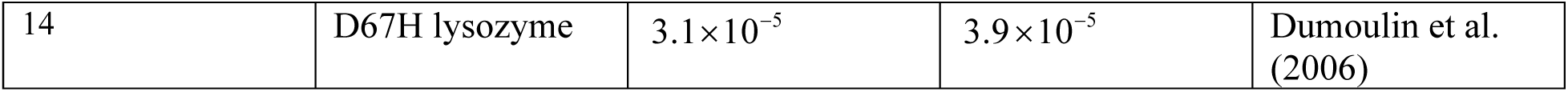
Data extracted from Table 2 of Morris et al. (2008), presenting best-fit values of rate constants, *k*_1_ and *k*_2_, used for simulating the aggregation of diverse proteins by the F-W model, *A*—*^k^*—^1^ →*B*, *A* + *B* —*^k^*—^2^ →2*B* ( *A* denotes monomers and *B* denotes aggregates). The data reported in Morris et al. (2008) have been converted into SI units to establish representative ranges for *k*_1_ and *k*_2_. This was done because values for *k*_1_ and *k*_2_ specific to TDP-43 are not available in the literature.

| Entry (same as in Morris et al. 2008) | System | $k_1, \text{s}^{-1}$ | $k_2, \mu\text{M}^{-1} \text{s}^{-1}$ | Reference |
| --- | --- | --- | --- | --- |
| 1 | amyloid $\beta$ | $2.2 \times 10^{-9}$ | $6.4 \times 10^{-7}$ | Bieschke et al. (2005) |
| 2 | amyloid $\beta$ | $1 \times 10^{-4}$ | $1.2 \times 10^{-5}$ | Vestergaard et al. (2005) |
| 3 | amyloid $\beta$ | $1.7 \times 10^{-10}$ | $8.3 \times 10^{-2}$ | Lu et al. (2003) |
| 4 | amyloid $\beta$, 1 mM peptide | $2.8 \times 10^{-8}$ | $1.6 \times 10^{-2}$ | Dong et al. (2006) |
| 5 | amyloid $\beta$, [Zn]:[peptide]=0.2 | $2.3 \times 10^{-7}$ | $1.8 \times 10^{-2}$ | Dong et al. (2006) |
| 6 | amyloid $\beta$, [Zn]:[peptide]=0.4 | $4.4 \times 10^{-8}$ | $1.9 \times 10^{-2}$ | Dong et al. (2006) |
| 7 | amyloid $\beta$, Aβ 1-16 | $8.1 \times 10^{-8}$ | $2.1 \times 10^{-5}$ | Zirah et al. (2006) |
| 8 | amyloid $\beta$, Aβ 5-16 | $2.4 \times 10^{-8}$ | $4.1 \times 10^{-5}$ | Zirah et al. (2006) |
| 9 | α-synuclein | $1.1 \times 10^{-8}$ | $1.1 \times 10^{-6}$ | Fink (2006) |
| 10 | α-synuclein | $2.8 \times 10^{-7}$ | $3.1 \times 10^{-4}$ | Sode et al. (2005) |
| 11 | polyglutamine | $7.2 \times 10^{-7}$ | $6.4 \times 10^{-8}$ | Chen et al. (2002) |
| 12 | polyglutamine | $2.8 \times 10^{-7}$ | $3.6 \times 10^{-6}$ | Wetzel (2006) |
| 13 | β-microglobulin | $1.7 \times 10^{-13}$ | $1.5 \times 10^{-4}$ | Ban et al. (2006) |
| 14 | D67H lysozyme | $3.1 \times 10^{-5}$ | $3.9 \times 10^{-5}$ | Dumoulin et al. (2006) |

### S2. Supplemental figures

Figs. S1-S12 and Figs. S13-S24 illustrate the time-dependent concentration profiles of TDP-43 dimers [*D*], monomers [*M*], free misfolded oligomers [*B*], and misfolded TDP-43 oligomers deposited into inclusion bodies [*I*] under different conditions. Figs. S1-S12 depict a situation with infinite half-lives of TDP-43 dimers, monomers, and free misfolded oligomers, while Figs. S13-S24 represent a scenario with physiologically relevant half-lives of these proteins. The numerical solution is obtained by solving Eqs. (4)-(7) numerically with initial conditions (8). The approximate analytical solution depicted in Figs. S1-S12 is computed utilizing Eqs. (14), (16), (17), and (20).

**Fig. S1.**
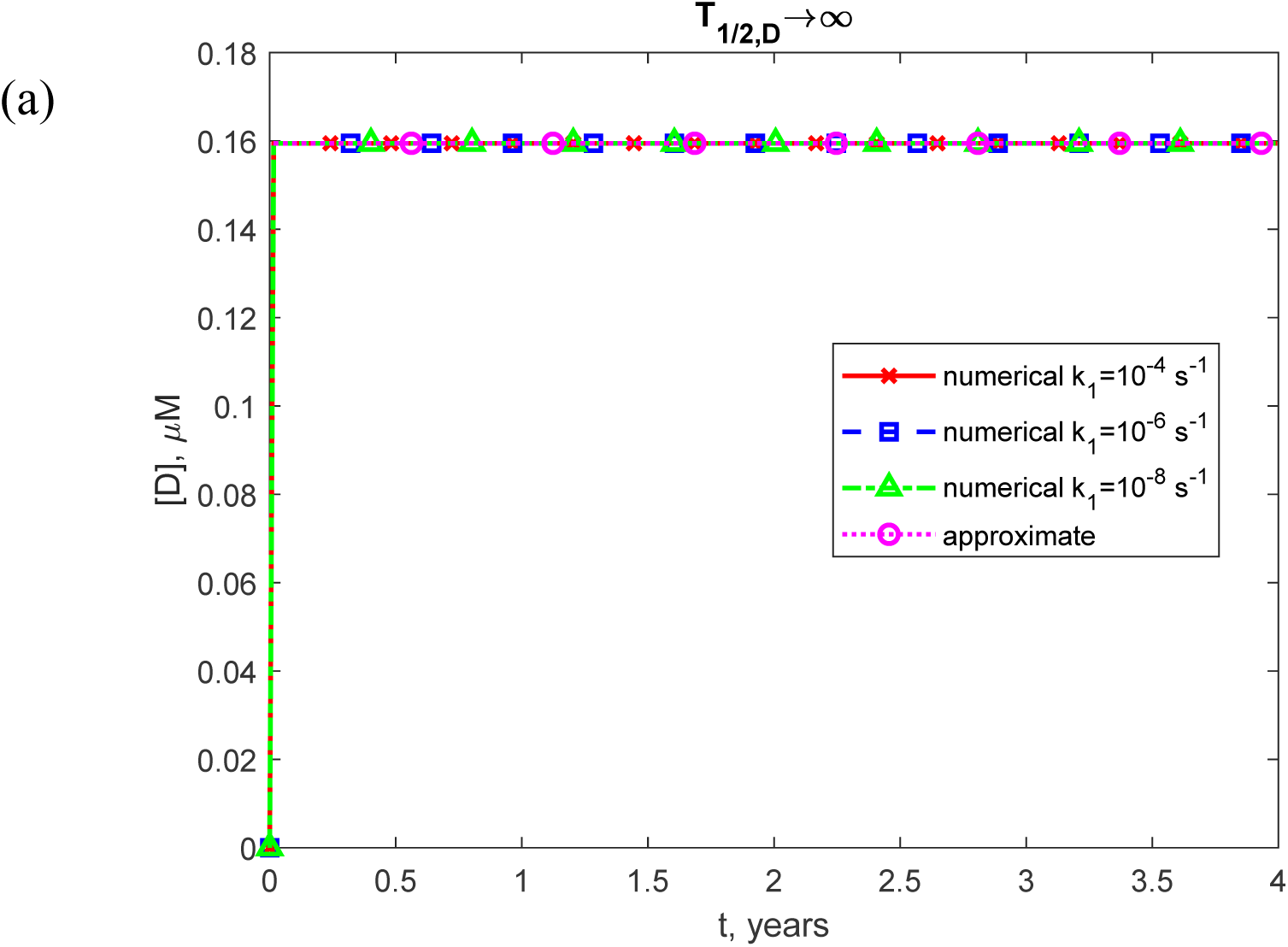
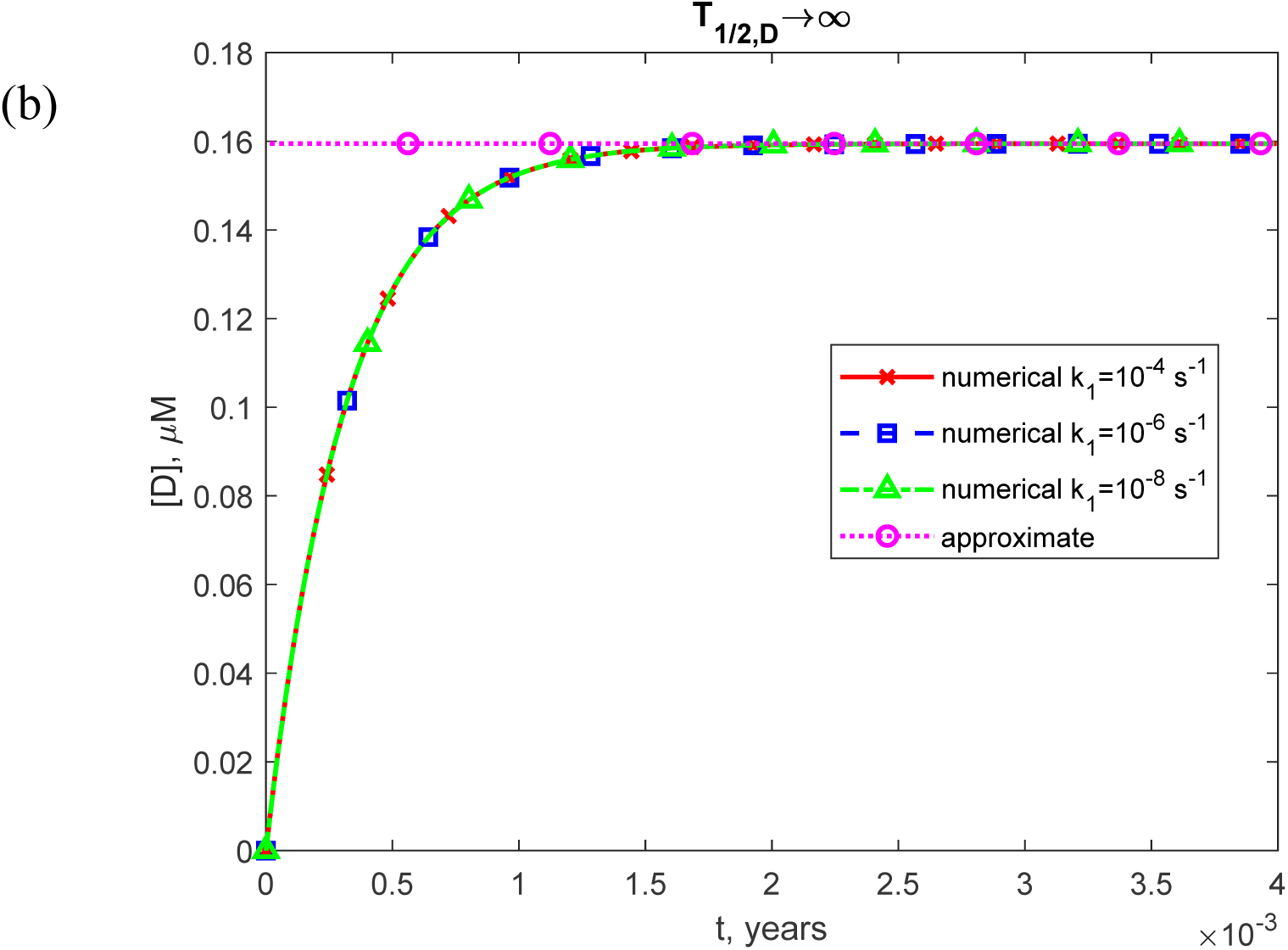
(a) Molar concentration of TDP-43 dimers, [*D*], as a function of time (in years). (b) Similar to Fig. S1a, but with a focus on the specific time range of [0, 0.004 years] on the *x*-axis. The numerical solution is obtained by solving Eqs. (4)-(7) with initial conditions (8). The approximate solution is determined using Eq. (14). The presented scenario assumes dysfunctional protein degradation machinery with *T*→∞. The results are shown for three values of *k*_1/2,*D*_. The case with *k*_1_ = 10^−6^ μM^-1^ s^-1^, *k*_2_ = 10^−4^ s^-1^, and *θ_M_* =10^7^ s.

**Fig. S2.**
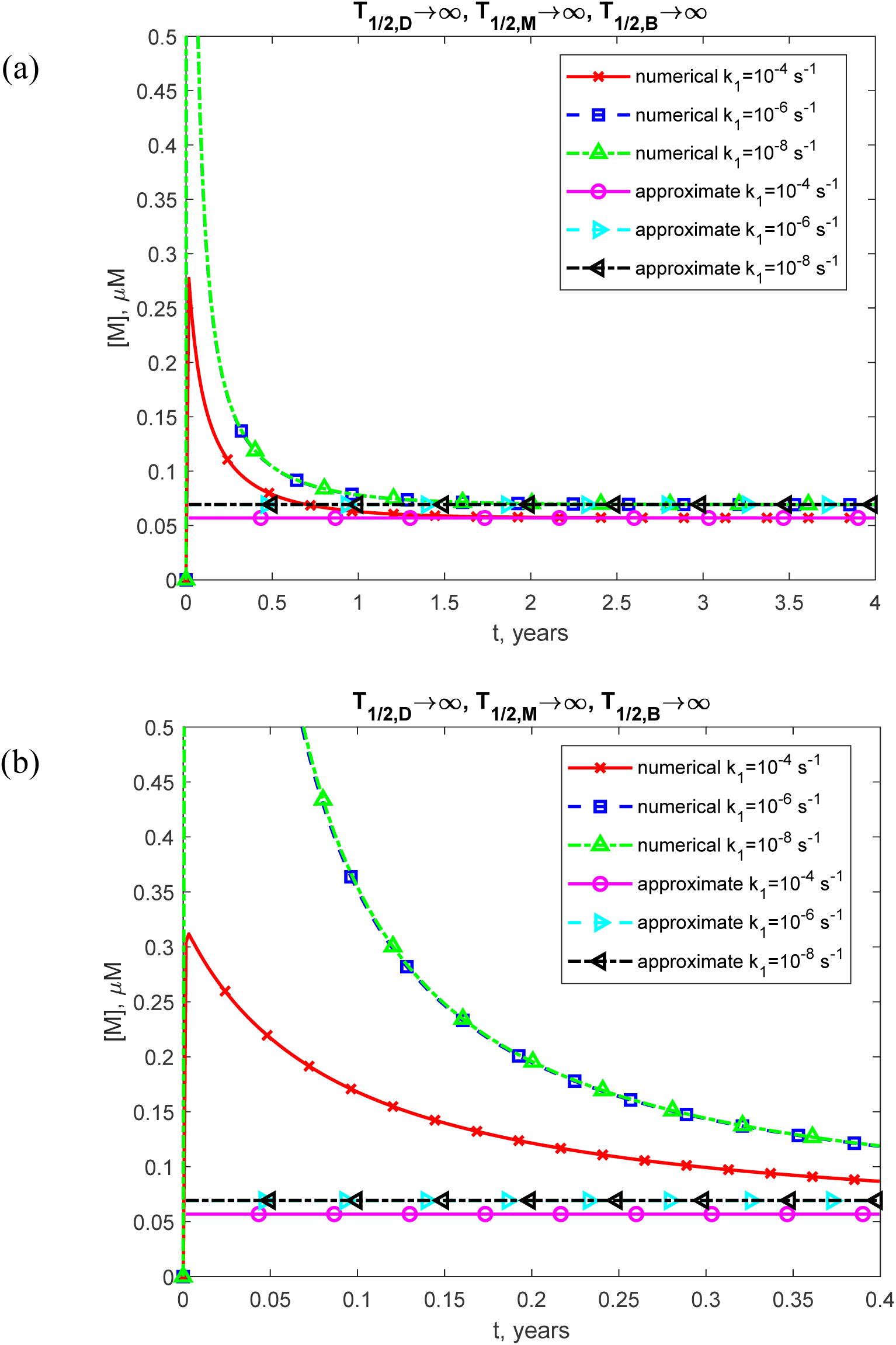
(a) Molar concentration of TDP-43 monomers, [*M*], as a function of time (in years). (b) Similar to Fig. S2a, but with a focus on the specific time range of [0, 0.4 years] on the *x*-axis. The numerical solution is obtained by solving Eqs. (4)-(7) with initial conditions (8). The approximate solution is determined using Eq. (17). The presented scenario assumes dysfunctional protein degradation machinery with *T*_1/ 2,*D*_ →∞, *T*_1/ 2,*M*_ →∞, and *T*_1/ 2,*B*_ →∞. The results are shown for three values of *k*_1_. The case with *k* = 10^−6^ μM^-1^ s^-1^, *k* = 10^−4^ s^-1^, and *θ* =10^7^ s.

**Fig. S3.**
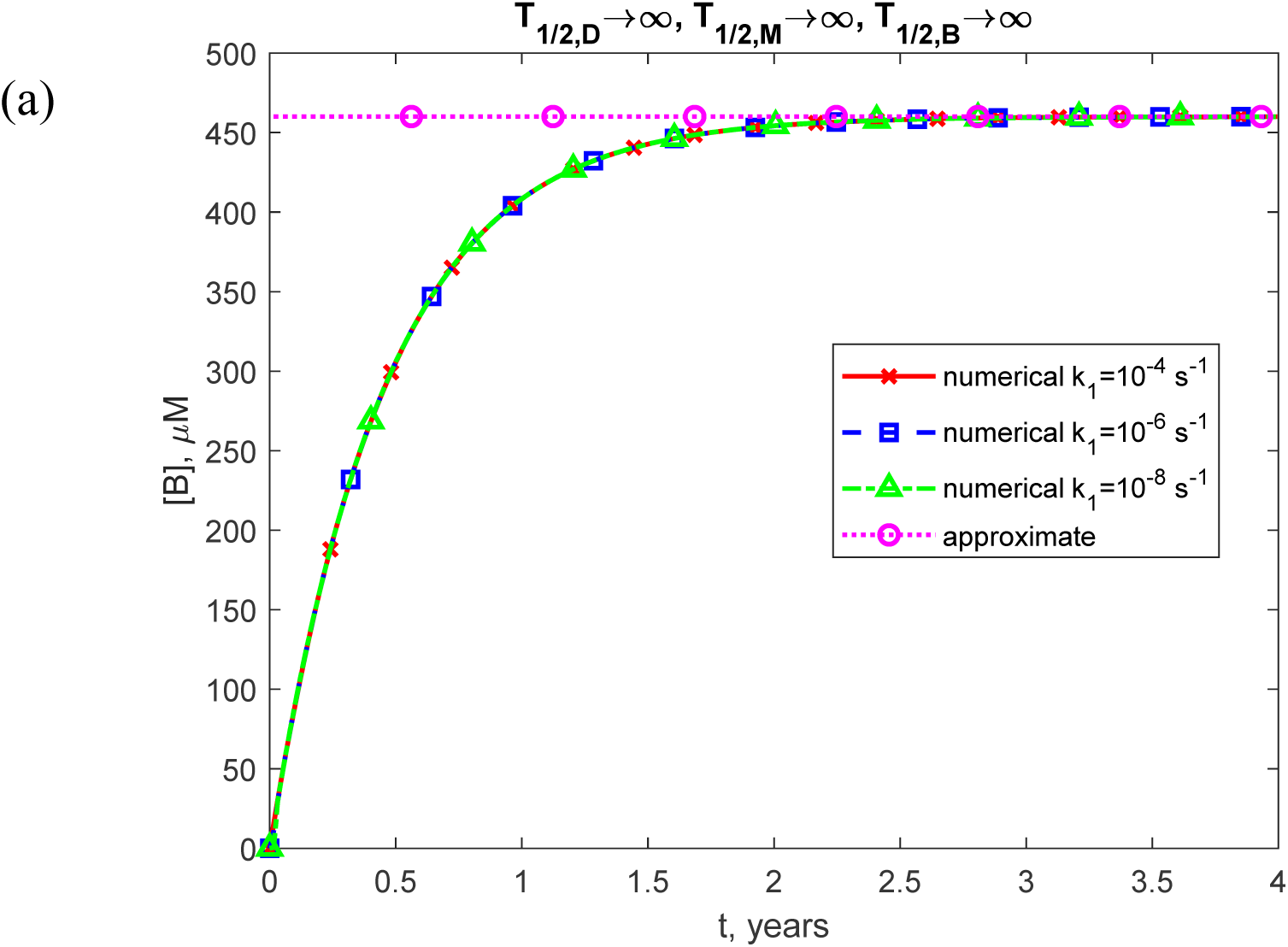
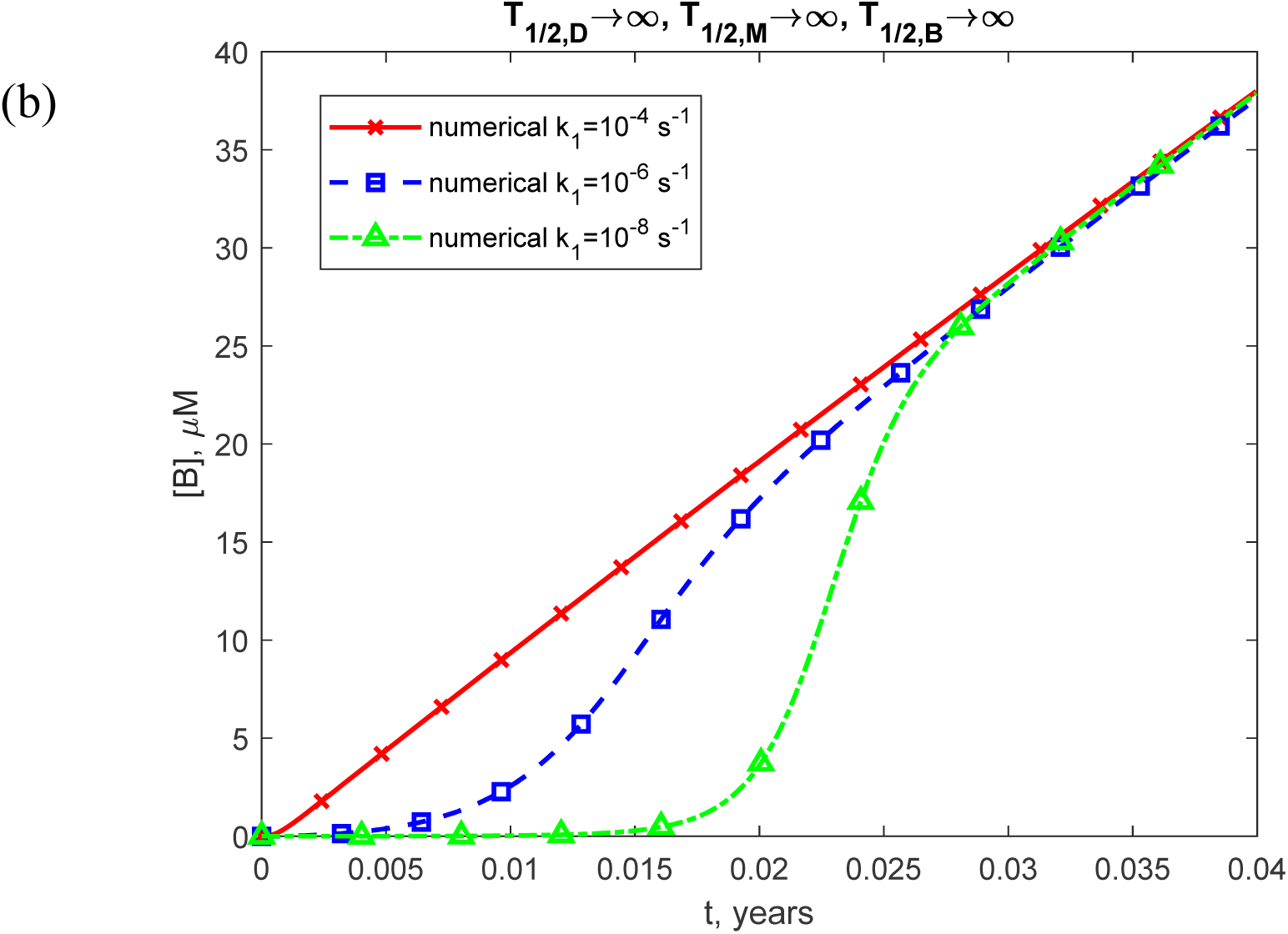
(a) Molar concentration of free misfolded oligomers of TDP-43, [*B*], as a function of time (in years). (b) Similar to Fig. S3a, but with a focus on the specific time range of [0, 0.04 years] on the *x*-axis. The numerical solution is obtained by solving Eqs. (4)-(7) with initial condition (8). The approximate solution is determined using Eq. (16). The presented scenario assumes dysfunctional protein degradation machinery with *T*_1/ 2,*D*_ →∞, *T*_1/ 2,*M*_ →∞, and *T*_1/ 2,*B*_ →∞. The results are shown for three values of *k*_1_. The case with *k*_2_ = 10^−6^ μM^-1^ s^-1^, *k* = 10^−4^ s^-1^, and *θ* =10^7^ s.

**Fig. S4.**
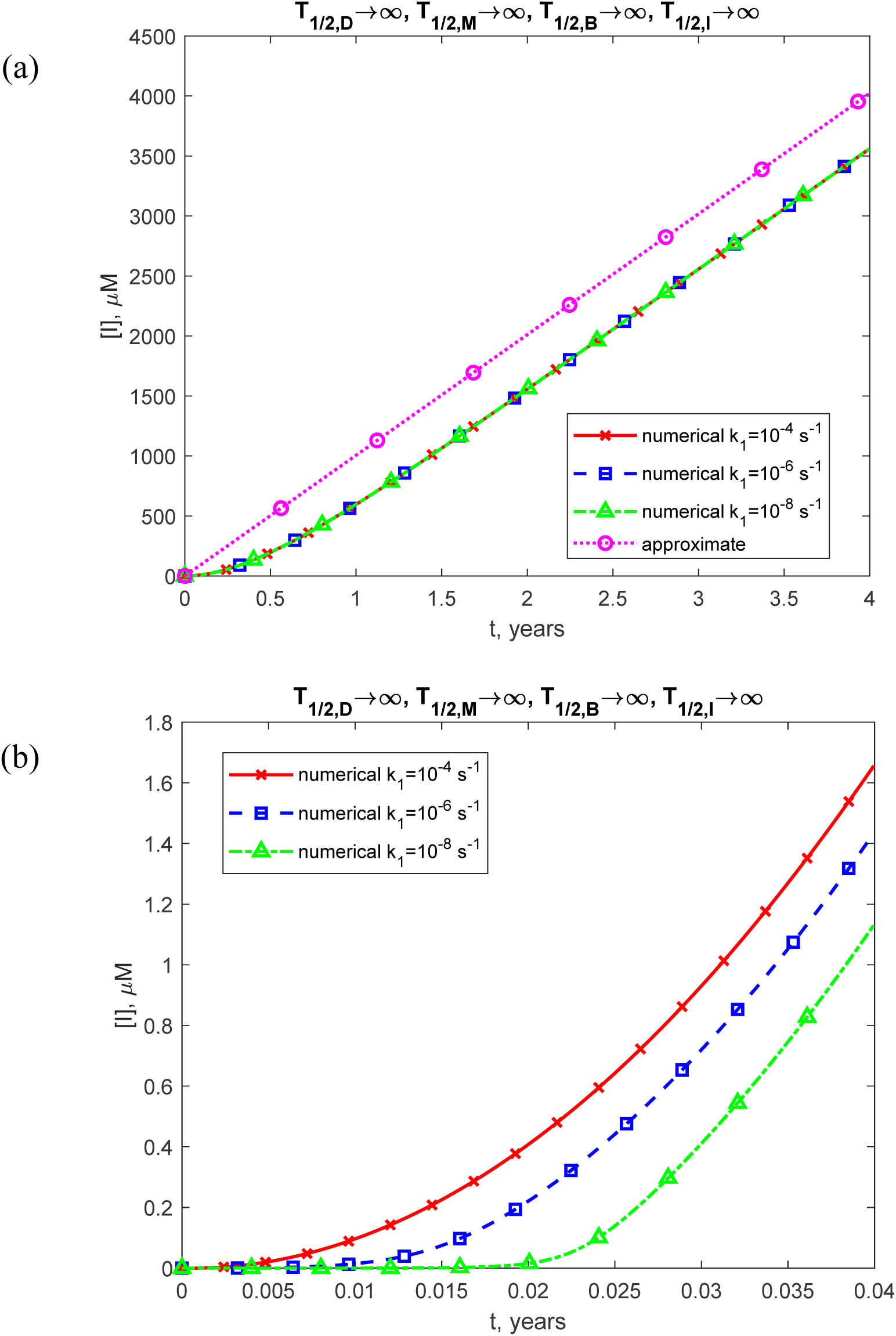
(a) Molar concentration of toxic TDP-43 oligomers incorporated into inclusion bodies, [*I*], as a function of time (in years). (b) Similar to Fig. S4a, but with a focus on the specific time range of [0, 0.04 years] on the *x*-axis. The numerical solution is obtained by solving Eqs. (4)-(7) with initial conditions (8). The approximate solution is determined using Eq. (20). The presented scenario assumes dysfunctional protein degradation machinery with *T*_1/ 2,*D*_ →∞, *T*_1/ 2,*M*_ →∞, *T*_1/ 2,*B*_ →∞, and *T*_1/ 2,*I*_ →∞. The results are shown for three values of *k*_1_. The case with *k*_2_ = 10^−6^ μM^-1^ s^-1^, *k* = 10^−4^ s^-1^, and *θ* =10^7^ s.

**Fig. S5.**
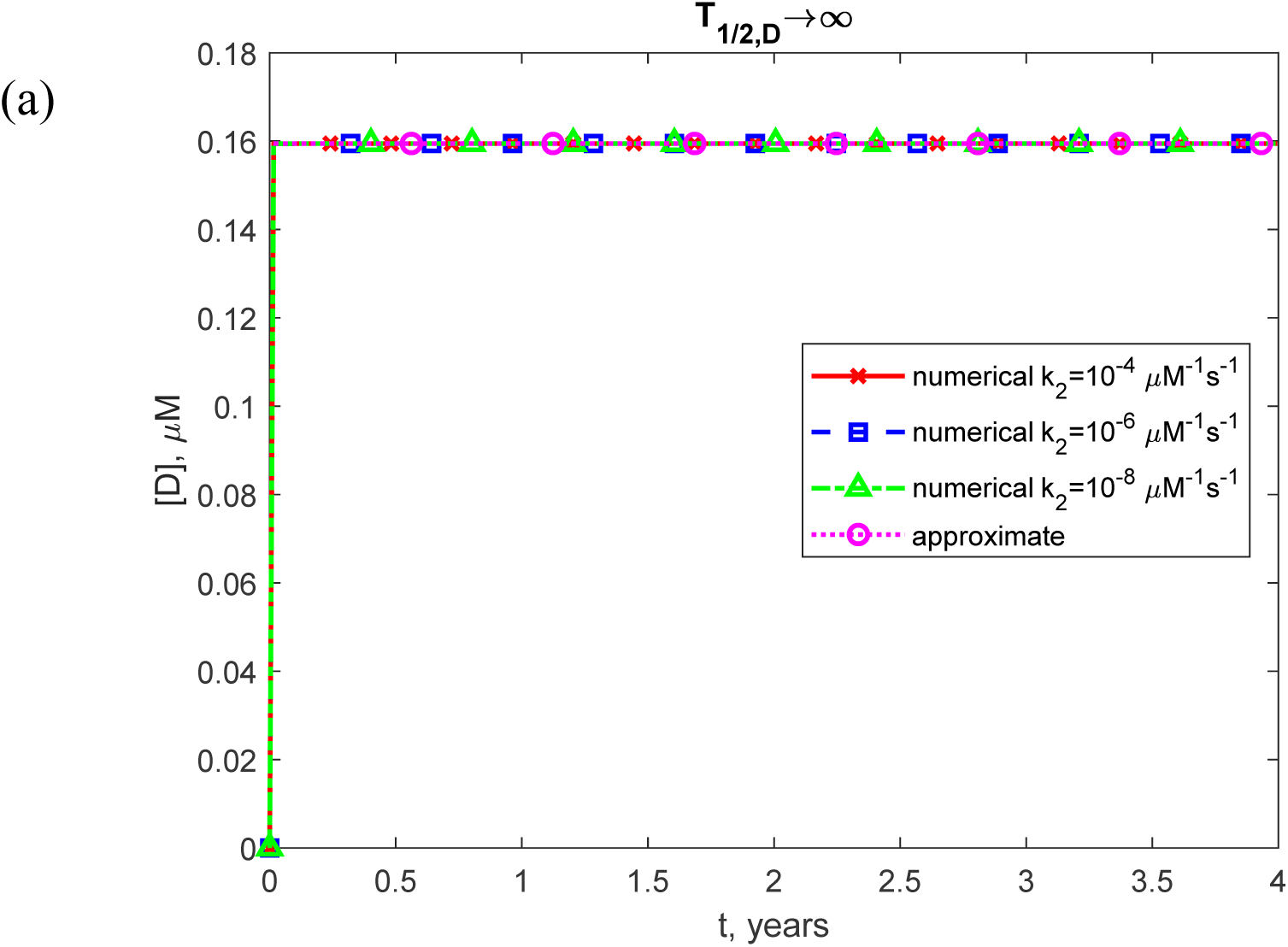
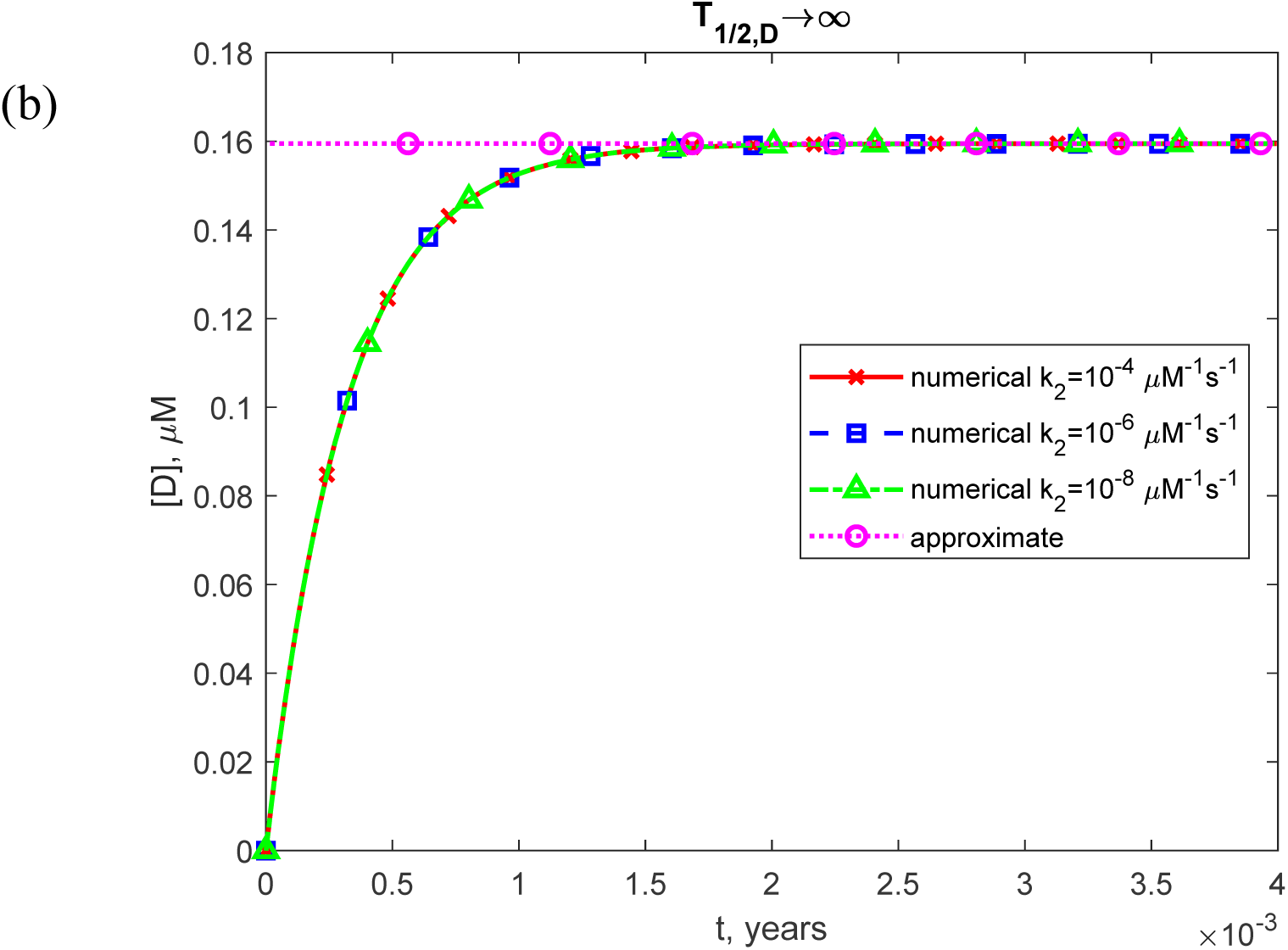
(a) Molar concentration of TDP-43 dimers, [*D*], as a function of time (in years). (b) Similar to Fig. S5a, but with a focus on the specific time range of [0, 0.004 years] on the *x*-axis. The numerical solution is obtained by solving Eqs. (4)-(7) with initial conditions (8). The approximate solution is determined using Eq. (14). The presented scenario assumes dysfunctional protein degradation machinery with *T*_1/2,*D*_ →∞. The results are shown for three values of *k*_2_. The case with *k*_1_ = 10^−6^ s^-1^, *k_M_* = 10^−4^ s^-1^, and *θ* =10^7^ s.

**Fig. S6.**
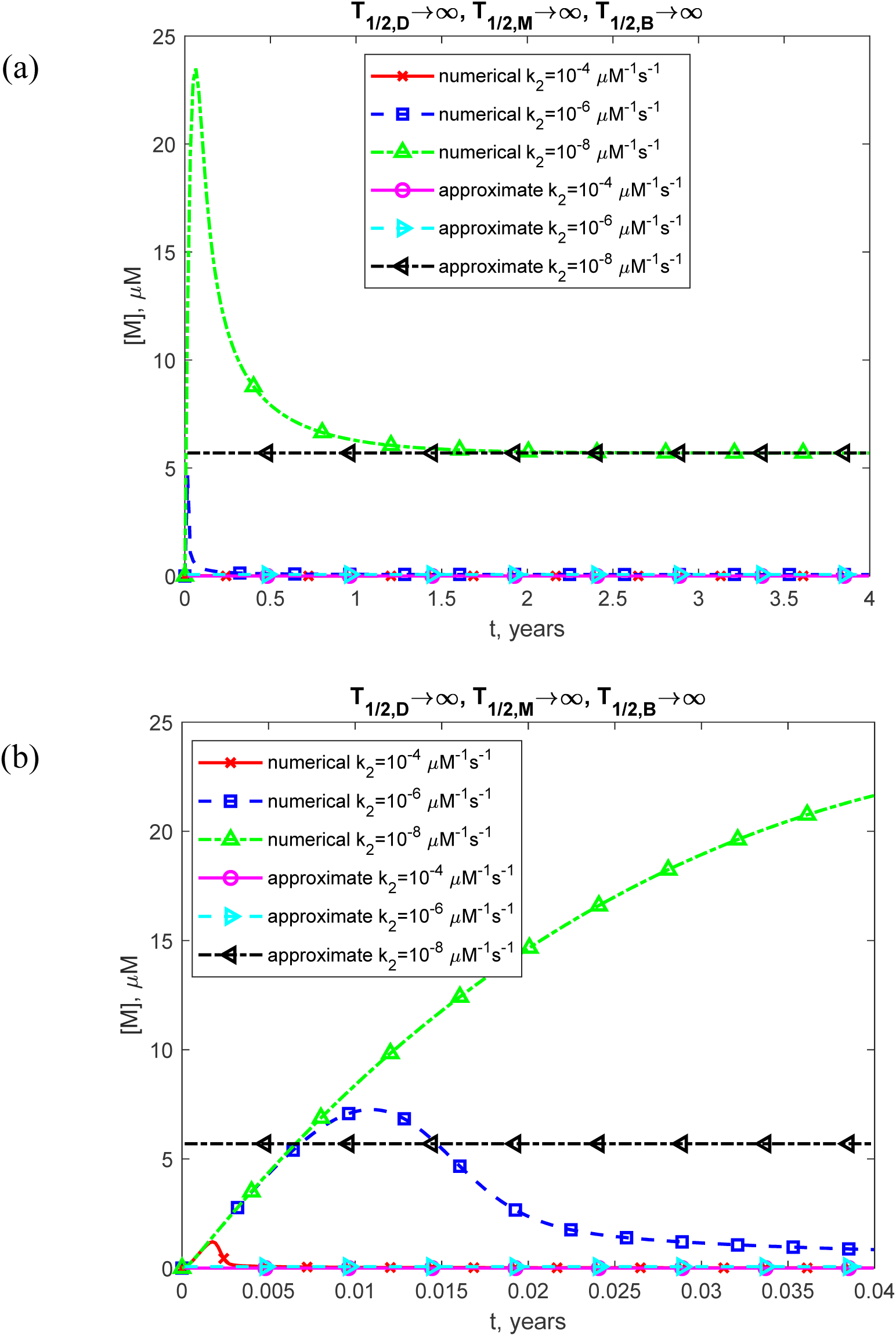
(a) Molar concentration of TDP-43 monomers, [*M*], as a function of time (in years). (b) Similar to Fig. S6a, but with a focus on the specific time range of [0, 0.04 years] on the *x*-axis. The numerical solution is obtained by solving Eqs. (4)-(7) with initial conditions (8). The approximate solution is determined using Eq. (17). The presented scenario assumes dysfunctional protein degradation machinery with *T*_1/ 2,*D*_ →∞, *T*_1/ 2,*M*_ →∞, and *T*_1/ 2,*B*_ →∞. The results are shown for three values of *k*_2_. The case with *k* = 10^−6^ s^-1^, *k* = 10^−4^ s^-1^, and *θ* =10^7^ s.

**Fig. S7.**
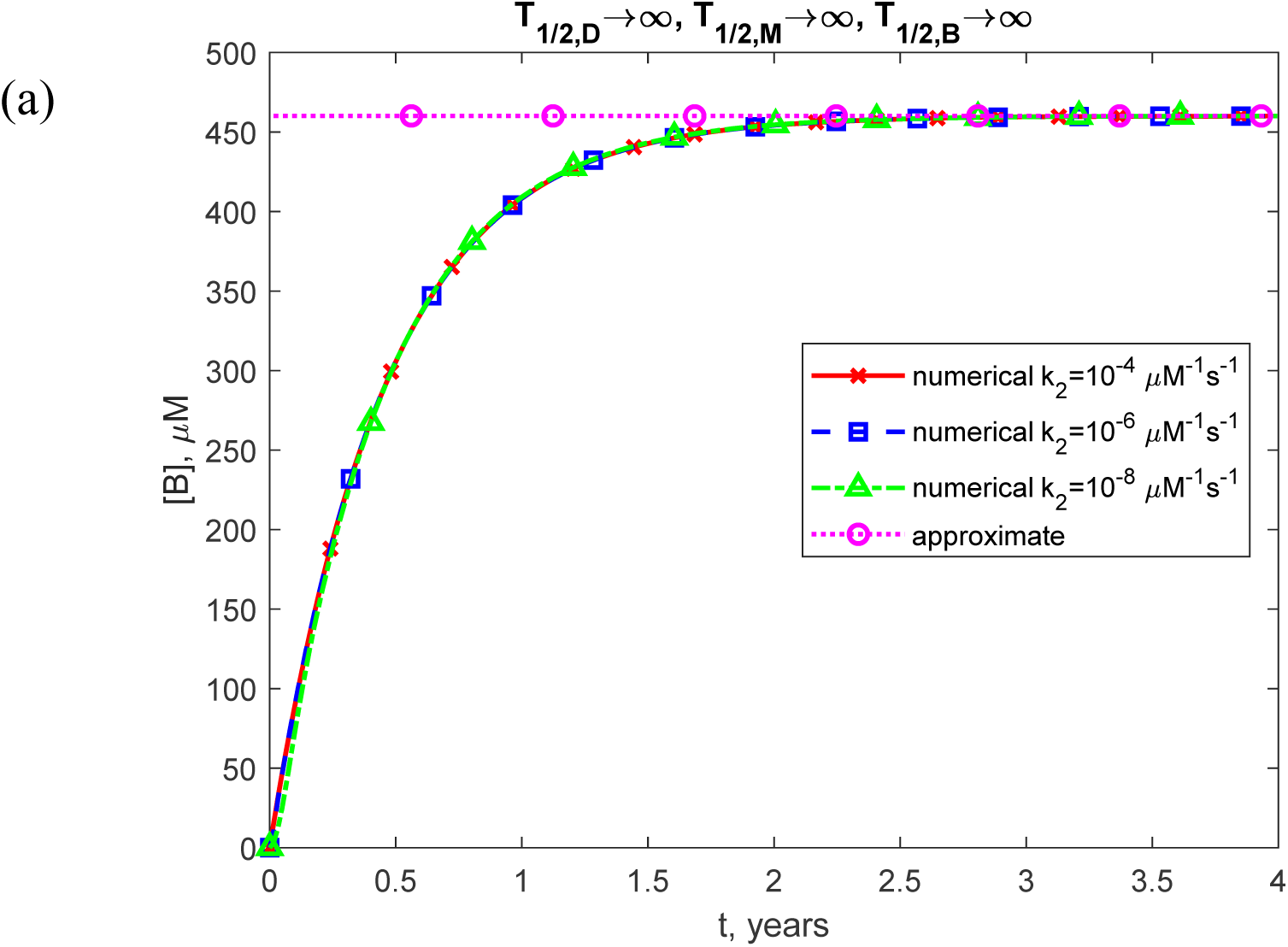
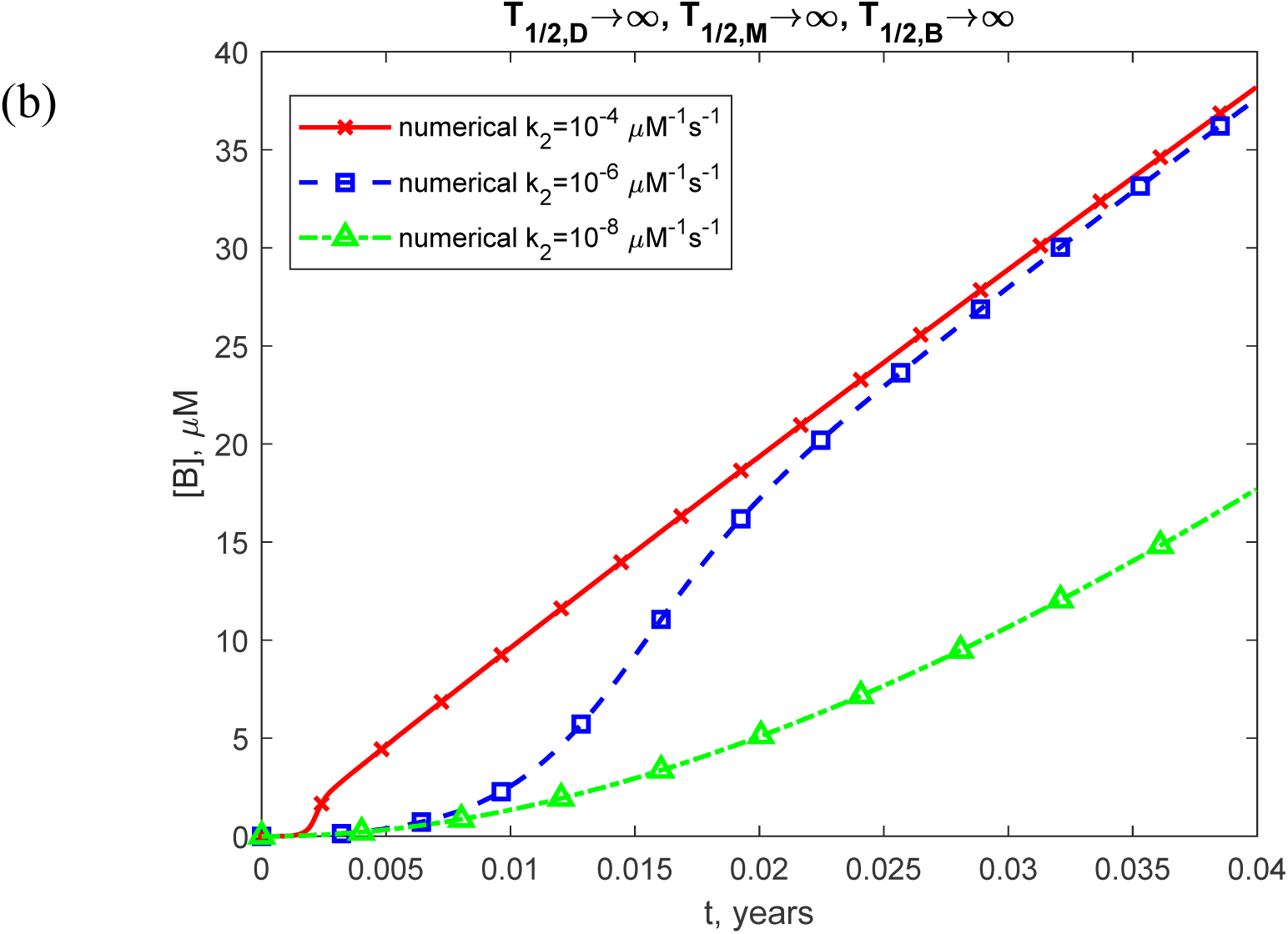
(a) Molar concentration of free misfolded TDP-43 oligomers, [*B*], as a function of time (in years). (b) Similar to Fig. S7a, but with a focus on the specific time range of [0, 0.04 years] on the *x*-axis. The numerical solution is obtained by solving Eqs. (4)-(7) with initial conditions (8), while the approximate solution is determined using Eq. (16). The presented scenario assumes dysfunctional protein degradation machinery with *T*_1/ 2,*D*_ →∞, *T*_1/ 2,*M*_ →∞, and *T*_1/ 2,*B*_ →∞. The results are shown for three values of *k*_2_. The case with *k*_1_ = 10^−6^ s^-1^, *k* = 10^−4^ s^-1^, and *θ* =10^7^ s.

**Fig. S8.**
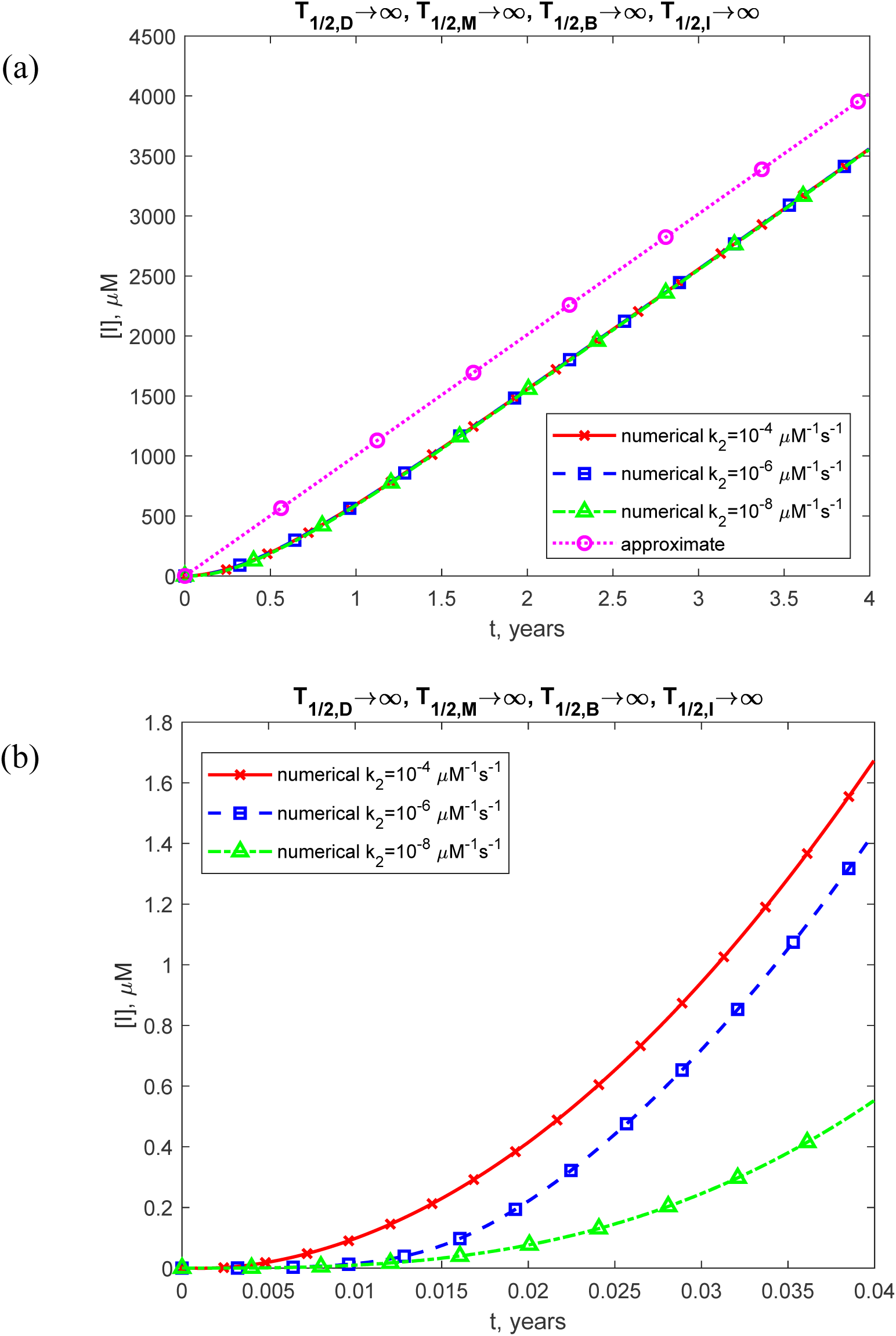
(a) Molar concentration of toxic TDP-43 oligomers incorporated into inclusion bodies, [*I*], as a function of time (in years). (b) Similar to Fig. S8a, but with a focus on the specific time range of [0, 0.04 years] on the *x*-axis. The numerical solution is obtained by solving Eqs. (4)-(7) with initial conditions (8), while the approximate solution is determined using Eq. (20). The presented scenario assumes dysfunctional protein degradation machinery with *T*_1/ 2,*D*_ →∞, *T*_1/ 2,*M*_ →∞, *T*_1/ 2,*B*_ →∞, and *T*_1/ 2,*I*_ →∞. The results are shown for three values of *k*_2_. The case with *k*_1_ = 10^−6^ s^-1^, *k_M_* = 10^−4^ s^-1^, and *θ*_1/2,*B*_ =10^7^ s.

**Fig. S9.**
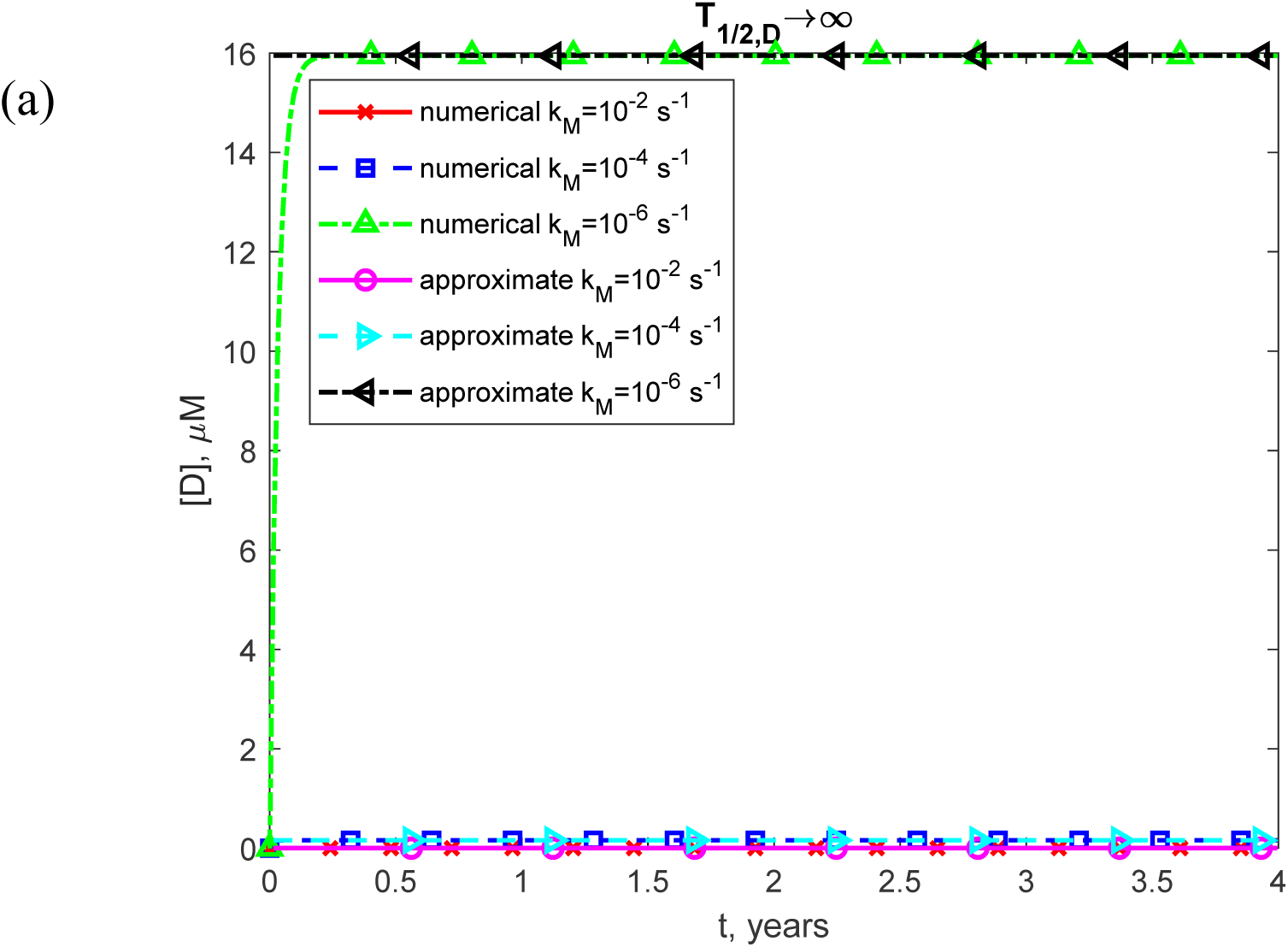
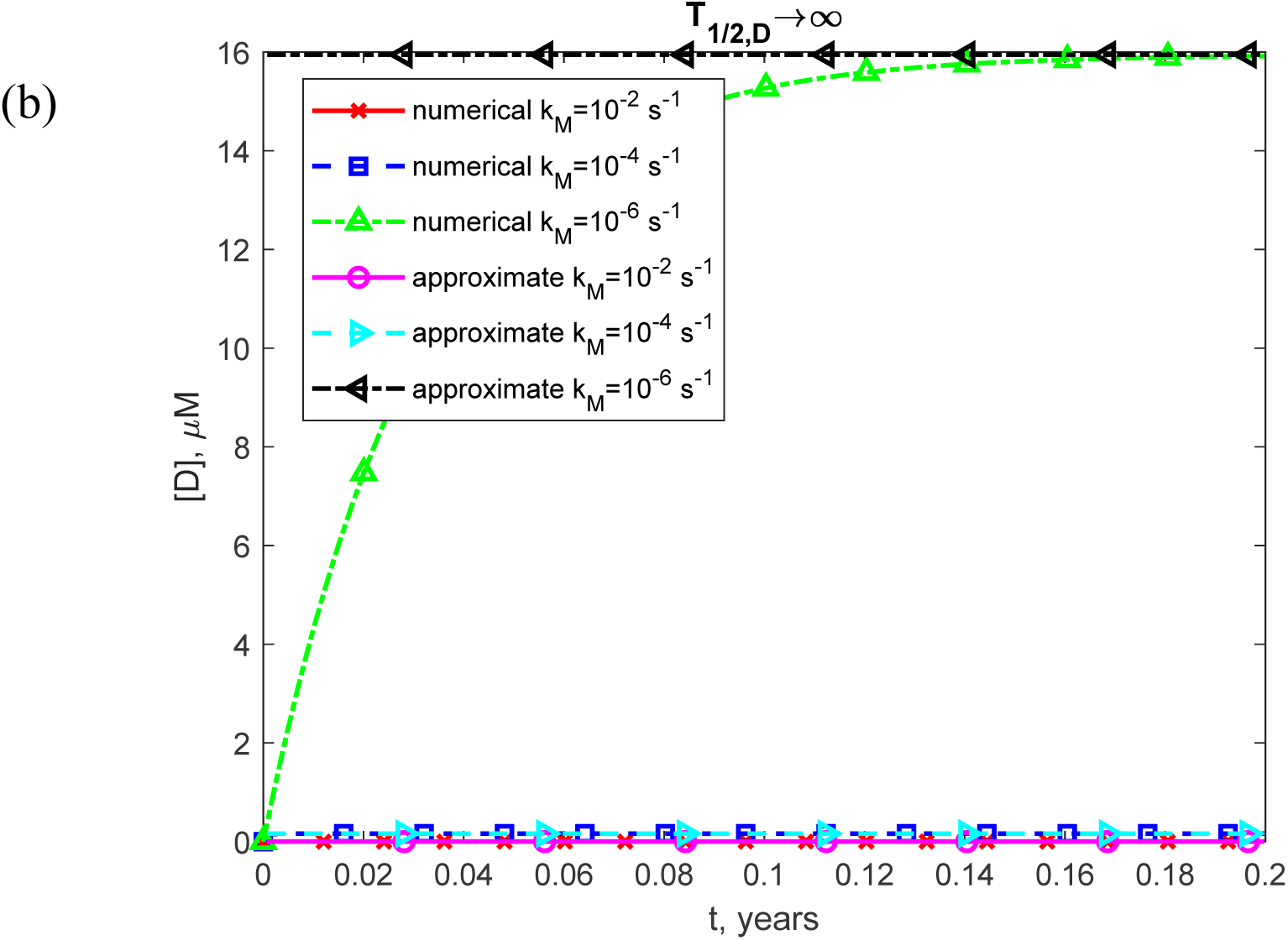
(a) Molar concentration of TDP-43 dimers, [*D*], as a function of time (in years). (b) Similar to Fig. S9a, but with a focus on the specific time range of [0, 0.2 years] on the *x*-axis. The numerical solution is obtained by solving Eqs. (4)-(7) with initial conditions (8), while the approximate solution is determined using Eq. (14). The presented scenario assumes dysfunctional protein degradation machinery with *T*→∞. The results are shown for three values of *k*_1/2,*D*_. The case with *k_M_* = 10^−6^ s^-1^, *k_1_* = 10^−6^ μM^-1^ s^-1^, and *θ_2_* =10^7^ s.

**Fig. S10.**
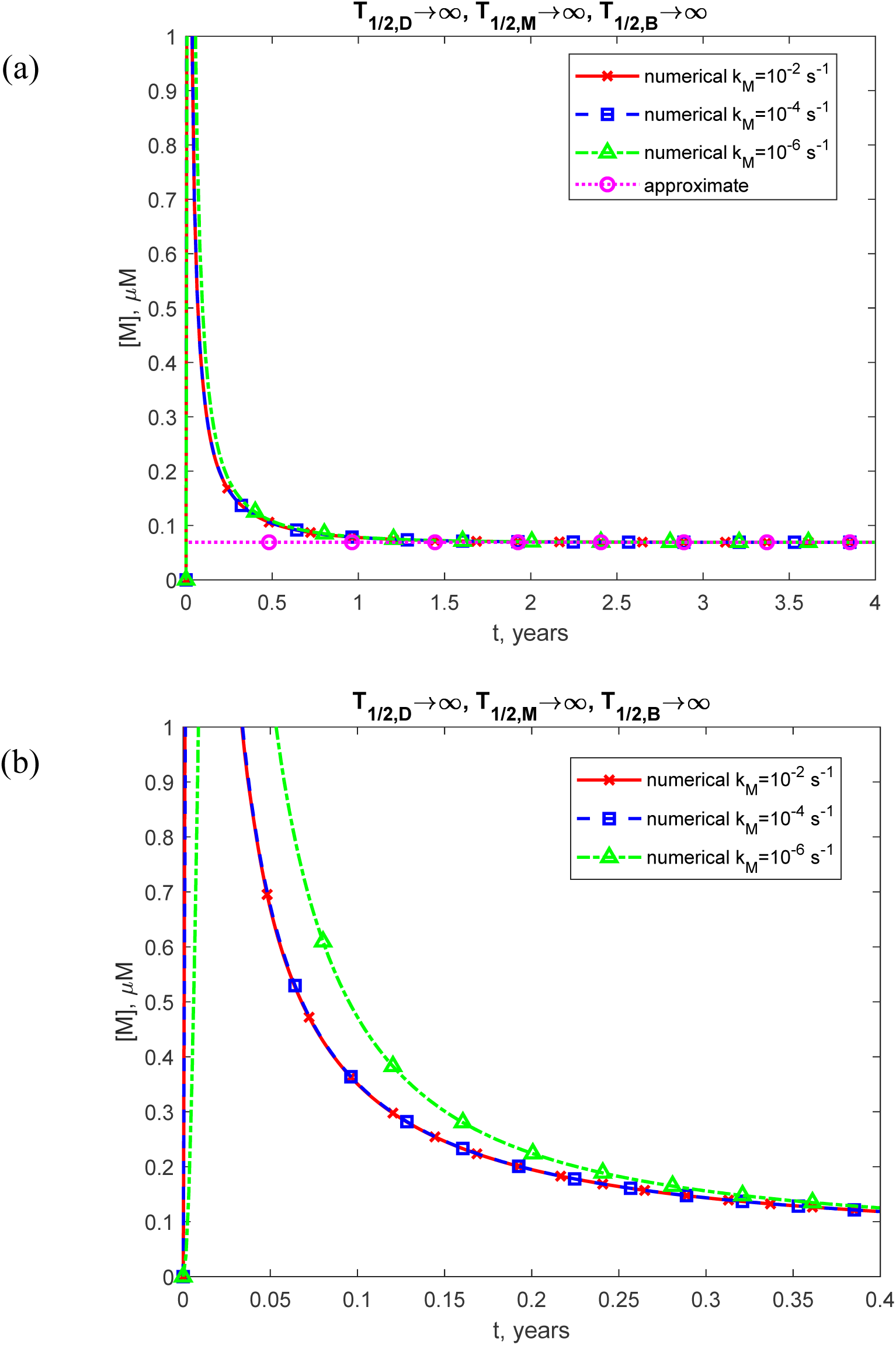
(a) Molar concentration of TDP-43 monomers, [*M*], as a function of time (in years). (b) Similar to Fig. S10a, but with a focus on the specific time range of [0, 0.4 years] on the *x*-axis. The numerical solution is obtained by solving Eqs. (4)-(7) with initial conditions (8), while the approximate solution is determined using Eq. (17). The presented scenario assumes dysfunctional protein degradation machinery with *T*_1/ 2,*D*_ →∞, *T*_1/ 2,*M*_ →∞, and *T*_1/ 2,*B*_ →∞. The results are shown for three values of *k_M_*. The case with *k*_1_ = 10^−6^ s^-1^, *k* = 10^−6^ μM^-1^ s^-1^, and *θ* =10^7^ s.

**Fig. S11.**
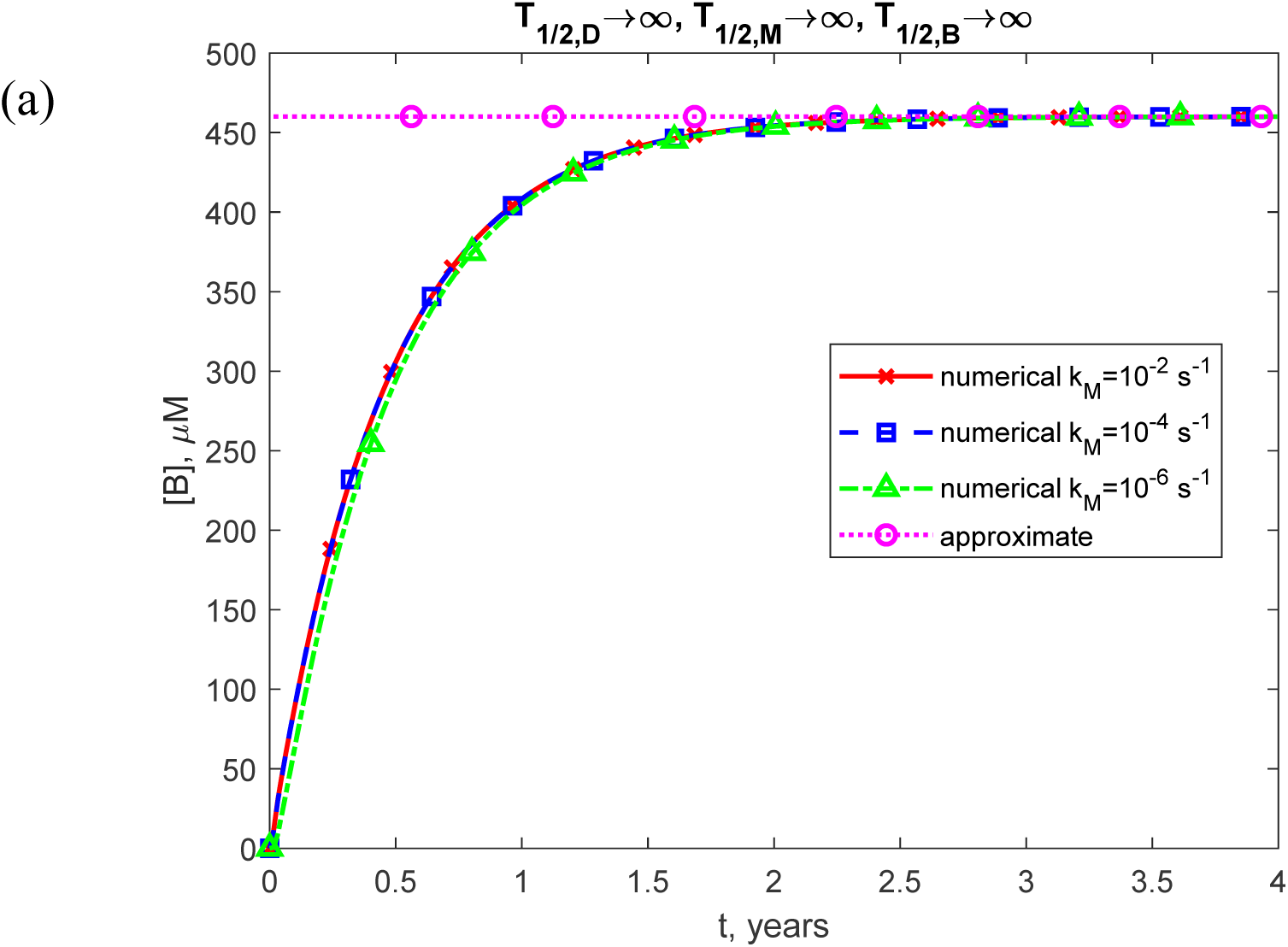
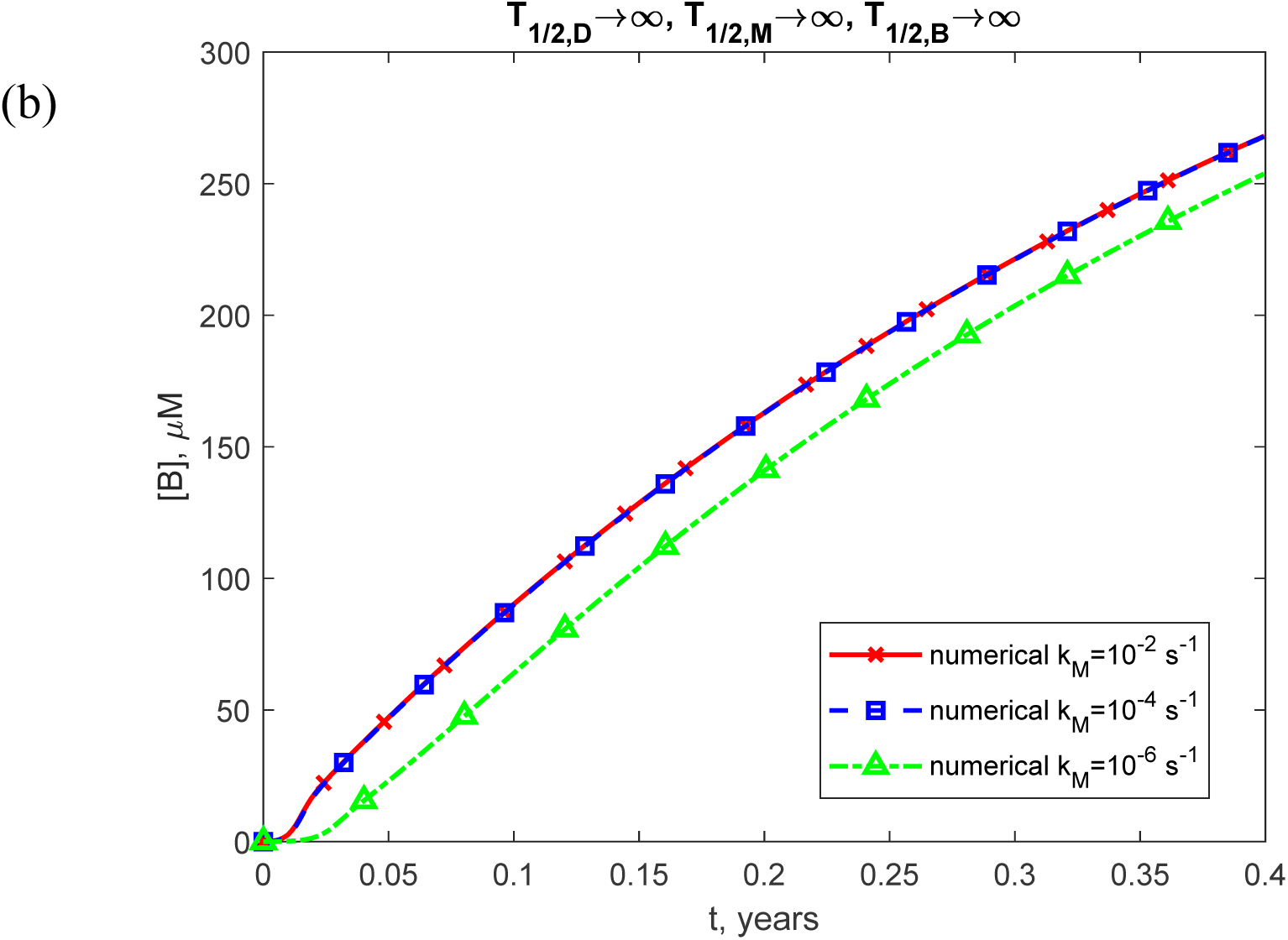
(a) Molar concentration of free misfolded oligomers of TDP-43, [*B*], as a function of time (in years). (b) Similar to Fig. S11a, but with a focus on the specific time range of [0, 0.4 years] on the *x*-axis. The numerical solution is obtained by solving Eqs. (4)-(7) with initial conditions (8), while the approximate solution is determined using Eq. (16). The presented scenario assumes dysfunctional protein degradation machinery with *T*_1/ 2,*D*_ →∞, *T*_1/ 2,*M*_ →∞, and *T*_1/ 2,*B*_ →∞. The results are shown for three values of *k_M_*. The case with *k_1_* = 10^−6^ s^-1^, *k_2_* = 10^−6^ μM^-1^ s^-1^, and *θ*_1/2,*B*_ =10^7^ s.

**Fig. S12.**
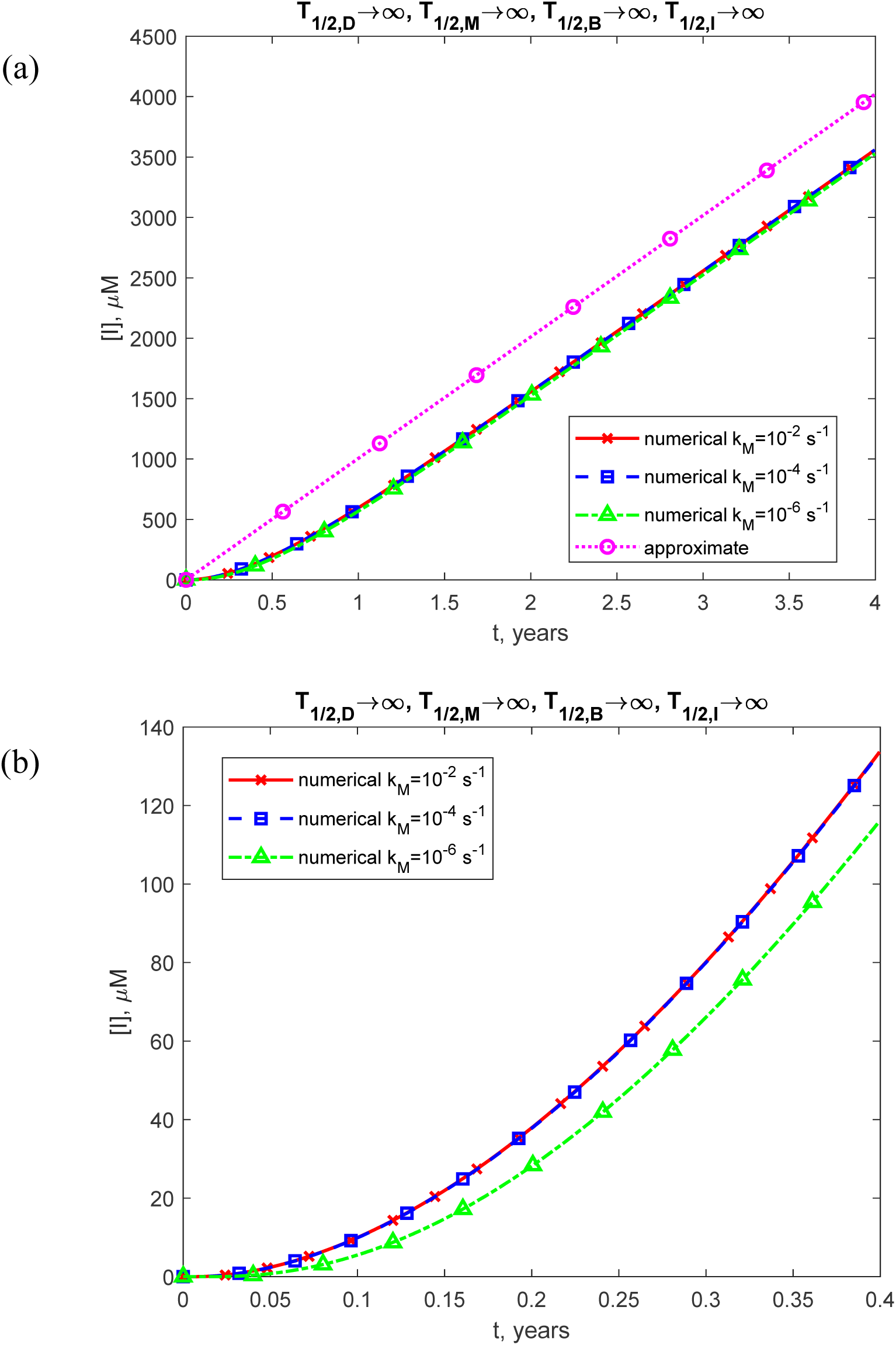
(a) Molar concentration of toxic TDP-43 oligomers incorporated into inclusion bodies, [*I*], as a function of time (in years). (b) Similar to Fig. S12a, but with a focus on the specific time range of [0, 0.4 years] on the *x*-axis. The numerical solution is obtained by solving Eqs. (4)-(7) with initial conditions (8), while the approximate solution is determined using Eq. (20). The presented scenario assumes dysfunctional protein degradation machinery with *T*_1/ 2,*D*_ →∞, *T*_1/ 2,*M*_ →∞, *T*_1/ 2,*B*_ →∞, and *T*_1/ 2,*I*_ →∞. The results are shown for three values of *k*. The case with *k_M_* = 10^−6^ s^-1^, *k_1_* = 10^−6^ μM^-1^ s^-1^, and *θ_2_* =10^7^ s.

**Fig. S13.**
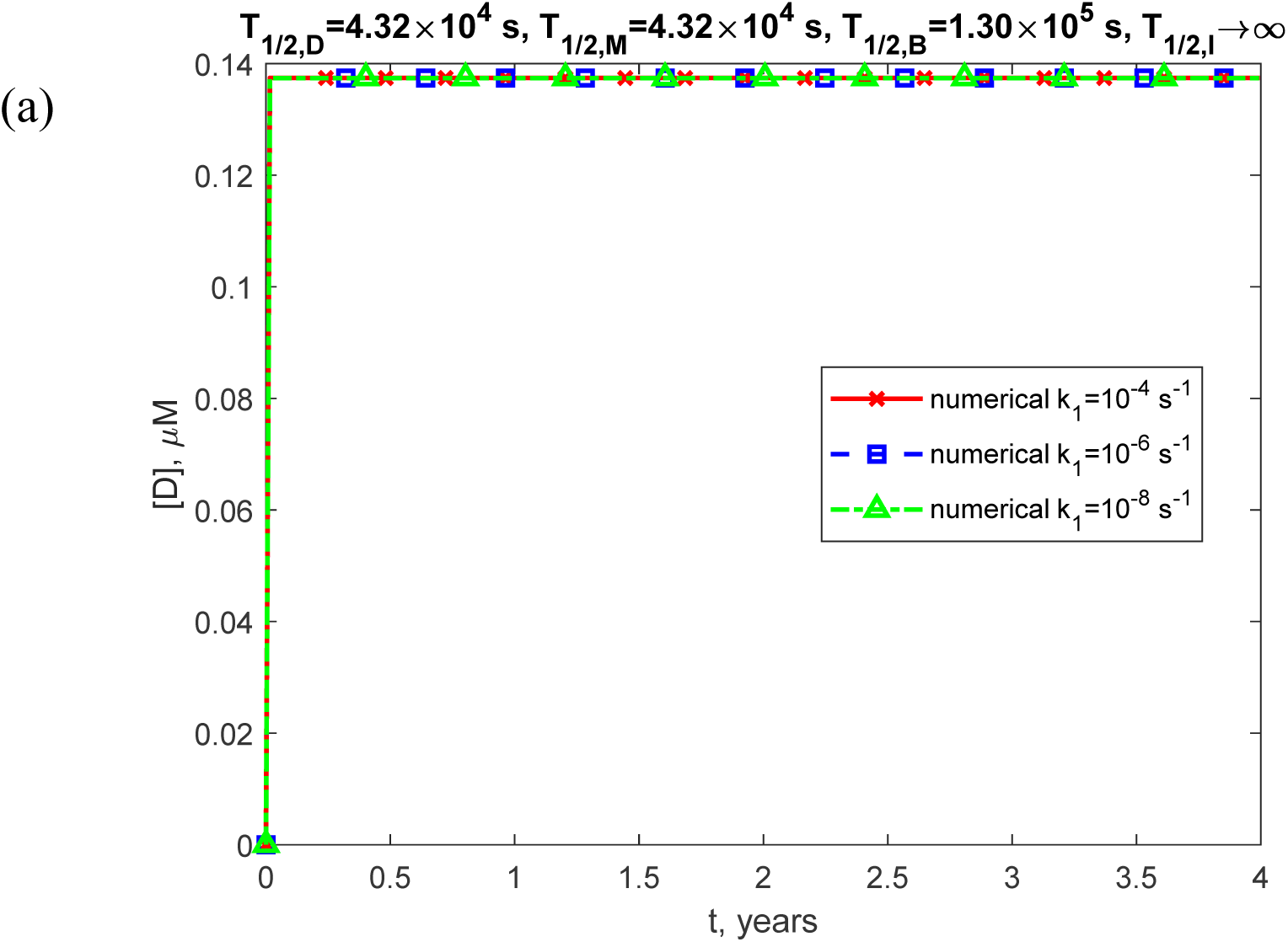
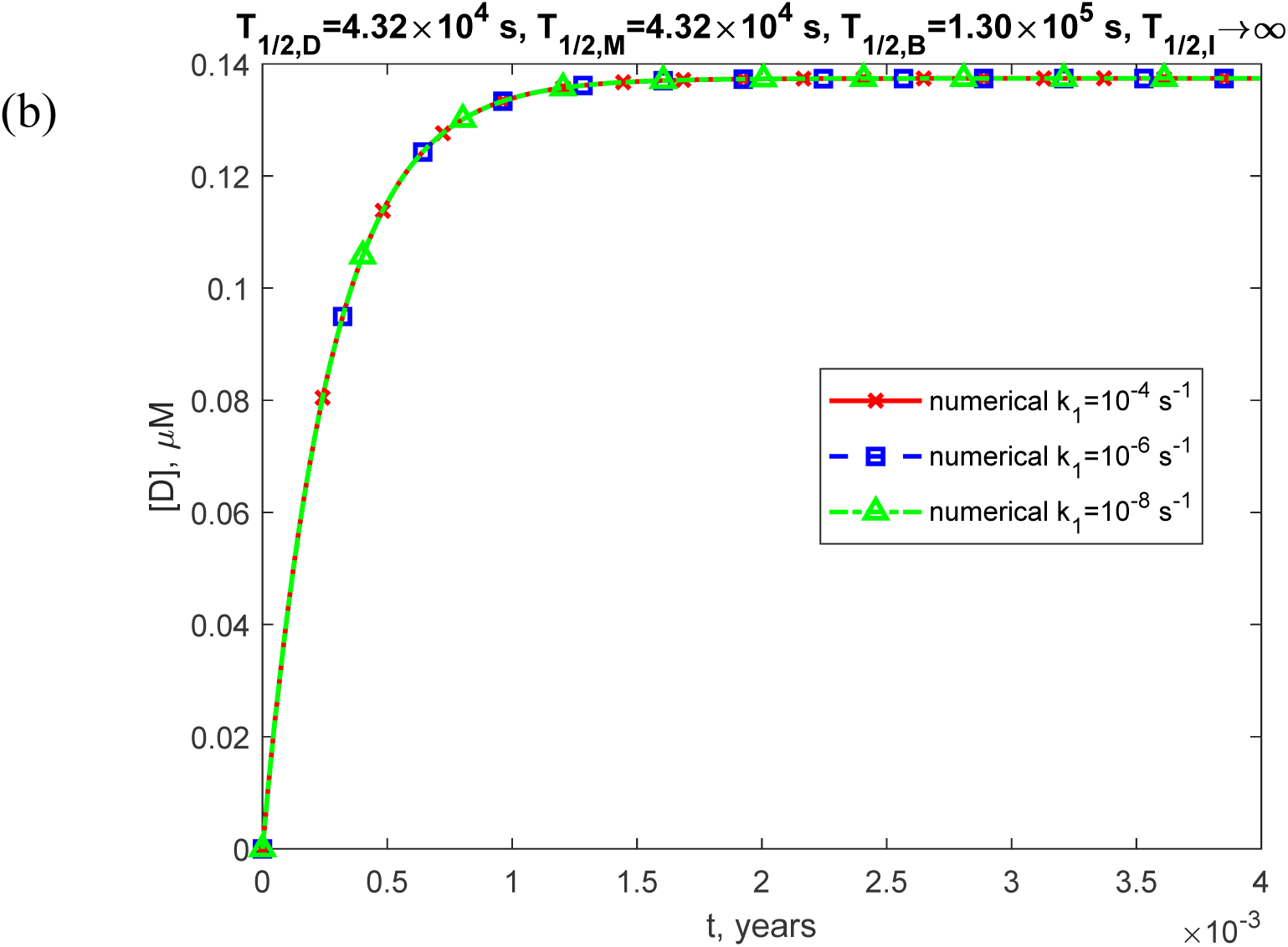
(a) Molar concentration of TDP-43 dimers, [*D*], as a function of time (in years). (b) Similar to Fig. S13a, but with a focus on the specific time range of [0, 0.004 years] on the *x*-axis. The numerical solution is obtained by solving Eqs. (4)-(7) with initial conditions (8). The presented scenario assumes a physiologically relevant half-life for TDP-43 dimers: *T* = 4.32×10^4^ s. The approximate solution is not shown, as it applies only to the scenario with infinite half-lives of TDP-43 dimers, monomers, and free misfolded oligomers. The results are displayed for three values of *k*_1_. The case with *k*_2_ = 10^−6^ μM^-1^ s^-1^, *k* = 10^−4^ s^-1^, and *θ* =10^7^ s.

**Fig. S14.**
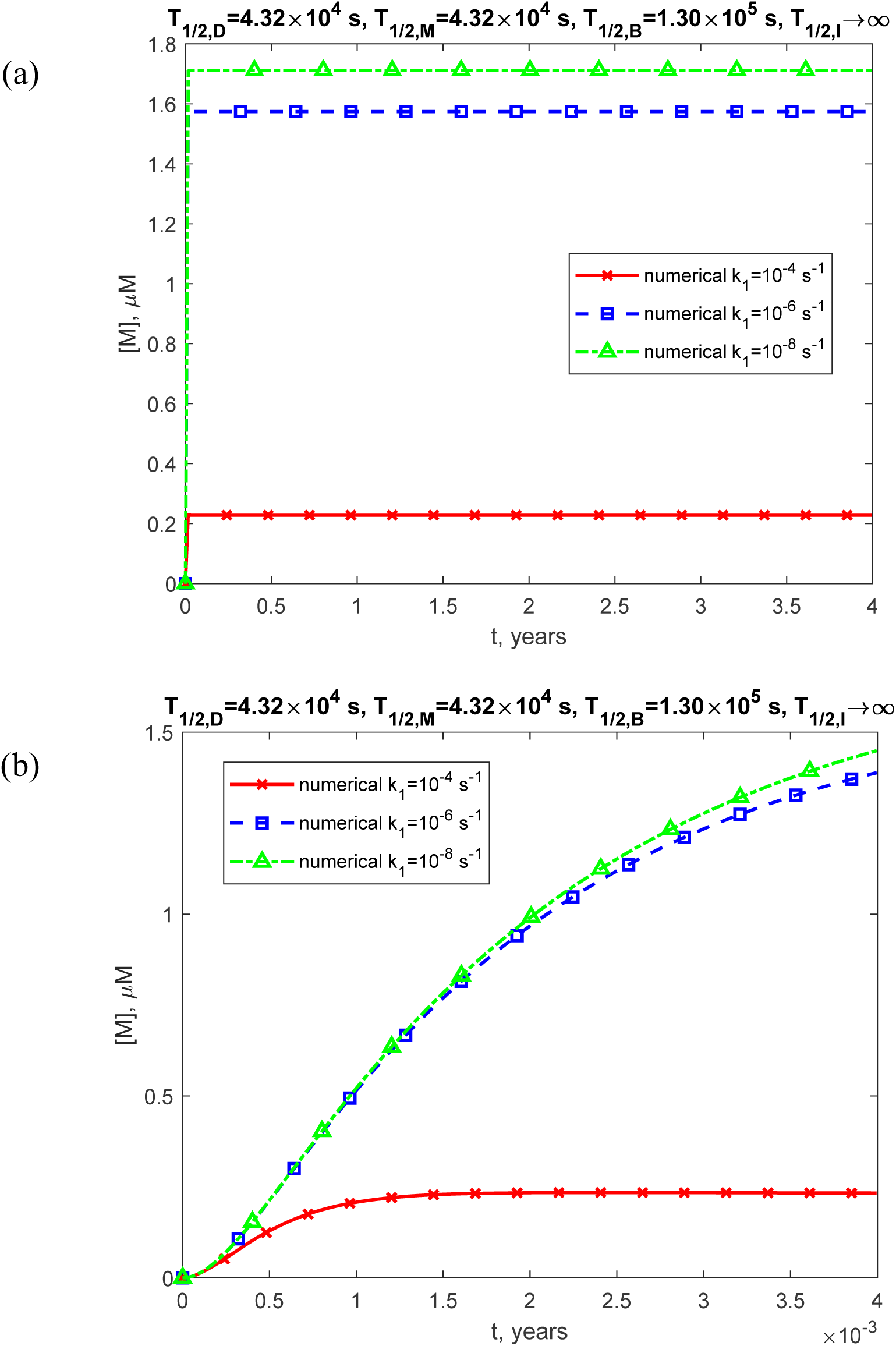
(a) Molar concentration of TDP-43 monomers, [*M*], as a function of time (in years). (b) Similar to Fig. S14a, but with a focus on the specific time range of [0, 0.004 years] on the *x*-axis. The numerical solution is obtained by solving Eqs. (4)-(7) with initial conditions (8). The presented scenario assumes physiologically relevant half-lives for TDP-43 dimers, monomers, and free misfolded oligomers: *T*_1/2,*D*_ = 4.32×10^4^ s, *T*_1/2,*M*_ = 4.32×10^4^ s, and *T*_1/2,*B*_ =1.30×10^5^ s. The approximate solution is not shown, as it applies only to the scenario with infinite half-lives of TDP-43 dimers, monomers, and free misfolded oligomers. The results are displayed for three values of *k*_1_. The case with *k*_2_ = 10^−6^ μM^-1^ s^-1^, *k* = 10^−4^ s^-1^, and *θ* =10^7^ s.

**Fig. S15.**
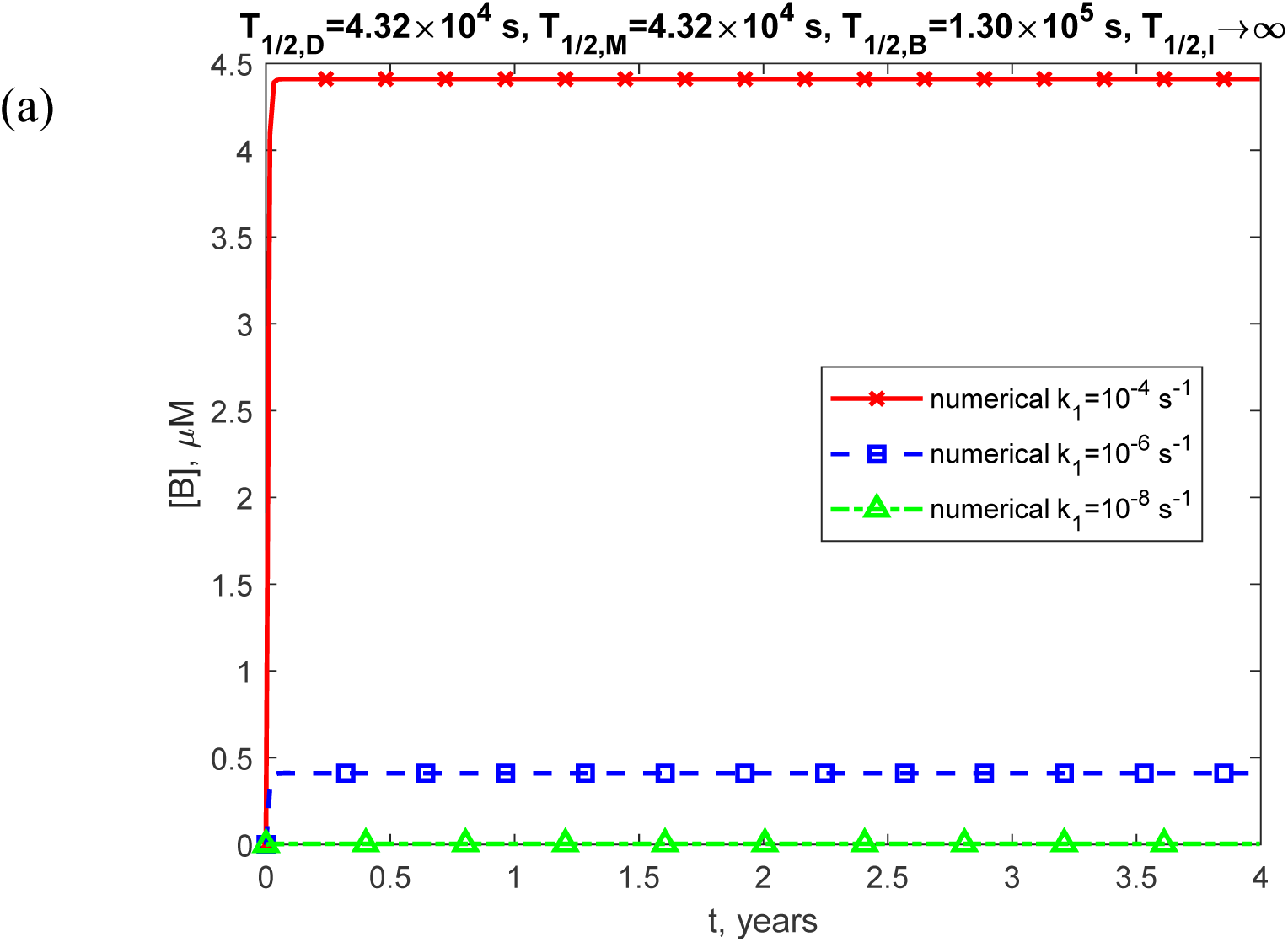
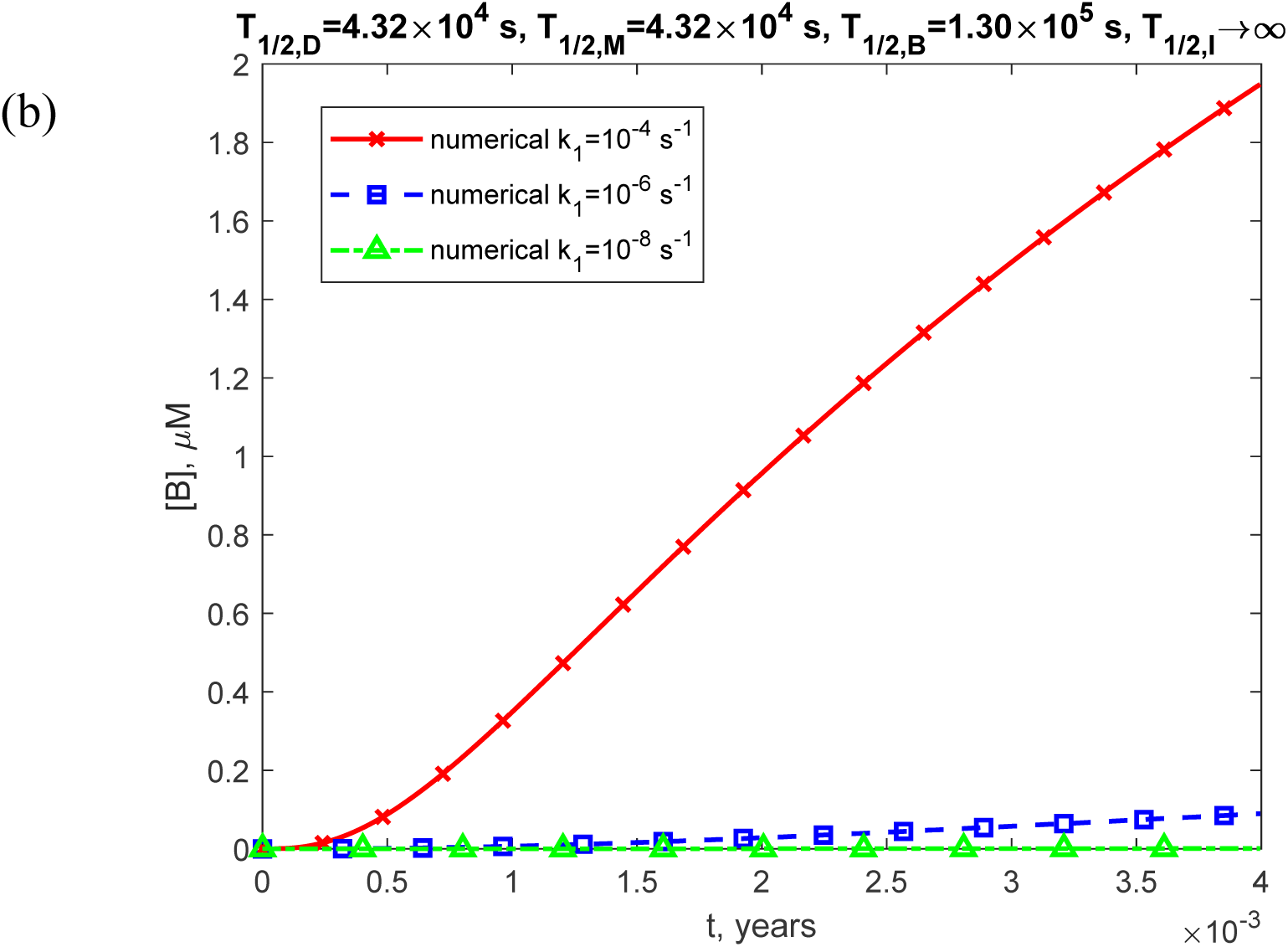
(a) Molar concentration of free misfolded oligomers of TDP-43, [*B*], as a function of time (in years). (b) Similar to Fig. S15a, but with a focus on the specific time range of [0, 0.004 years] on the *x*-axis. The numerical solution is obtained by solving Eqs. (4)-(7) with initial conditions (8). The presented scenario assumes physiologically relevant half-lives for TDP-43 dimers, monomers, and free misfolded oligomers: *T*_1/2,*D*_ = 4.32×10^4^ s, *T*_1/2,*M*_ = 4.32×10^4^ s, and *T*_1/2,*B*_ =1.30×10^5^ s. The approximate solution is not shown, as it applies only to the scenario with infinite half-lives of TDP-43 dimers, monomers, and free misfolded oligomers. The results are displayed for three values of *k*_1_. The case with *k*_2_ = 10^−6^ μM s^1^, *k_M_* = 10^−4^ s^−1^, and *θ* =10^7^ s.

**Fig. S16.**
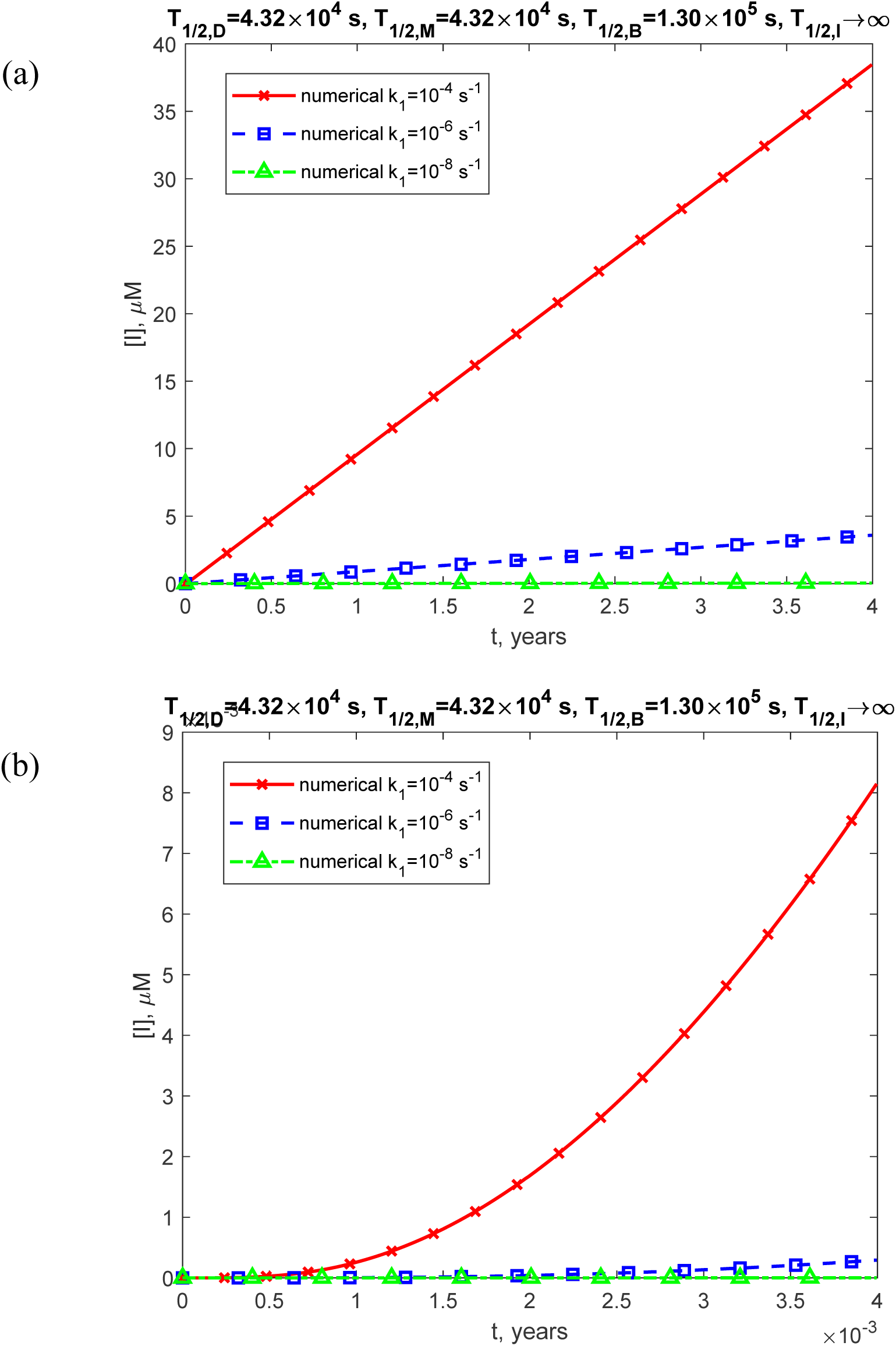
(a) Molar concentration of toxic TDP-43 oligomers incorporated into inclusion bodies, [*I*], as a function of time (in years). (b) Similar to Fig. S16a, but with a focus on the specific time range of [0, 0.004 years] on the *x*-axis. The numerical solution is obtained by solving Eqs. (4)-(7) with initial conditions (8). The presented scenario assumes physiologically relevant half-lives for TDP-43 dimers, monomers, free misfolded oligomers, and oligomers deposited into inclusion bodies: *T*_1/2,*M*_ = 4.32×10^4^ s, *T*_1/2,*B*_ = 4.32×10^4^ s, *T* =1.30×10^5^ s, and *T*_1/2,*I*_ →∞. The approximate solution is not shown, as it applies only to the scenario with infinite half-lives of TDP-43 dimers, monomers, and free misfolded oligomers. The results are displayed for three values of *k*_1_. The case with *k*_2_ = 10^−6^ μM^-1^ s^-1^, *k* = 10^−4^ s^-1^, and *θ* =10^7^ s.

**Fig. S17.**
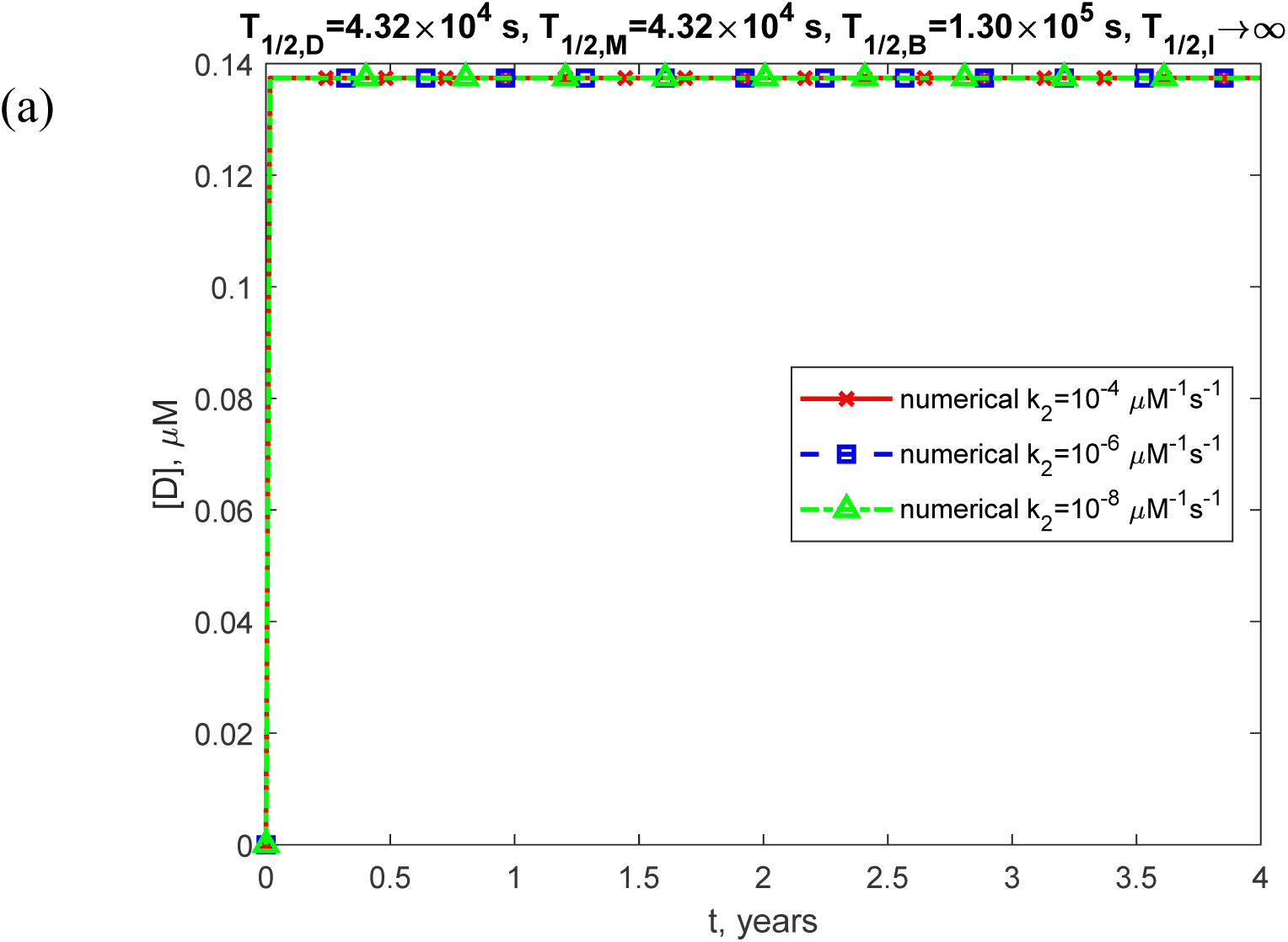
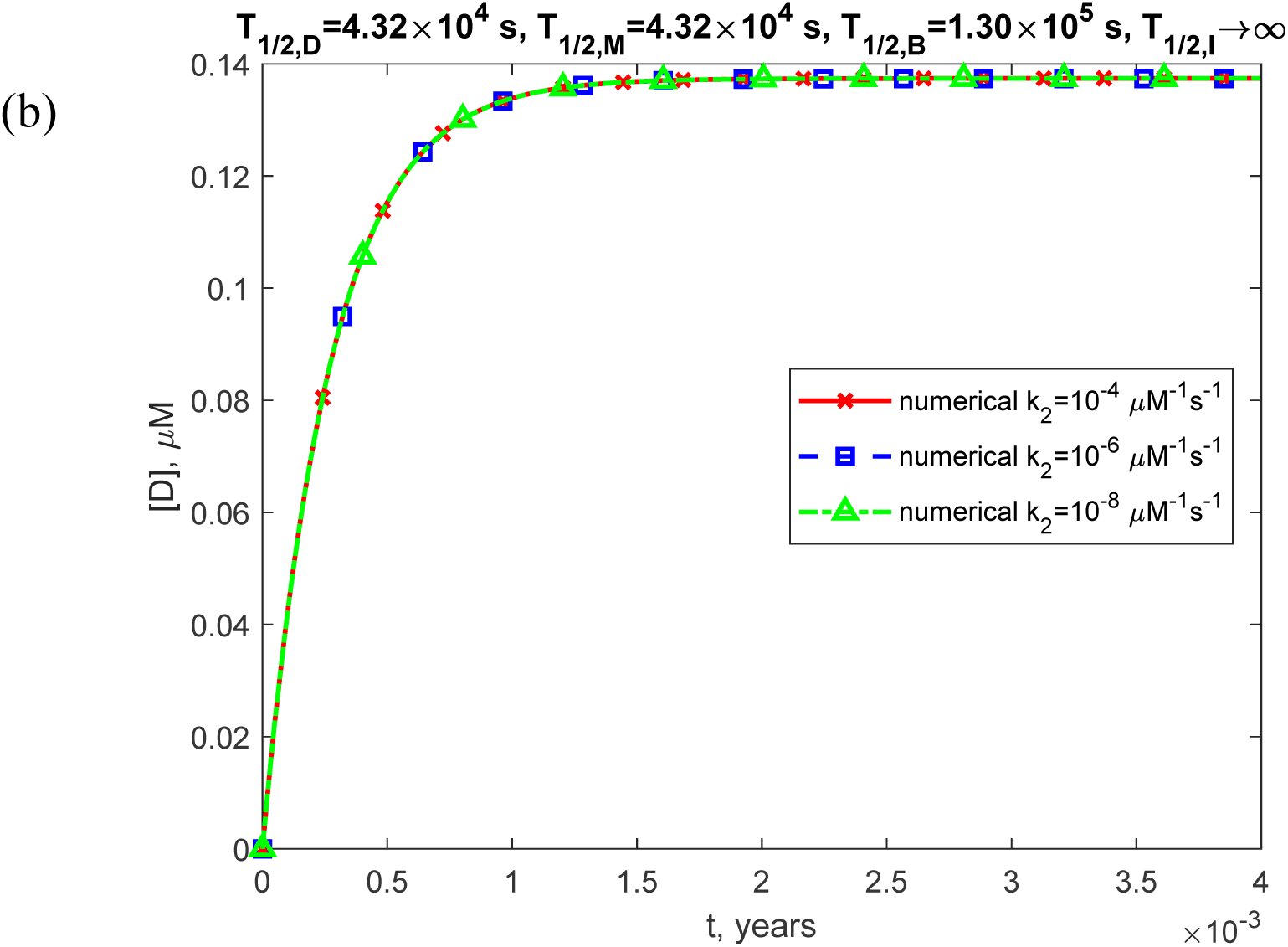
(a) Molar concentration of TDP-43 dimers, [*D*], as a function of time (in years). (b) Similar to Fig. S17a, but with a focus on the specific time range of [0, 0.004 years] on the *x*-axis. The numerical solution is obtained by solving Eqs. (4)-(7) with initial conditions (8). The presented scenario assumes a physiologically relevant half-life for TDP-43 dimers: *T* = 4.32×10^4^ s. The approximate solution is not shown, as it applies only to the scenario with infinite half-lives of TDP-43 dimers, monomers, and free misfolded oligomers. The results are displayed for three values of *k*_2_. The case with *k*= 10^−6^ s^-1^, *k*_1_ = 10^−4^ s^-1^, and *θ* =10^7^ s.

**Fig. S18.**
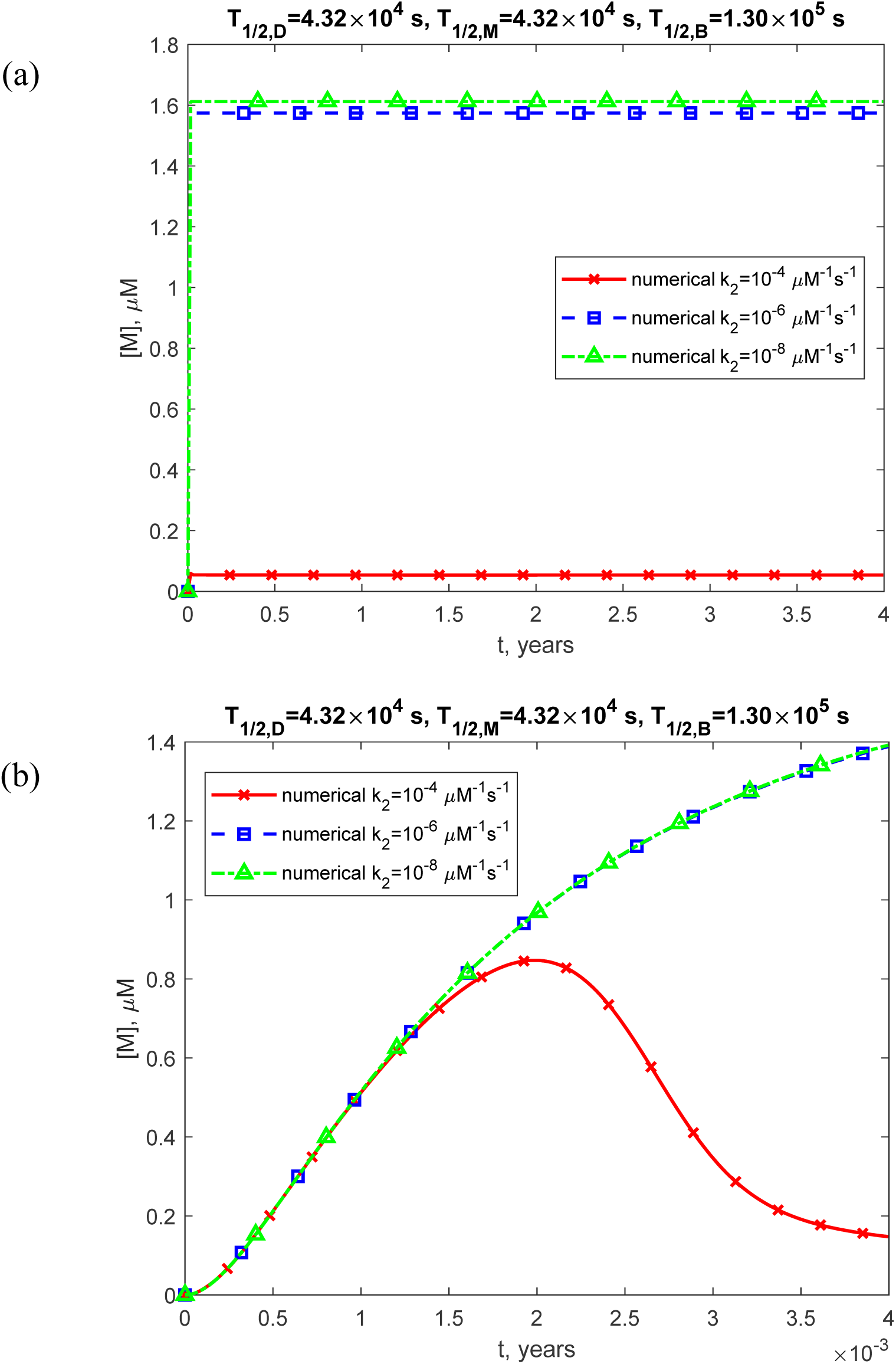
(a) Molar concentration of TDP-43 monomers, [*M*], as a function of time (in years). (b) Similar to Fig. S18a, but with a focus on the specific time range of [0, 0.004 years] on the *x*-axis. The numerical solution is obtained by solving Eqs. (4)-(7) with initial conditions (8). The presented scenario assumes physiologically relevant half-lives for TDP-43 dimers, monomers, and free misfolded oligomers: *T*_1/2,*D*_ = 4.32×10^4^ s, *T*_1/2,*M*_ = 4.32×10^4^ s, and *T*_1/2,*B*_ =1.30×10^5^s. The approximate solution is not shown, as it applies only to the scenario with infinite half-lives of TDP-43 dimers, monomers, and free misfolded oligomers. The results are displayed for three values of *k*_2_. The case with *k*_1_ = 10^−6^ s^-1^, *k_M_* = 10^−4^ s^-1^, and *θ* =10^7^ s.

**Fig. S19.**
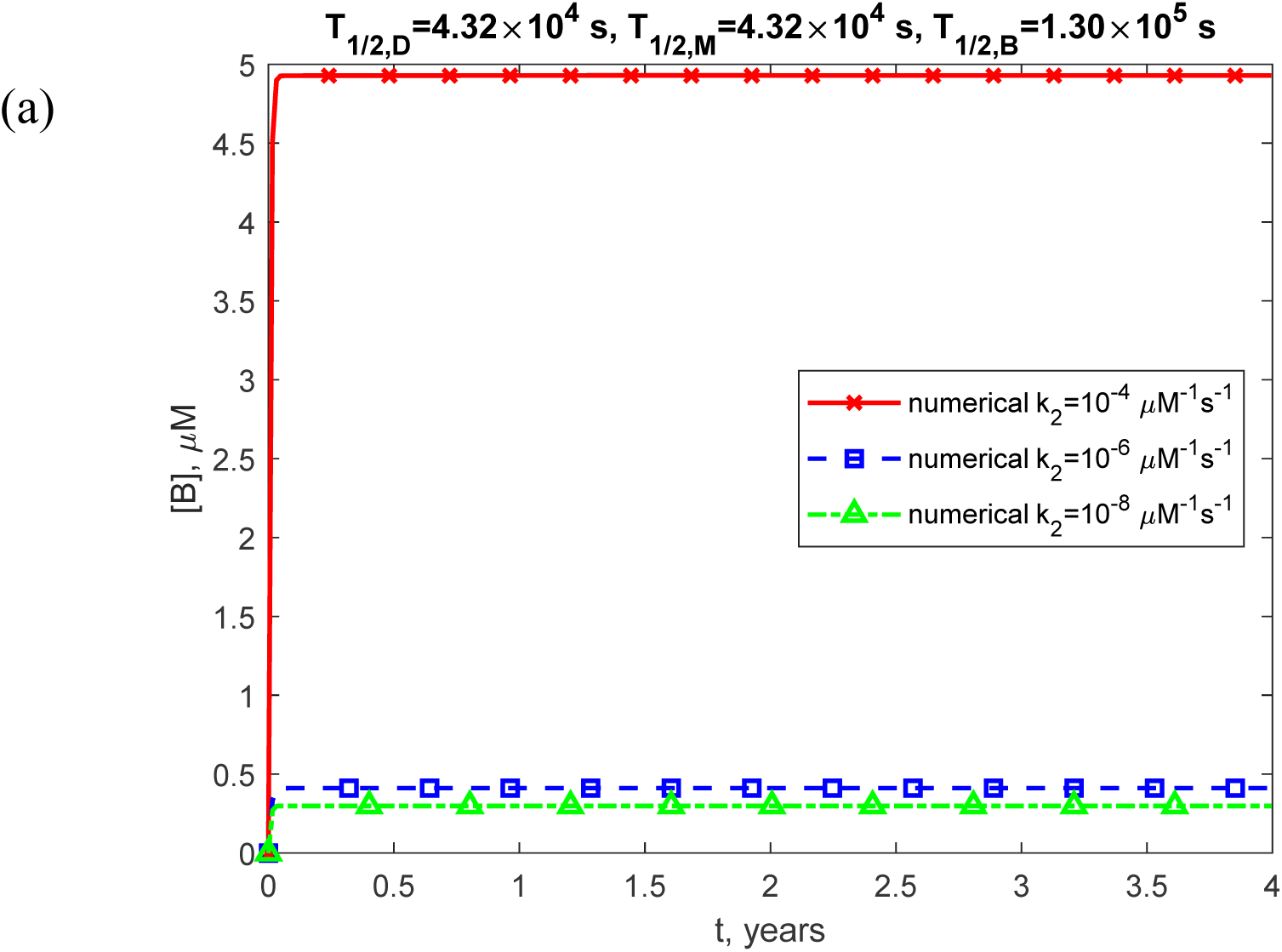
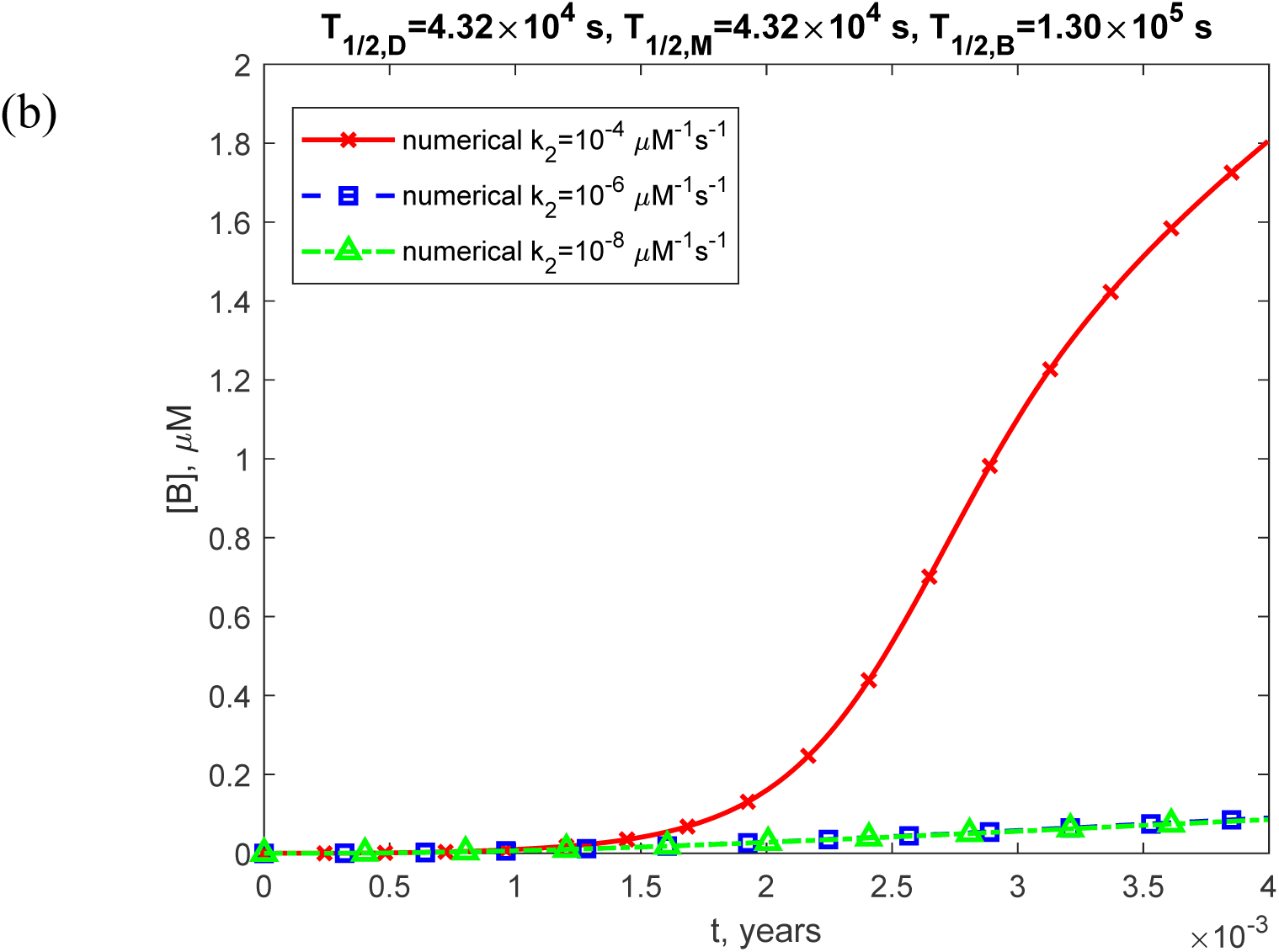
(a) Molar concentration of free misfolded TDP-43 oligomers, [*B*], as a function of time (in years). (b) Similar to Fig. S19a, but with a focus on the specific time range of [0, 0.004 years] on the *x*-axis. The numerical solution is obtained by solving Eqs. (4)-(7) with initial conditions (8). The presented scenario assumes physiologically relevant half-lives for TDP-43 dimers, monomers, and free misfolded oligomers: *T*_1/2,*D*_ = 4.32×10^4^ s, *T*_1/2,*M*_ = 4.32×10^4^ s, and *T*_1/2,*B*_ =1.30×10^5^ s. The approximate solution is not shown, as it applies only to the scenario with infinite half-lives of TDP-43 dimers, monomers, and free misfolded oligomers. The results are displayed for three values of *k*_2_. The case with *k*_1_ = 10^−6^ s^-1^, *k*= 10^−4^ s^-1^, and *θ* =10^7^ s.

**Fig. S20.**
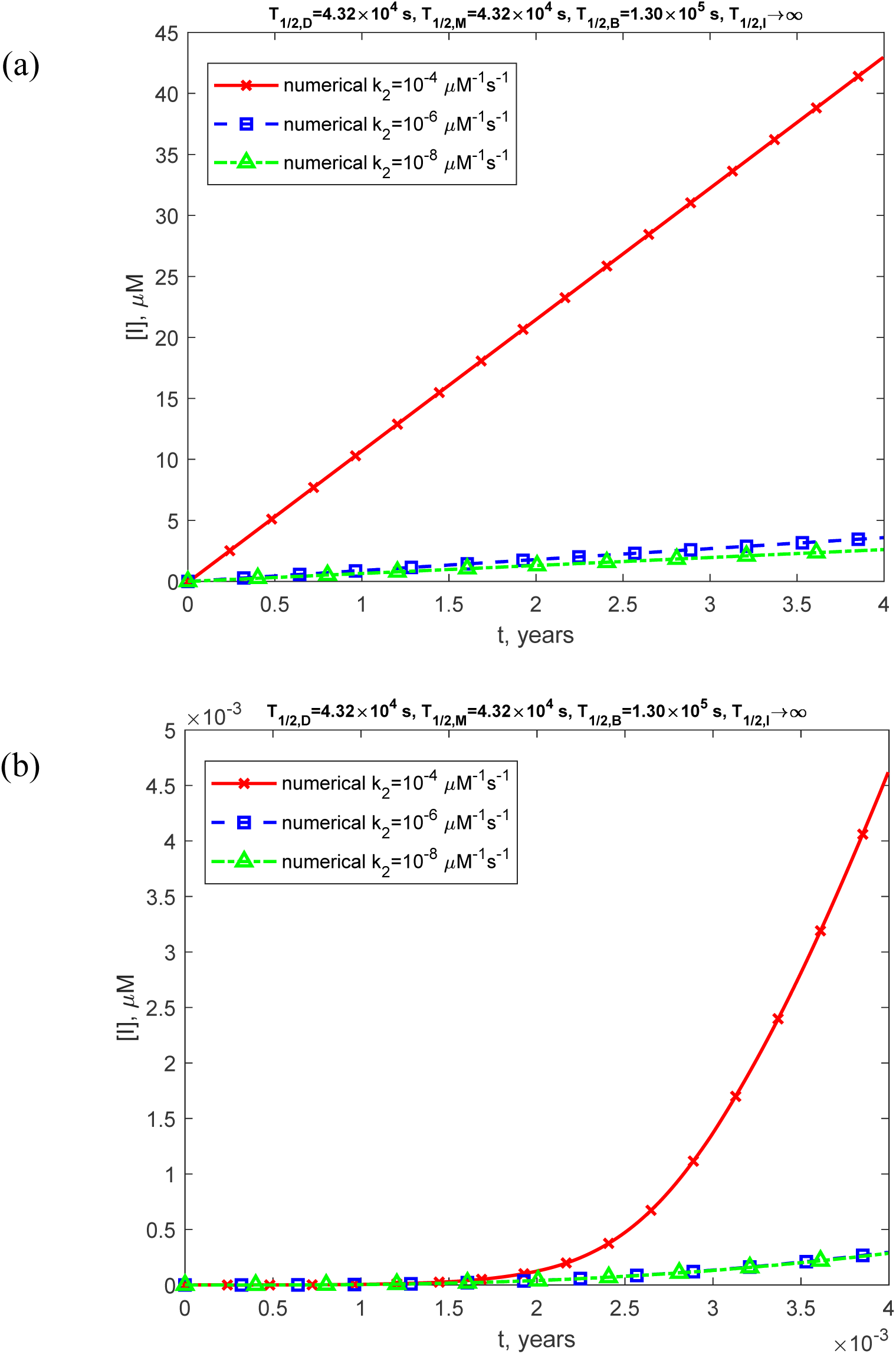
(a) Molar concentration of toxic TDP-43 oligomers incorporated into inclusion bodies, [*I*], as a function of time (in years). (b) Similar to Fig. S20a, but with a focus on the specific time range of [0, 0.004 years] on the *x*-axis. The numerical solution is obtained by solving Eqs. (4)-(7) with initial conditions (8). The presented scenario assumes physiologically relevant half-lives for TDP-43 dimers, monomers, free misfolded oligomers, and oligomers deposited into inclusion bodies: *T*_1/2,*M*_ = 4.32×10^4^ s, *T*_1/2,*B*_ = 4.32×10^4^ s, *T* =1.30×10^5^ s, and *T*_1/2,*I*_ →∞. The approximate solution is not shown, as it applies only to the scenario with infinite half-lives of TDP-43 dimers, monomers, and free misfolded oligomers. The results are displayed for three values of *k*_2_. The case with *k*_1_ = 10^−6^ s^-1^, *k_M_* = 10^−4^ s^-1^, and *θ* =10^7^ s.

**Fig. S21.**
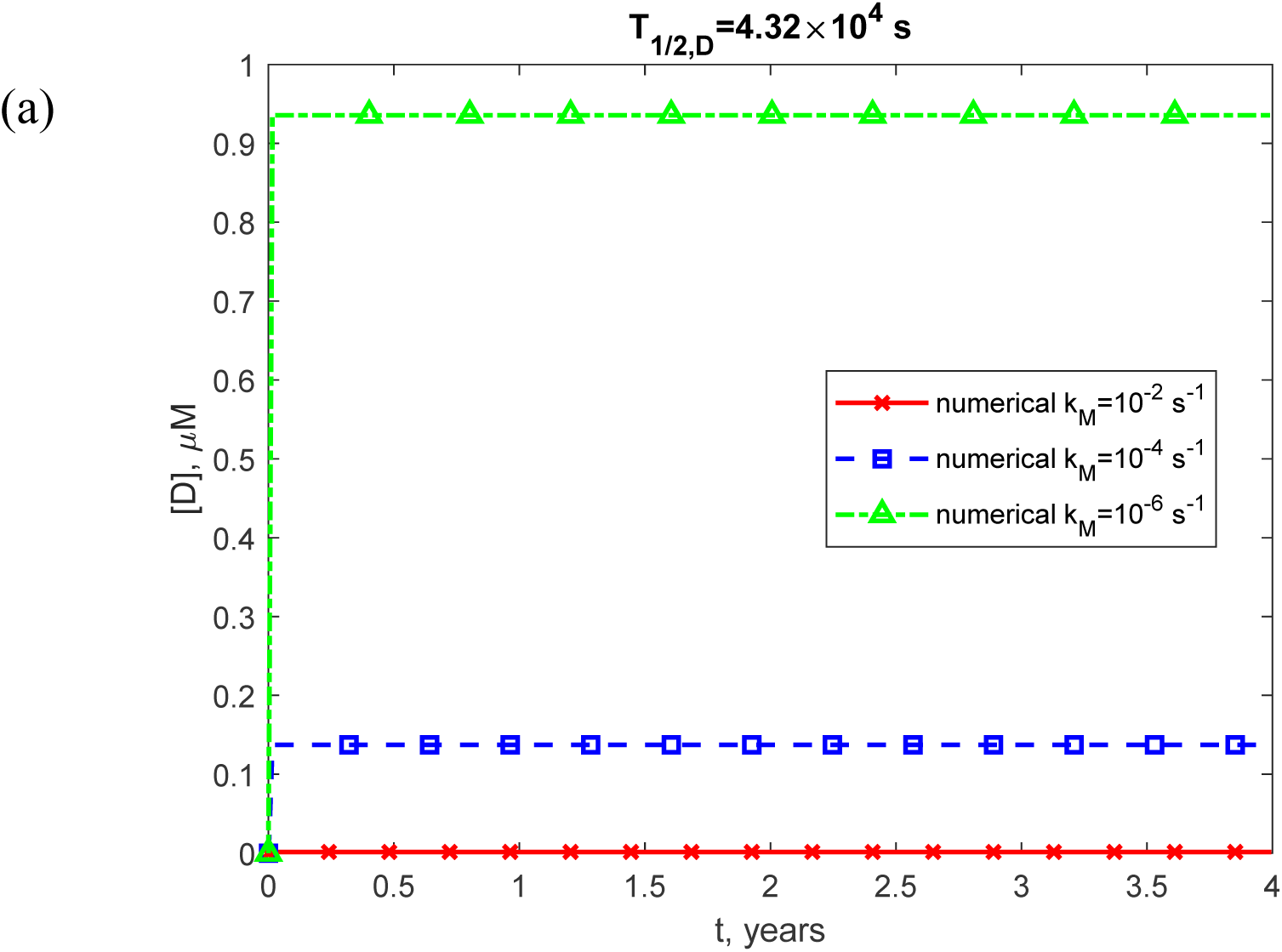
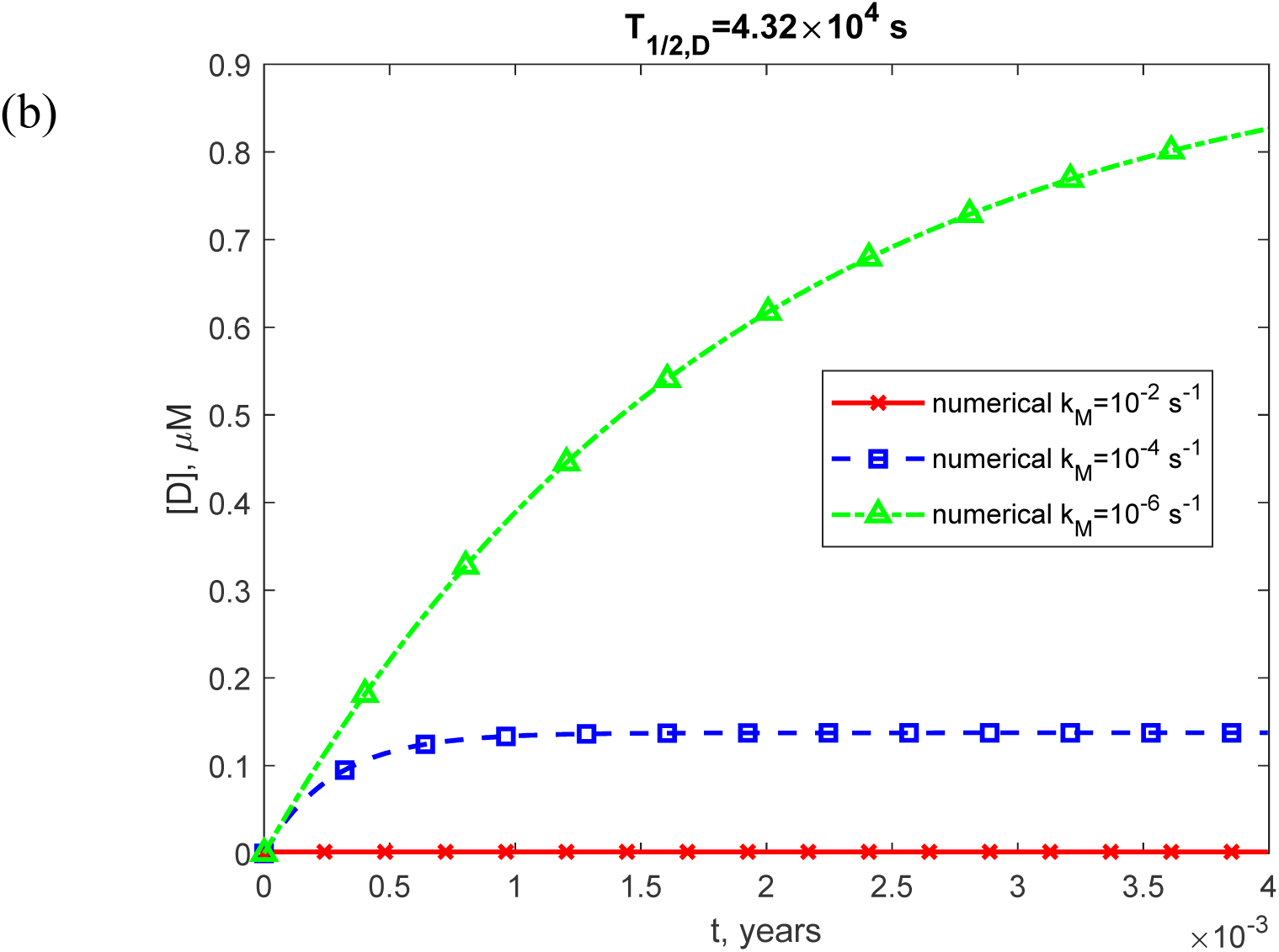
(a) Molar concentration of TDP-43 dimers, [*D*], as a function of time (in years). (b) Similar to Fig. S21a, but with a focus on the specific time range of [0, 0.004 years] on the *x*-axis. The numerical solution is obtained by solving Eqs. (4)-(7) with initial conditions (8). The presented scenario assumes a physiologically relevant half-life for TDP-43 dimers: *T* = 4.32×10^4^ s. The approximate solution is not shown, as it applies only to the scenario with infinite half-lives of TDP-43 dimers, monomers, and free misfolded oligomers. The results are displayed for three values of *k_M_*. The case with *k*_1_ = 10^−6^ s^-1^, *k* = 10^−6^ μM^-1^ s^-1^, and *θ* =10^7^ s.

**Fig. S22.**
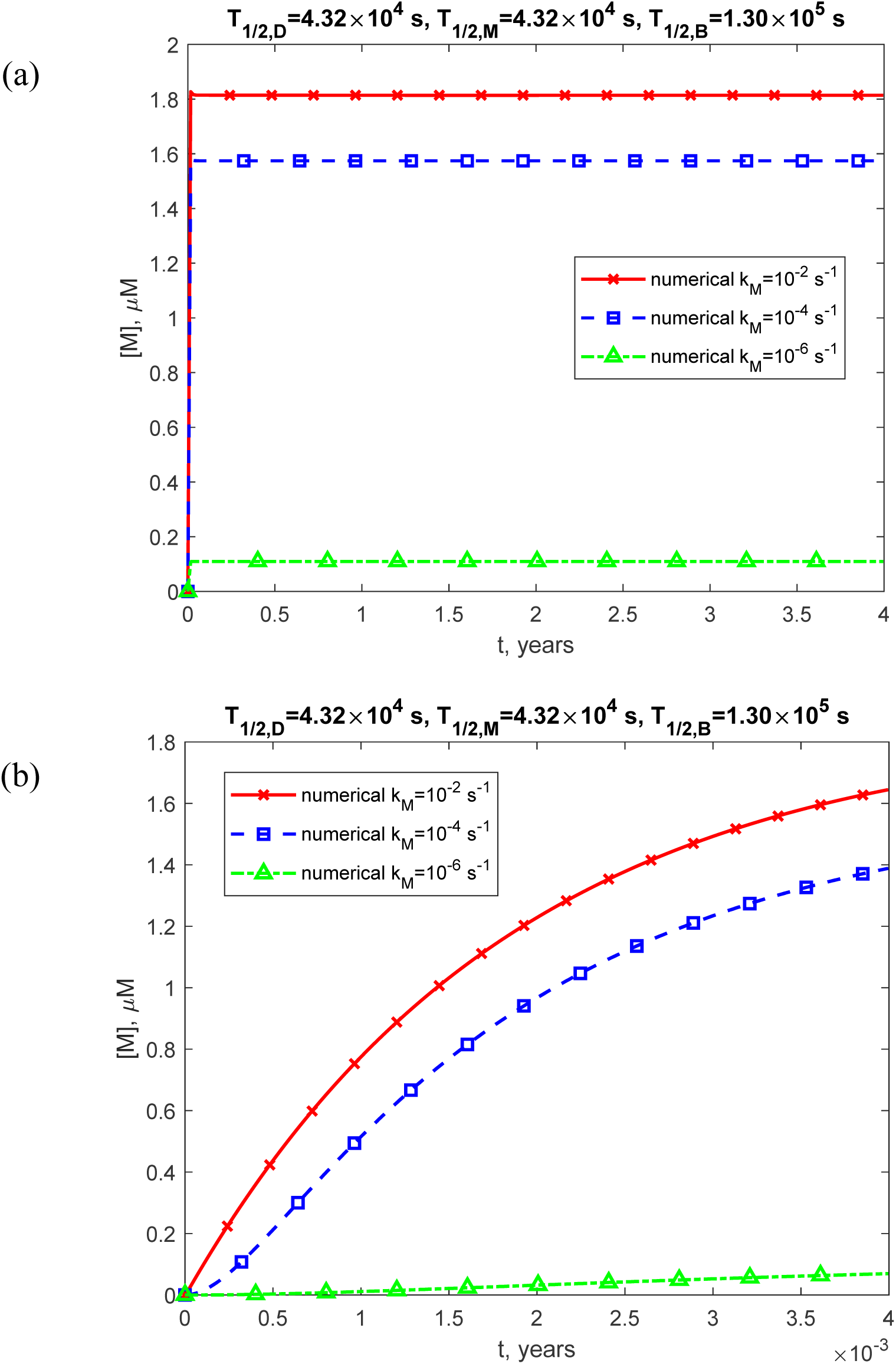
(a) Molar concentration of TDP-43 monomers, [*M*], as a function of time (in years). (b) Similar to Fig. S22a, but with a focus on the specific time range of [0, 0.004 years] on the *x*-axis. The numerical solution is obtained by solving Eqs. (4)-(7) with initial conditions (8). The presented scenario assumes physiologically relevant half-lives for TDP-43 dimers, monomers, and free misfolded oligomers: *T*_1/2,*D*_ = 4.32×10^4^ s, *T*_1/2,*M*_ = 4.32×10^4^ s, and *T*_1/2,*B*_ =1.30×10^5^s. The approximate solution is not shown, as it applies only to the scenario with infinite half-lives of TDP-43 dimers, monomers, and free misfolded oligomers. The results are displayed for three values of *k_M_*. The case with *k_1_* = 10^−6^ s^-1^, *k_2_* = 10^−6^ μM^-1^ s^-1^, and *θ* =10^7^ s.

**Fig. S23.**
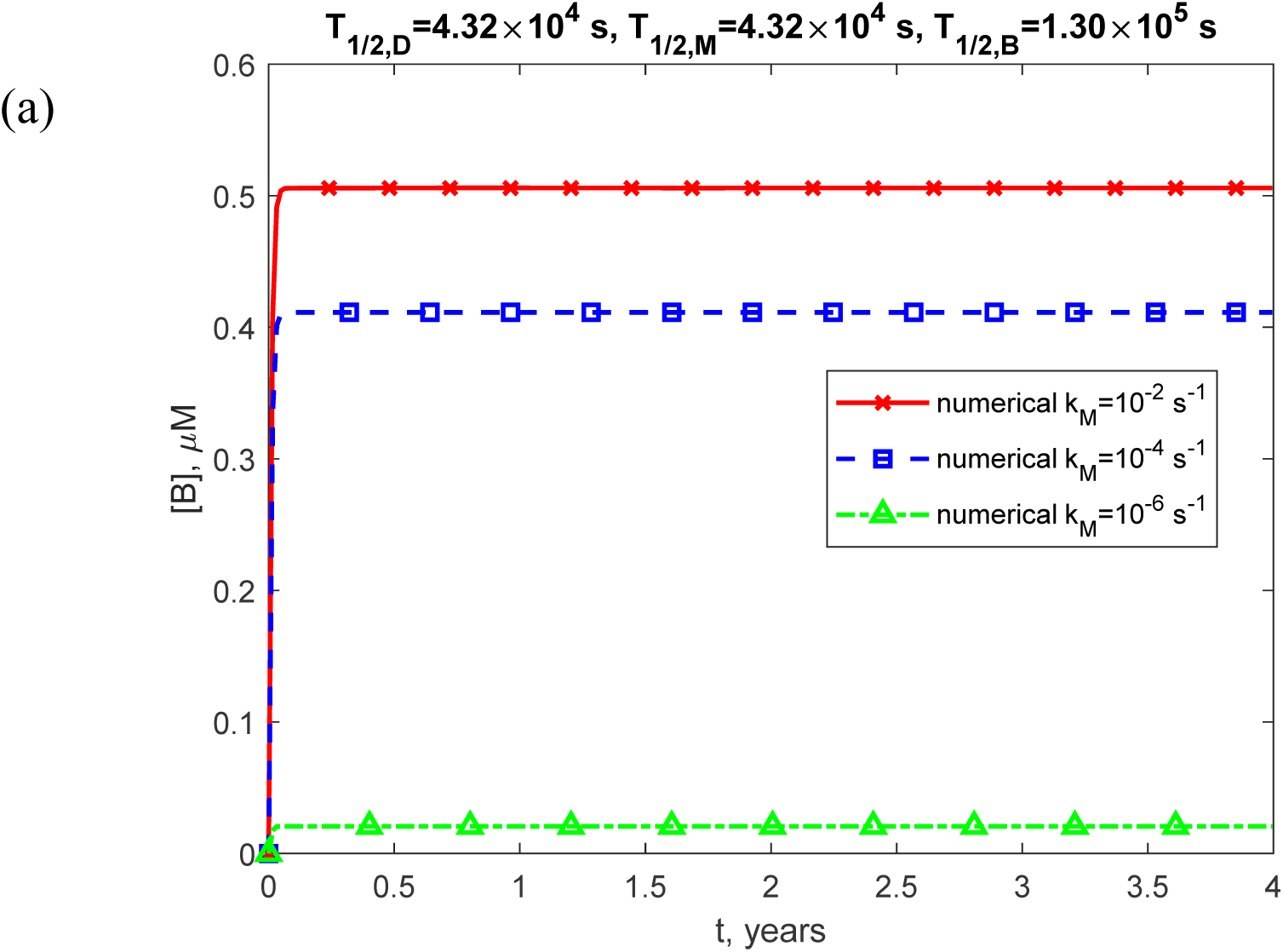
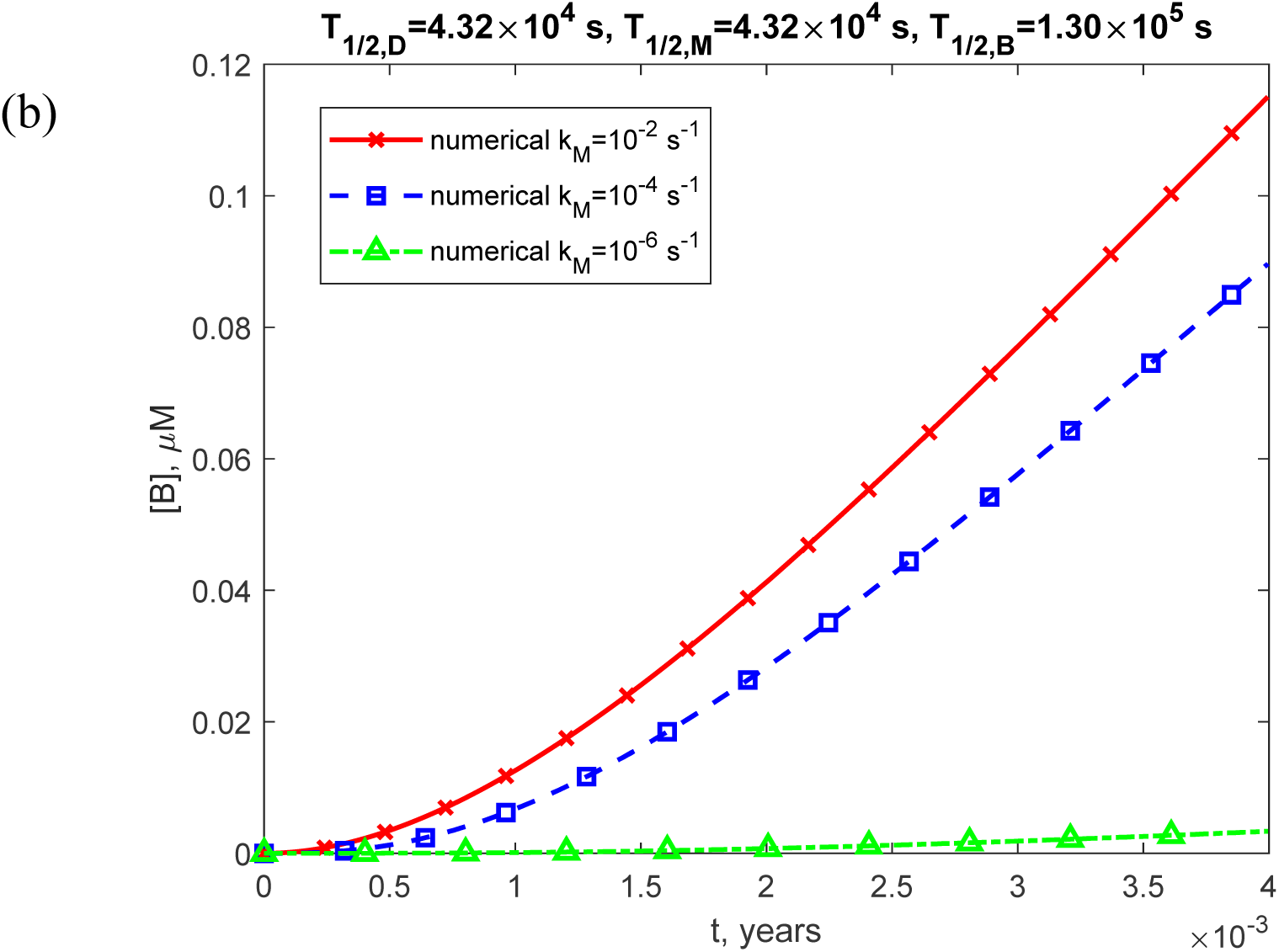
(a) Molar concentration of free misfolded TDP-43 oligomers, [*B*], as a function of time (in years). (b) Similar to Fig. S23a, but with a focus on the specific time range of [0, 0.004 years] on the *x*-axis. The numerical solution is obtained by solving Eqs. (4)-(7) with initial conditions (8). The presented scenario assumes physiologically relevant half-lives for TDP-43 dimers, monomers, and free misfolded oligomers: *T*_1/2,*D*_ = 4.32×10^4^ s, *T*_1/2,*M*_ = 4.32×10^4^ s, and *T*_1/2,*B*_ =1.30×10^5^ s. The approximate solution is not shown, as it applies only to the scenario with infinite half-lives of TDP-43 dimers, monomers, and free misfolded oligomers. The results are displayed for three values of *k_M_*. The case with *k*_1_ = 10^−6^ s^-1^, *k* = 10^−6^ μM^-1^ s^-1^, and *θ* =10^7^ s.

**Fig. S24.**
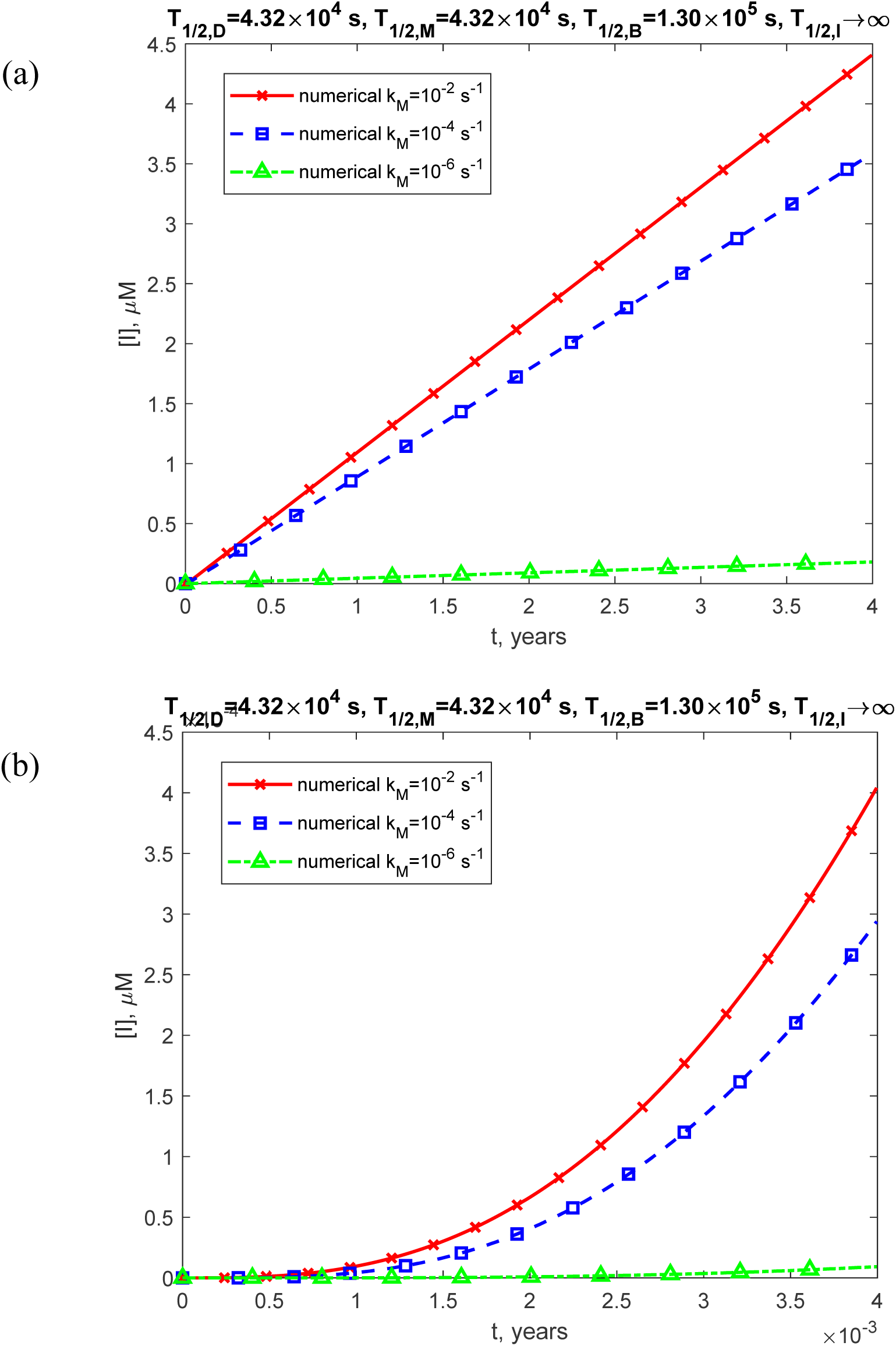
(a) Molar concentration of toxic TDP-43 oligomers incorporated into inclusion bodies, [*I*], as a function of time (in years). (b) Similar to Fig. S24a, but with a focus on the specific time range of [0, 0.004 years] on the *x*-axis. The numerical solution is obtained by solving Eqs. (4)–(7) numerically with initial conditions (8). The presented scenario assumes physiologically relevant half-lives for TDP-43 dimers, monomers, free misfolded oligomers, and oligomers deposited into inclusion bodies: *T*_1/2,*D*_ = 4.32×10^4^ s, *T*_1/2,*M*_ = 4.32×10^4^ s, *T*_1/2,*B*_ =1.30×10^5^ s, and *T*_1/2,*I*_ →∞. The approximate solution is not shown, as it applies only to the scenario with infinite half-lives of TDP-43 dimers, monomers, and free misfolded oligomers. The results are displayed for three values of *k_M_*. The case with *k*_1_ = 10^−6^ s^-1^, *k* = 10^−6^ μM^-1^ s^-1^, and *θ* =10^7^ s.

